# Functional and intricate interaction network connecting *Helicobacter pylori* Cag Type 4 Secretion System surface proteins with outer membrane proteins HopQ and HopZ

**DOI:** 10.1101/2025.09.01.673027

**Authors:** Felix Metz, Johanna Beilmann, Simon H. Bats, Andreas Latoscha, Gregor Witte, Remco T. A. Megens, Karl-Peter Hopfner, Kaisa Thorell, Wolfgang Fischer, Laurent Terradot, Sebastian Suerbaum, Christine Josenhans

## Abstract

The *Helicobacter pylori cag* pathogenicity island (*cag*PAI) encodes a complex type IV secretion system (CagT4SS) which is an important virulence factor of *H. pylori*. Recently, structural detail on the CagT4SS has been substantially improved by cryo-EM. However, important structural and functional information, in particular on protein interactions between T4SS surface proteins, and of T4SS surface proteins with other proteins, is missing. In the present study, we followed the hypothesis that *H. pylori* T4SS external proteins may form a surface protein assembly, together with other, non-CagT4SS proteins, which may also be essential for T4SS function. Using interaction screens of *H. pylori* CagT4SS surface proteins, followed by biochemical and functional characterization, we have enhanced the knowledge on protein-protein interactions of CagT4SS extracellular proteins. This also includes newly identified interactions of CagT4SS surface proteins, for instance the VirB2 homolog CagC, the VirB5 homolog CagL and the surface protein CagN of unknown function, with outer membrane proteins HopQ and HopZ. We have further identified and quantitated direct interactions of T4SS surface proteins with outer membrane proteins HopZ and HopQ, which play a role in T4SS functions, and of both HopZ and HopQ with themselves and with host cell factors CEACAM and integrin. Furthermore, we determined an influence of pH on interactions between HopQ/HopZ and CagT4SS components. Utilizing protein tag insertions in *H. pylori*, we detected surface-exposed association of HopQ and HopZ with T4SS components on bacteria without or with (for HopQ) human gastric epithelial cells. Functionally antagonistic roles of HopQ and HopZ were uncovered in T4SS-dependent early pro-inflammatory human epithelial cell activation. In summary, we identified a network of interactions between *H. pylori* outer membrane proteins and CagT4SS surface proteins and characterized them as functionally important for transport processes. This will help to refine structural and functional details regarding surface-exposed proteins of the CagT4SS.

## Introduction

*Helicobacter pylori* is a Gram-negative human stomach pathogen, which is strongly linked to the causation of several gastric diseases in humans (1, 2). *H. pylori* infection is primarily acquired during early childhood via fecal-oral or oral-oral transmission and affects approximately half of the world population (3–6). The consequences of *H. pylori* infections depend on a number of environmental, bacterial and host specific factors (3), and most infected persons never develop overt disease. Nonetheless, about 15% of affected patients will develop serious long-term health consequences. Not only do studies show a link to the development of gastric cancer (7, 8) but also the enhanced risk of severe gastritis, peptic ulcer disease, gastric cancer, and MALT lymphomas (6).

*H. pylori* has co-evolved with its human host for at least 100,000 years, into different ethnic and further admixed bacterial populations (9, 10). At some point in time, it has acquired, from an unknown source, one of its major immune modulators, the Cag type IV secretion system (CagT4SS), a complex bacterial membrane transport system (11–15). The individual virulence of *H. pylori* strains and severity of disease caused are closely related to the presence of the CagT4SS (8, 10, 16). Together with its main effector protein CagA, the T4SS is encoded on the *cag* pathogenicity island (*cag*PAI ((11, 17)), present in about 70% of strains and highly genetically variable between strains (10). The *H. pylori cag*PAI comprises about 28 genes, roughly 13 of which are coding for homologs of the *Agrobacterium tumefaciens* Vir T4SS model system (18). The remaining genes code for additional *H. pylori*-specific T4SS components (16, 19–21) and also comprise genes of unknown function (22).

A high extent of genetic variability is found between different T4SS (reviewed in (18, 23, 24)), whose paradigm is the VirB/D system of the plant pathogen *A. tumefaciens* (25). More complex Type 4 systems are, for instance, the Dot/Icm system (26) of *Legionella pneumophila* and the *H. pylori* CagT4SS (27, 20, 28). The precise architecture of the various T4SS protein assemblies and machineries is not fully understood yet, despite a growing number of excellent high-resolution Cryo-EM (18, 21, 28–30), and protein-protein interaction studies (31, 32). As those findings show, the core structure of the *H. pylori* CagT4SS associated with the bacterial outer membrane (outer membrane core complex, OMCC), is composed of mainly five proteins (CagY, CagX, CagT, CagM and Cag3), all present in multiple copies (30, 33). The CagT4SS OMCC (OMCC_Cag_)(18, 33) is localized between the bacterial inner and the outer membranes, and CagY spans both the inner and outer membranes (33) and is therefore not restricted to the OMCC_Cag_. As far as it has already been visualized, the CagT4SS inner membrane complex (IMC) consists of three ring structures around a central channel, which are, among other proteins, formed by CagY, and contains the ATPases Cagβ, CagE, and Cagα at the inner membrane (20, 28).

In addition to the structurally important, membrane associated, IMC and OMC(C), subassemblies of the *H. pylori* T4SS (30, 31, 34), some Cag proteins seem not to possess a structural role so far (22, 35–37). We and others previously characterized a CagT4SS protein, CagN, of unknown function (38), which is partially surface-associated in the CagT4SS (21) and interacts with CagM, a component of the OMCC_Cag_ (24, 33). The proteins CagL, CagC, CagH and CagI were reported before to be surface-associated in the bacteria (35, 36, 39). Of those, CagL, CagI and CagH are already known to interact with each other (36, 40). Each of those proteins influence the expression as well as the stability of the respective other proteins and are essential for the formation of a connecting structure that was detected in scanning electron microscopy (EM) on the surface of *H. pylori* in contact with human gastric epithelial cells (37, 40, 41). Although CagC alone appeared not to be strictly required for the formation of such a structure (39), CagC was also located on the surface of *H. pylori* by EM and fluorescence microscopy, and, similar to the other CagT4SS surface proteins, is essential for the translocation of CagA and the induction of pro-inflammatory IL-8 secretion in human host cells (35–37, 42, 43).

The Cag proteins CagL and CagI, the oncogenic effector CagA, and the OMCC protein CagY interact directly with host cell integrin receptors, for instance with α_5_β_1_ integrin (37, 44–46). While the integrin interaction seems to be dispensable for the so far known T4SS functions, the interaction of the *H. pylori* outer membrane autotransporter protein HopQ with human cell surface receptors <u>c</u>arcino<u>e</u>mbryonic <u>a</u>ntigen-related <u>c</u>ell <u>a</u>dhesion <u>m</u>olecules (CEACAMs) was found to be important for CagA transport into human cells (15, 47). HopQ is one member of a large family (> 30 members) of outer membrane proteins (OMPs) in *H. pylori*, making up roughly 4% of the *H. pylori* genome (48). The OMP family also includes *H. pylori*’s main adhesins BabA and SabA (48–50). The HopQ proteins are subdivided into two closely related but sequence-divergent allelic families, HopQI and HopQII (51). While the majority of *H. pylori* strains carries only one HopQ allele, of which HopQI is more common, some strains carry both HopQ alleles, HopQI and HopQII (52). The HopQI genotype is significantly linked to a *cagPAI*- and therefore CagT4SS-positive status, as well as to the presence of the *type* s1-m1 VacA (which has higher vacuolating activity and disease association than other VacA types (53, 54)) and is dominant in Asian strains (47, 55, 56). The interaction of both HopQ alleles, HopQI and HopQII with human CEACAM was studied previously in some detail (56, 57). This interaction, which can also dissolve CEACAM dimers on the host cell side, has been shown to mediate both the induction of pro-inflammatory signaling and translocation of CagA into the human host cell (47, 58, 59).

A highly sequence-related paralog of the same family of *H. pylori* OMPs is HopZ (60). Like HopQ, it presents as two allelic genotypes, *hopZI* and *hopZII,* which are usually alternatively present in *H. pylori*. The two *hopZ* alleles belong to the most strain-variable outer membrane proteins of *H. pylori*, indicating that they are under strong diversifying selection (60, 61). While large parts of the N- and C-terminal regions of the HopZ amino acid sequence, which constitute the common beta-barrel structure of the OMPs embedded in the outer membrane (48, 56), are highly conserved between different strains and between different OMPs, especially the predicted surface-exposed segments in the sequence of the mature outer membrane proteins are rather variable between strains, although they belong recognizably to one of the two HopZ types (60). The most notable difference between the protein sequences of the two HopZ alleles is a 20 amino acid stretch of HopZI (aa179 to aa198) that is missing in HopZII (61). Other *H. pylori* OMPs are much less strain variable (56).

Potential host cell receptor(s) of HopZ are unknown, and conflicting results were reported regarding a role of HopZ in human cell adhesion (60, 61). Interestingly, the *hopZ* coding region carries a phase-variable CT dinucleotide-repeat in its 5’ coding mRNA region that may function as an ON/OFF switch for protein expression (60, 61). No clear correlation between *hopZ* status, *cag*PAI and clinical disease was previously identified. While, in contrast to HopQ, no statistically significant correlation between *hopZ* allele and *cag*PAI-positivity was reported so far, however a trend for *hopZI* to genomically co-occur with *cag*PAI-positivity, and a significant correlation for *hopZI* with the more active *vacAs1m1*, *cag*PAI associated, genotype of the VacA toxin exists (60). *hopZ* (both alleles) was found frequently in CT repeat-OFF status during chronic infection, but underwent a CT-ON-switch in one vaccine study upon acute infection of healthy volunteers (62) We hypothesized from the available evidence that further *H. pylori* Cag surface proteins, that we also term “Cag outer proteins”, can interact with each other in a more complex network. A possible protein assembly at the outer surface of the CagT4SS machinery may have a contact and conduit function towards human host cells. In addition, we assumed that a functional and a direct physical interaction of HopQ, for which a relevance for CagT4SS functionalities was shown before, and HopZ may exist with CagT4SS proteins. Therefore, the initial aim of this study was to determine and characterize so far unknown/novel protein-protein interactions of the *H. pylori* CagT4SS surface proteins, with the goal to better understand the potential interaction network, especially of a proposed Cag surface protein assembly. Bacterial Two-Hybrid (BACTH) assays were used to perform an interactome screen for potential novel interaction partners in- and outside of the T4SS, in particular with respect to Cag surface proteins and the two *H. pylori* outer membrane proteins HopQ and HopZ. Thereby, we have identified novel interactions at the proposed surface assembly of the CagT4SS, and we obtained further evidence for direct physical binding of Cag proteins with the two outer membrane proteins HopQ and HopZ. The interaction network that results of the present study, jointly with functional changes that we obtained with bacterial mutants, suggests new characteristics of the CagT4SS and a novel functional role of HopZ with respect to the CagT4SS.

## Results

### Bacterial Two-Hybrid (BACTH) assay reveals novel interactions of outer CagT4SS proteins in a partially unbiased screen

In order to screen, in a partially unbiased manner, regarding the selected combinations of protein expression constructs in the test array, for protein-protein interactions between members of the outer proteins of the *H. pylori* T4SS, bacterial two hybrid assays were performed to detect novel binary interactions. For this purpose, we used expression constructs for proteins of the CagT4SS OMCC and Cag surface proteins. The existing BACTH set-up, which we had designed before for CagM interaction analysis (22) was expanded by 39 new plasmids (different insertions in the four available BACTH vectors; see Methods) encoding the proteins CagH, CagI, CagL, CagC, CagA, HopQI (S1 Fig), HopQII (S1 Fig), HopZI (S1 Fig), HopZII (S1 Fig) and human CEACAM1.

After screening 327 new plasmid combinations, in comparison with common negative and positive controls (21) (S2 Table, Fig 1A, and Methods), we identified 8 novel CagT4SS protein-protein interactions not previously known. All obtained β-galactosidase results higher than 1.5-fold of the respective negative control values were scored as positive (Methods) and are included in the all-results matrix (Fig 1A) as fold-changes over the negative control of the respective assay.

**Fig 1.**
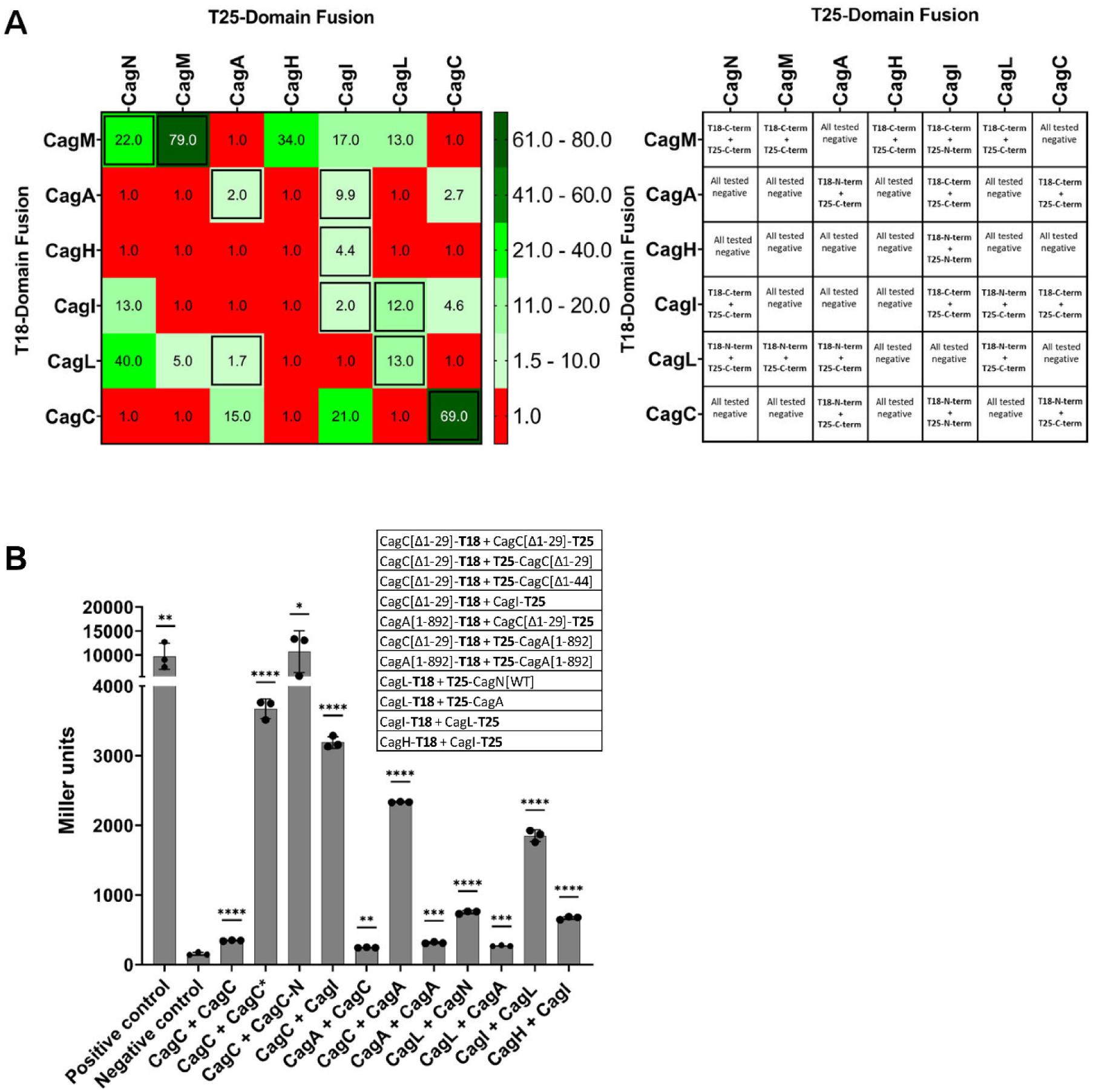
Bacterial-Two-Hybrid-Assay uncovers novel interactions between proteins of the *H. pylori* CagT4SS surface proteins. **A)** intensity matrix of unbiased BACTH results (β-galactosidase assays, determined as Miller units) to detect and quantitate novel protein-protein interactions between the CagT4SS outer proteins. Red color indicates no detectable interaction above the negative control (ratio of 1 to the negative control), green shades of color indicate detectable interactions (significantly above background levels; detection limits were defined as below 1.5-fold the average of the negative control background over all experiments (Methods)); the values in the matrix boxes correspond to the ratio between the Miller unit results of each combination measured over the negative control (fold change); for all tested interactions with positive outcome, exclusively the BACTH plasmid combination for each interaction pair with the highest quantitative outcome of interaction is included in the matrix. If all assayed interactions were negative, one negative result is shown. Negative and positive controls were run alongside each separate assay and are explained in the Methods. Previously reported interactions of CagT4SS surface proteins (33, 36, 40, 67) are boxed. Tabular depiction of the precise plasmid combinations used in the matrix is shown to the right. Additional assay results and Western Blots are shown in S2 Fig). **B)** BACTH results of β-galactosidase activity of selected new interactions between CagT4SS outer proteins, depicted as bar graphs in absolute values (Miller units). All assays were performed and quantitated in three-fold replicates. The first listed interaction partner on the x-axis in each combination was expressed as an N-terminal fusion in the pUT18 BACTH plasmid; the second listed interaction partner in each pair was expressed in pKT25 or pKNT25 (see also legend table and Methods). The legend table included in panel B contains the full information about the BACTH plasmid fusions and combinations used in each assay. Statistical analysis for significant difference was performed for the comparison between each sample versus the negative control (Student’s *t*-test, two-tailed unpaired), and significant p values are shown above each sample. *p<0.05; **p<0.01; ***p<0.001; ****p<0.0001. CagC was fused as a shorter variant missing a putative leader peptide sequence (Δaa1-29); CagC* is a fusion of the same N-terminally truncated protein product which is expressed as a C-terminal fusion in BACTH plasmid pKT25. CagC-N designates an N-terminally truncated version of CagC (Δaa1-44) and was cloned as a C-terminal fusion in pKT25. CagA was expressed as an N-terminal fusion of a C-terminally truncated variant, only comprising the N-terminal domain consisting of aa1-892 in either pUT18 or in pKT25. CagA full-length was also cloned and tested, however was poorly expressed and always gave negative results in BACTH. All used expression constructs including CagL, CagI and CagH are described in S2 Table. Selected Western blots (e.g. CagC*) of the BACTH samples, which demonstrate protein expression of fusion proteins in *E. coli*, are shown in the supplementary figures (S2 Fig). Detailed BACTH analyses of CagM interactions with Cag surface proteins are shown in the supplementary figures (S2 Fig).

Within the CagT4SS protein interaction network itself, and with a focus on the Cag proteins that might be located at the surface of the T4SS, the novel set of interactions revealed included, among others, the self-interactions of CagC and CagA, as well as the heterologous interactions of CagC with CagA, CagL and CagI with CagM, CagI with CagH, and CagN with both CagL and CagI, respectively (Fig 1A, 1B; S2A, S2B, S2C, S2F Fig). Some known direct interactions of CagT4SS surface proteins, such as those between CagL and CagI, CagA and CagL, CagA and CagI, determined previously by biophysical methods (46), were confirmed. Significantly positive, partially novel, BACTH interactions of the translocated effector CagA with CagC (VirB2 homolog) or the CagL (VirB5 homolog), as well as CagC-CagC and CagA- CagA self-interactions, are separately depicted as β-galactosidase assays (Fig 1B).

### **BACTH screen reveals novel interactions between proposed CagT4SS surface proteins and** *H. pylori* outer membrane proteins HopQ and HopZ

Owing to the previous characterization of a direct functional interaction between HopQ and host CEACAM proteins, strongly influencing CagT4SS functionality, including CagA transport and inflammatory processes by various transported molecules in the target human cells (47, 56), we also included both HopQI and HopQII (expressed without the transmembrane β-barrel) in our BACTH screen. We also hypothesized that both HopZI and HopZII alleles, due to the fact that HopZ is a closely related paralog with substantial amino acid sequence and structural homology to HopQ (S1 Fig), are additional candidates for being potential direct interaction partners of components of the *H. pylori* CagT4SS. Hence, HopZ, in its two allelic types, was also included in our present BACTH test arrays in *E. coli*. Thereby, we found, in addition to the 8 novel Cag-Cag interactions (above), that both alleles of HopQ as well as HopZ exhibited multiple additional interactions in this assay (Fig 2). HopQ and HopZ showed homotypic and heterotypic self-interactions (except for HopZII with HopQI) (Fig 2A, 2B). In addition, the introduction of HopQ and HopZ as potential new interaction partners of the T4SS proteins revealed 10 novel Hop interactions with specific *H. pylori* CagT4SS proteins (Fig 2G; S2G Fig). While both Hop proteins showed positive interactions with CagA (Fig 2A, C) and CagL (Fig 2A, D) in various BACTH combinations, assays involving either of both HopZ alleles also revealed additional interactions with CagI, CagC, and CagN (Fig 2A, 2E; S2D, S2E Fig).

**Fig 2.**
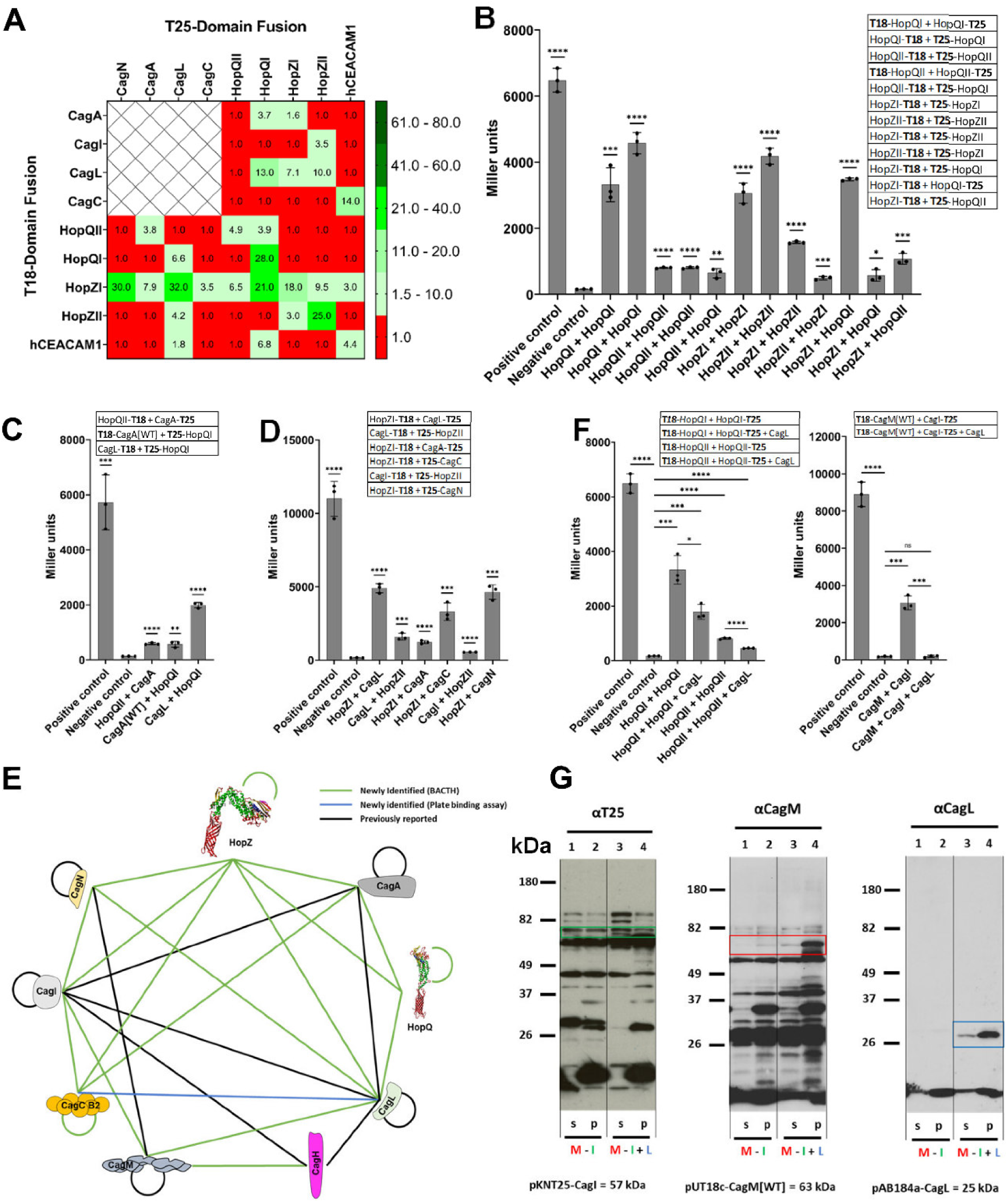
BACTH and newly devised Bacterial-Three-Hybrid-Assay (BAC3H) characterize novel direct interactions between HopQ and HopZ proteins, the. *H. pylori* **CagT4SS surface proteins, and between human CEACAM1, CagT4SS proteins and HopQ/HopZ, predicting an entirely novel interaction network and competitive interactions. A)** intensity matrix of BACTH results (β-galactosidase assays, quantitated in Miller units) to quantitate selected protein-protein interactions between various proteins of the CagT4SS outer proteins with outer membrane proteins HopQ and HopZ. Red color indicates no detectable interaction above the negative control (ratio of 1 over the negative control), green shades of color indicate detectable interactions; the values in the matrix boxes correspond to the ratios between the Miller unit results measured for each new interaction and that of the negative control (fold change). Positive controls with leucine zipper fusions of both BACTH plasmids were always run alongside. Negative controls, containing the two BACTH plasmid types together but without inserts, and selected pairs of BACTH plasmids with inserts combined with a respective empty plasmid of the complementary compatibility type were also tested alongside (not shown). For all tested interactions with positive outcome, exclusively the BACTH plasmid combination for each interaction pair with the highest quantitative outcome of interaction is included in the matrix. If all assayed interactions were negative, one negative result is shown. Human CEACAM1 self-interaction (homodimer) was used as a control condition for the human CEACAM1 functionality in BACTH. The matrix fields designated with crosses refer to Cag-Cag interactions shown in Fig 1, which are not relevant for HopQ or HopZ interactions and therefore not included in this panel again. **B)** BACTH results of b- galactosidase activity of interactions between HopQ (types I and II) and HopZ (types I and II) as homo- and heterodimers, depicted as bar graphs (absolute values in Miller units). Detection limit of positive interaction was at 1.5-fold the negative control value as in Fig 1. **C)** BACTH results of relevant β-galactosidase activities of interactions between HopQ (types I and II) and CagT4SS outer proteins, depicted as bar graphs (Miller units); **D)** BACTH results of b-galactosidase activity of interactions between HopZ (types I and II) and CagT4SS outer proteins, depicted as bar graphs in absolute values (Miller units). **E)** Shows a complete graphical network model of novel interactions between *H. pylori* CagT4SS outer proteins and outer membrane proteins HopQ and HopZ, summarizing all results from Fig 1 and Fig 2; green lines are novel interactions found by BACTH, blue line indicates one single interaction not found in BACTH but verified by plate-binding assay (see S5 Fig), which confirms the potential of the CagC (VirB2) and CagL (VirB5) orthologs of the T4SS to bind to each other. **F)** Novel Bacterial Three-Hybrid Assay (BAC3H) results of relevant combinations of proteins (results shown in Miller units) suggest competitive interactions between HopQ (types I and II) and CagL as well as between CagM, CagL and CagI. The legend tables included in each of the panels B, C, D, F contain the full information about the plasmid combination used in each assay. Detection limits of the BAC3H were determined by negative controls containing all three plasmids as empty versions which were included in all experiments, and were at or below 1.5-fold of the Miller units of the negative control. Each result consists of three data points from three independent measurements. Statistical analysis for significant difference in B, C, D, F was performed for the comparison between each sample versus the negative control (Student’s *t*-test, two-tailed unpaired), and significant p values are shown above each sample. *p<0.05; **p<0.01; ***p<0.001; ****p<0.0001. **G)** Immunoblot from *E. coli* cell lysates (supernatant (s) and pellet (p) fractions) detecting expression of the proteins assayed in the BAC3H. Blot incubated for detection with α-T25 antiserum (rabbit, 1:20,000) αCagM antiserum (rabbit, 1:10,000) and αCagL antiserum (rabbit, 1:20,000). Soluble (s) as well as insoluble (p) fractions of the same samples were tested for each sample to determine solubility of the expressed proteins. Fixed amounts of 10 µg protein were loaded on each lane. Antisera against bacterial GAPDH was used as a loading and fractionation control (as in S2 Fig) and confirmed even loading on the blots.

Due to earlier findings of HopQ interaction with human CEACAMs (56), we also decided to use human CEACAM1 as a positive control for HopQI binding to verify the validity of this assay for detecting the human-bacterial protein interactions. As expected for HopQ due to previous findings (15, 47, 56), HopQ large <u>b</u>inding <u>d</u>omain (lbd), contained in the cloned construct (amino acids 22 to 468), of which segments between aa100 to aa160 (including clasped loop CL1 and loop CL1-H4 (56) – the latter included in Loop2 as designated in the present study; S1A Fig) have been found to be crucially involved in binding hCEACAM in a co-crystallization analysis (56), interacted with human CEACAM1 in the BACTH set-up (Fig 2A). We also observed the known CEACAM1-CEACAM1 homotypic interaction. By this assay, also three novel hCEACAM1 interactions (with HopZI, CagC and CagL; Fig 2A; S2G Fig) were suggested.

### Novel Bacterial-Three-Hybrid assay was developed to refine the new model based on CagT4SS and Cag-Hop interactions

Next, we established and performed selected bacterial three-hybrid (BAC3H) assays for novel CagT4SS and HopQ interactions, with the aim of achieving a better understanding of a selected set of protein-protein combinations. By co-transforming a third expression plasmid without an interactive protein fusion (pAB148 derivative (Methods)), we aimed to indicate a potential competitiveness or synergistic interactions, through a decrease or increase in comparison to the ß-galactosidase activity measured in parallel in the corresponding two-hybrid combinations.

As shown in Fig 2F, the introduction of CagL as a third interaction partner to the HopQI - HopQI or HopQII – HopQII self-interaction led to a significant reduction for both interactions, as quantitated by ß-galactosidase activity (Fig 2F). This suggests that CagL can weaken HopQI (or HopQII) self-interactions (Fig 2F). For testing the second hypothesis that the interaction between the two surface proteins CagI and CagL interferes with the CagI - CagM interaction, we combined the latter three proteins by adding non-tagged CagL, which indeed resulted in a strong reduction of the CagI-CagM galactosidase measurement (Fig 2F), while soluble CagI presence was increased in Western blot (Fig 2G). Although this has to be interpreted with caution, the apparent competition may imply that an alteration of interactions occurs for selected transported Cag surface proteins, including CagI, CagL, CagH (40): we suggest they may undergo transient CagM interactions (22), which can accompany their transport through the system, before they coalesce on the bacterial surface.

### Direct physical interactions of CagL, CagN, CagA, HopQI, HopQII, HopZI, and HopZII were verified and quantitated in pair-wise combinations using Biolayer Interferometry (BLI) and plate-binding assays

In order to better characterize and quantify the newly discovered protein-protein interactions, the genes coding for proteins CagL, CagN, CagA, HopZI, HopQI and HopQII (the latter cloned in two variants, from strains PNGhigh12A [*cag*PAI-positive strain; designated as HopQII throughout the manuscript] and K26A [hpAfrica2 strain from Khoisan population, *cag*-negative] (https://enterobase.warwick.ac.uk/)) were cloned into expression vectors, expressed in *E. coli* Rosetta pLysS, and purified by affinity chromatography (S3A Fig and Methods) (22). The phylogeographic population hpAfrica2 is always devoid of a *cag*PAI and thought to predate the acquisition of the *cag*PAI by *H. pylori* (10) and is therefore an interesting variant to test for interactions of HopQ(II) with Cag proteins. The *hopQII* allele of an hpAfrica2 strain should therefore not be affected by prior selection pressure exerted by possible interactions with CagT4SS components. CagC was cloned and expressed as a GST fusion for interaction assays (Methods), to improve its solubility, and purified alongside non-fused GST (negative control).

The purified recombinant proteins (S3A Fig) were used for Biolayer Interferometry measurements using amine-reactive 2^nd^ generation biosensors with covalent coupling of the ligand. BLI (Octet) measurement of protein-loaded biosensors for association and dissociation of an analyte in solution added in increasing concentrations also permitted us to calculate K_D_ values for the tested interactions (Table 1). We performed those for newly detected CagT4SS protein interactions and also included human proteins CEACAM1 and integrin-α5β1 (commercial purified proteins) for selected interaction partners (Fig 3A, 3B). We tested most interactions at two different pH values, pH 6 and pH 7, since the natural habitat of *H. pylori* deep in the gastric mucus alternates naturally between neutral and (slightly) acidic (63, 64). Table 1 shows the calculated K_D_-values for the 23 protein-protein interactions tested in BLI at both pH 7.0 and pH 6.0, assuming a 1:1 interaction model.

**Figure 3:**
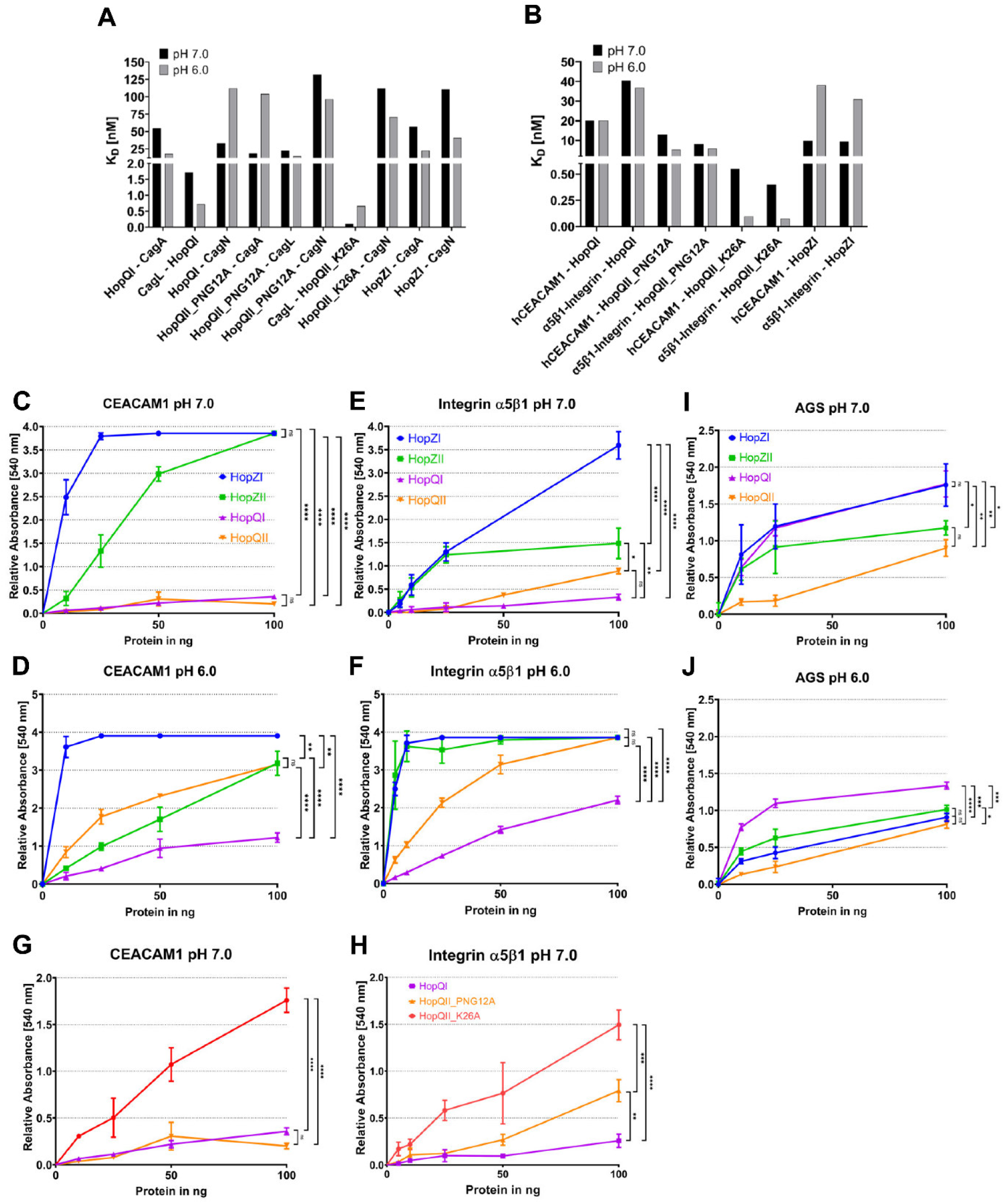
**Biolayer Interferometry and selected plate-binding assays to detect concentration-dependent binding of purified recombinant HopZ and HopQ (variants) to T4SS outer proteins, human gastric epithelial cells and human proteins (CEACAM1, integrin-**α**5**β**1).** Assays detect differential binding of HopQ and HopZ and their type variants, and pH-dependent substrate binding. **A)** and **B)** Overview bar graphs of biolayer interferometry assays quantitate different affinities of HopQ or HopZ interactions with *H. pylori* CagT4SS proteins (A), as well as HopQ or HopZ interactions with human CEACAM1 or integrinα5β1 (B), and demonstrate that some affinities differ between pH 7 and pH 6. **A)** Different HopQ and HopZ types were purified and assayed in BLI against T4SS outer proteins CagL, CagA, and CagN. **B)** HopQ or HopZ interactions with human CEACAM1 or integrinα5β1 were tested by BLI. The interactions between ancient HopQII (strain K26A) or HopZI and either CEACAM1 or integrinα5β1 were pH-dependent, while the interactions between modern HopQI(26695) or HopQII(PNGhigh12A) types and the human receptors were rather not. Ancient type HopQII_K26A (cloned from Africa2 strain without a *cag*PAI) showed high-affinity interactions with human CEACAM1 or integrin-α5β1. The first-named proteins on the X axis were bound to the sensors, and the second-listed were used as soluble analytes in BLI. See full results of BLI assays in Table 1. We did not assay CagC and HopZII in BLI, which were only purified in small quantities due to low solubility. Panels **C)** and **D)** illustrate results of concentration-dependent binding in plate-assays for all four purified HopQ and HopZ types to human CEACAM1 (CEACAM1) at pH values of 7 and 6. For all those conditions, HopZI bound best to CEACAM1, while HopQI had the lowest binding characteristics in this assay system. pH 7 enhanced HopZ binding to CEACAM1 over pH 6. Panels **E)** and **F)** show results of concentration-dependent binding in plate-assays for all four HopQ and HopZ allelic variants to human integrin-a5b1 at pH 7 and pH 6, respectively. HopZI showed the strongest binding in both conditions and increased binding at pH 6. HopZII also exhibited stronger binding at pH 6 than at pH 7, while HopQI and HopQII binding to integrin was less influenced by pH variation. Panels **G**) and **H)** illustrate concentration-dependent comparative quantitative plate-binding assays of three different HopQ variants (HopQI, HopQII modern [PNGhigh12A] and HopQII ancient [K26A]) to human CEACAM or integrin-a5b1, respectively. All plate-binding assays were at least performed twice independently in triplicates. **I), J)** depict results of comparative multi-well plate-binding assay of all four HopQ and HopZ allelic variants to gastric epithelial cells (AGS) at two different pH settings; HopQI and HopZI showed the strongest binding at pH 7; while HopQI binding was stronger at pH 7 compared to pH 6, it bound better than HopZI at pH 6. HopQII showed the weakest cell binding under all conditions. For all panels: recombinantly expressed HopQI was cloned from strain 26695, modern HopQII from strain PNGhigh12A (here designated as: PNG12A), HopZI from strain Su2 and HopZII from strain 26695 (Methods); purification results and quantitation for all used proteins are shown in supplemental materials, S4 Fig; recombinantly expressed CagN, CagA (aa1-892), and CagL were cloned from strain 26695. For the plate-binding assays, we used bovine serum albumin (BSA) as a negative control analyte against all bound proteins, which did not indicate binding (not shown). Statistical differences in C, D, E, F, G, H, I, J were calculated using One-way ANOVA, followed by Tukey’s post-test, and are included in each panel. *p<0.05; **p<0.01; ***p<0.001; ****p<0.0001; ns = non-significant.

**Table 1.**
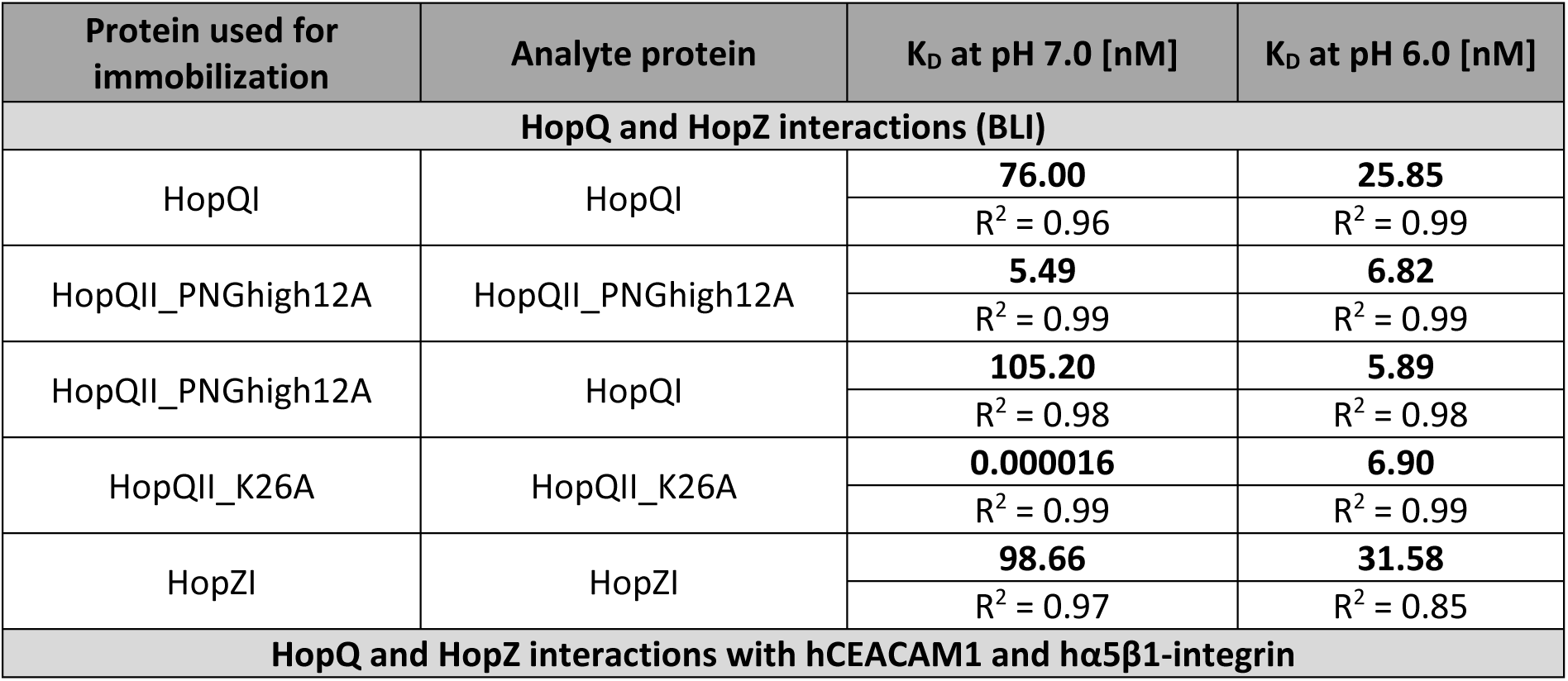
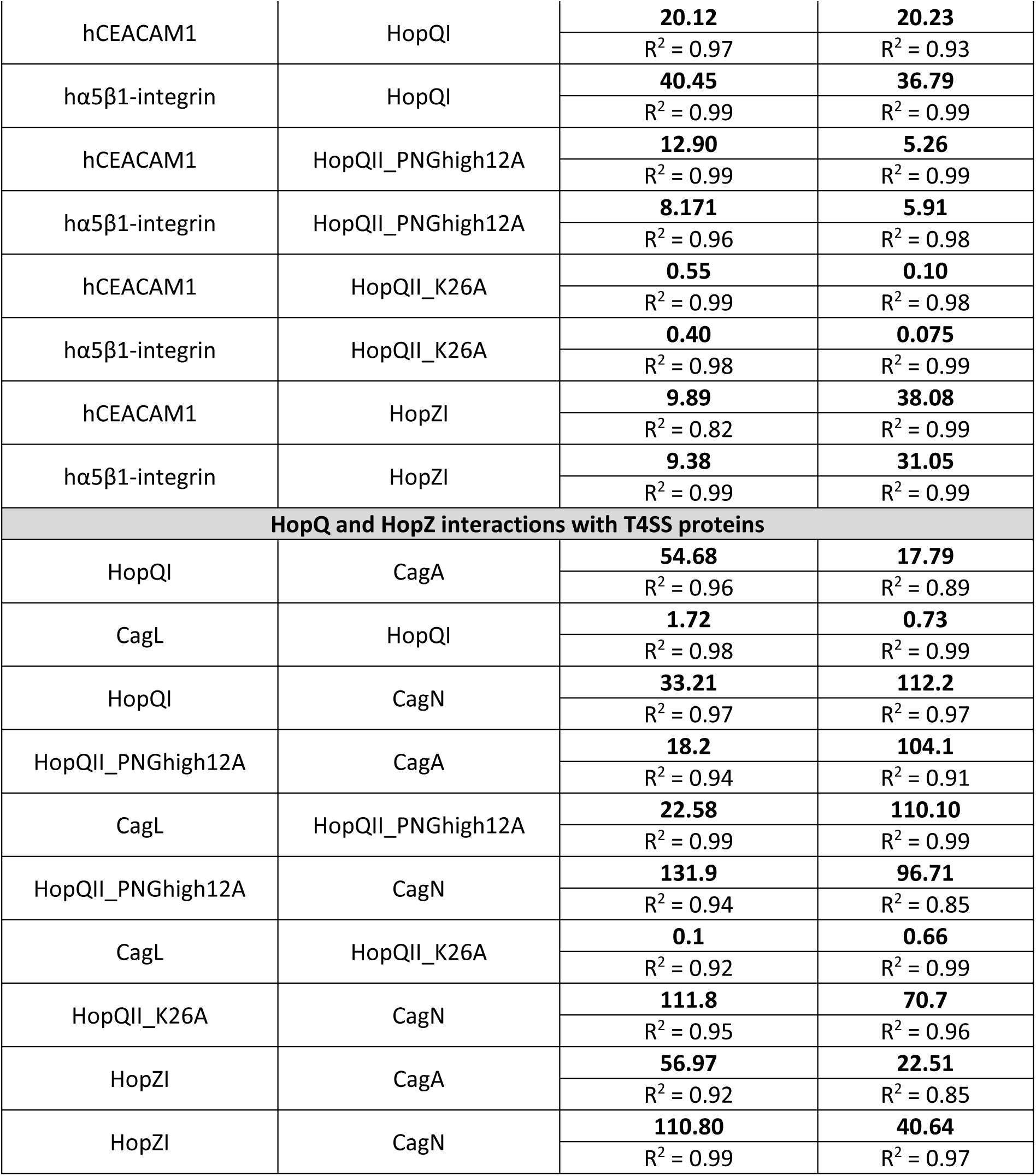
Kinetic parameters of HopQ and HopZ homo- and heterodimerization and their interactions with T4SS proteins CagL, CagN, CagA as well as host cell factors hCEACAM1 and hα5β1-integrin measured by Biolayer Interferometry (BLI) using the Octet system (Methods). A 1:1 model was used for the data fitting and KD determination. Two different variants of HopQII were purified and used, one variant cloned from the *cag*PAI-positive strain PNGhigh12A, from Papua New Guinea, and from strain K26A, a Khoisan hpAfrica2 isolate from an ancient *H. pylori* population which does not harbor a *cag*PAI (10). Only interactions that had shown a positive outcome in at least one plasmid combination of the BACTH assay and for one of the HopQ or HopZ allelic variants were tested in Octet. In addition, hα5β1-integrin interactions of HopQ and HopZ that could not be tested using BACTH, were investigated. Coefficient of Determination (R^2^) values (most were higher than 0.9) were calculated as a statistical measure to determine the data fit to the model. Due to remaining protein degradation products after purification (S4A Fig), KD determination of some interactions may not be reliable.

| Protein used for immobilization | Analyte protein | K<sub>D</sub> at pH 7.0 [nM] | K<sub>D</sub> at pH 6.0 [nM] |
| --- | --- | --- | --- |
| HopQ and HopZ interactions (BLI) |  |  |  |
| HopQI | HopQI | 76.00 | 25.85 |
|  |  | R<sup>2</sup> = 0.96 | R<sup>2</sup> = 0.99 |
| HopQII_PNGhigh12A | HopQII_PNGhigh12A | 5.49 | 6.82 |
|  |  | R<sup>2</sup> = 0.99 | R<sup>2</sup> = 0.99 |
| HopQII_PNGhigh12A | HopQI | 105.20 | 5.89 |
|  |  | R<sup>2</sup> = 0.98 | R<sup>2</sup> = 0.98 |
| HopQII_K26A | HopQII_K26A | 0.000016 | 6.90 |
|  |  | R<sup>2</sup> = 0.99 | R<sup>2</sup> = 0.99 |
| HopZI | HopZI | 98.66 | 31.58 |
|  |  | R<sup>2</sup> = 0.97 | R<sup>2</sup> = 0.85 |
| HopQ and HopZ interactions with hCEACAM1 and hα5β1-integrin |  |  |  |
| hCEACAM1 | HopQI | **20.12** | **20.23** |
|  |  | R<sup>2</sup> = 0.97 | R<sup>2</sup> = 0.93 |
| hα5β1-integrin | HopQI | **40.45** | **36.79** |
|  |  | R<sup>2</sup> = 0.99 | R<sup>2</sup> = 0.99 |
| hCEACAM1 | HopQII_PNGhigh12A | **12.90** | **5.26** |
|  |  | R<sup>2</sup> = 0.99 | R<sup>2</sup> = 0.99 |
| hα5β1-integrin | HopQII_PNGhigh12A | **8.171** | **5.91** |
|  |  | R<sup>2</sup> = 0.96 | R<sup>2</sup> = 0.98 |
| hCEACAM1 | HopQII_K26A | **0.55** | **0.10** |
|  |  | R<sup>2</sup> = 0.99 | R<sup>2</sup> = 0.98 |
| hα5β1-integrin | HopQII_K26A | **0.40** | **0.075** |
|  |  | R<sup>2</sup> = 0.98 | R<sup>2</sup> = 0.99 |
| hCEACAM1 | HopZI | **9.89** | **38.08** |
|  |  | R<sup>2</sup> = 0.82 | R<sup>2</sup> = 0.99 |
| hα5β1-integrin | HopZI | **9.38** | **31.05** |
|  |  | R<sup>2</sup> = 0.99 | R<sup>2</sup> = 0.99 |
| HopQ and HopZ interactions with T4SS proteins |  |  |  |
| HopQI | CagA | **54.68** | **17.79** |
|  |  | R<sup>2</sup> = 0.96 | R<sup>2</sup> = 0.89 |
| CagL | HopQI | **1.72** | **0.73** |
|  |  | R<sup>2</sup> = 0.98 | R<sup>2</sup> = 0.99 |
| HopQI | CagN | **33.21** | **112.2** |
|  |  | R<sup>2</sup> = 0.97 | R<sup>2</sup> = 0.97 |
| HopQII_PNGhigh12A | CagA | **18.2** | **104.1** |
|  |  | R<sup>2</sup> = 0.94 | R<sup>2</sup> = 0.91 |
| CagL | HopQII_PNGhigh12A | **22.58** | **110.10** |
|  |  | R<sup>2</sup> = 0.99 | R<sup>2</sup> = 0.99 |
| HopQII_PNGhigh12A | CagN | **131.9** | **96.71** |
|  |  | R<sup>2</sup> = 0.94 | R<sup>2</sup> = 0.85 |
| CagL | HopQII_K26A | **0.1** | **0.66** |
|  |  | R<sup>2</sup> = 0.92 | R<sup>2</sup> = 0.99 |
| HopQII_K26A | CagN | **111.8** | **70.7** |
|  |  | R<sup>2</sup> = 0.95 | R<sup>2</sup> = 0.96 |
| HopZI | CagA | **56.97** | **22.51** |
|  |  | R<sup>2</sup> = 0.92 | R<sup>2</sup> = 0.85 |
| HopZI | CagN | **110.80** | **40.64** |
|  |  | R<sup>2</sup> = 0.99 | R<sup>2</sup> = 0.97 |

The BLI measurements permitted a quantitative assessment of important novel interactions revealed in the BACTH screen. They also clearly demonstrated that the tested pH value has a significant influence on the binding affinity of many interactions including T4SS proteins and HopQI or HopZII. Many tested interactions were stronger at pH 6.0 when compared to pH 7.0 (higher K_d_ at pH 7) (Table 1; Fig 3A, 3B). For HopQII interactions, this difference was less apparent. HopQI and HopQII self- interactions and hetero-oligomerization were quantitated, with high affinities, as well, but showed less pH-dependence in BLI, in particular the HopQII homotypic interactions. (Table 1). Results of analytical size-exclusion chromatography (SEC) performed for purified recombinant HopQI protein further supported the formation of homotypic oligomers (possibly including tetramers) in solution (S3B Fig). HopQI was confirmed in BLI to interact directly with purified CagN, CagL (medium affinities), and with CagA at high affinities (Table 1; Fig 3A). HopQII self-interaction and HopQI-CagN interactions were pH- dependent and stronger at pH 7, while HopQI-CagA and -CagL interactions were determined to be of higher affinity at pH 6 (Table 1). Moreover, HopZI self-interaction was quantitated as high-affinity. HopZI had lower affinities to CagA and CagN than HopQI, which were stronger at pH 6 (Table 1 and Fig 3A). The affinities of interactions with CagL and CagA were significantly stronger for HopQII (PNGhigh12A) as compared to HopQI at the same conditions (pH 7). Both the homo-dimerization assay of HopQII (variant PNGhigh12A) as well as the hetero-dimerization of HopQII (PNGhigh12A) and HopQI in BLI had higher calculated affinities than the HopQI homotypic dimerization (Table 1; S4A Fig).

As for the additionally BLI-tested interactions with human receptor proteins, HopQI had a lower affinity for both CEACAM1 and integrin-α5β1 than HopQII and HopZI (Table 1). While HopQI and HopQII_PNG12A showed no or little pH-dependency for integrin or CEACAM binding, CEACAM1 and integrin binding for HopQII_K26A (from *cag*PAI-negative hpAfrica2 strain K26A; see also below) and HopZI were pH-dependent and exhibited lower or higher affinities, respectively, at neutral pH 7 (Fig 3B; Table 1). Since the non-membrane portion of HopZII which was also cloned and expressed for purification in *E. coli* (S1 Fig) was extremely insoluble and therefore difficult to purify in sufficient amounts, we did not perform BLI with this protein. To complement this deficit, we developed a multi- well plate assay (Methods) to test binding with smaller amounts of protein, which also allowed for a direct comparative testing. CagC interaction with CagL, for which we could also not perform BLI due to low yields of purified CagC, was likewise confirmed using a plate assay (S4B Fig).

Multi-well plate-binding assays also confirmed, concentration-dependently, the interactions of all four HopQ and HopZ alleles with human CEACAM1 and integrin (Fig 3C, 3D, 3E, 3F), which were not testable for the integrin heterodimers with Hop proteins in BACTH. Direct comparative assays in multi-well plate format performed for the four HopZ and HopQ variants with human CEACAM1 showed the strongest interaction of HopZI, followed by HopZII and HopQII (PNGhigh12A), and finally HopQI (Fig 3C, 3D). This was the case at both tested pH values, with slightly higher HopZ binding detected at pH 7. Similar results as for hCEACAM1 in the plate-binding assays were also obtained for integrin (α5β1) binding to both HopZ and HopQ variants, with the highest binding determined for HopZI and HopZII, compared to lower binding for both HopQ allelic types (Fig 3E, 3F). The highest binding for the HopZ variants to integrins was preserved at pH 6, while all of the four Hop proteins showed increased binding at pH 6 compared to pH 7. The latter results, in addition to the newly discovered Cag to Cag protein and Cag protein to HopZ or Cag protein to HopQ interactions, also add a complex layer of multiple integrin-CEACAM-Cag-Hop interactions (S2G Fig) to the proposed Cag surface protein assembly constituents.

### Archetypal HopQII from Africa2 strains reveal higher-affinity protein-protein interactions to human receptors integrin and CEACAM in comparison to current, Cag-associated, HopQI allelic variants

Different allelic variants of HopQ from different *H. pylori* populations exist, which have evolved in the presence or absence of a CagT4SS. In order to test for potential differences between them, we tested both, a current (“modern”) *hopQII* allele, cloned from a CagT4SS-positive *H. pylori* strain (PNGhigh_12A, isolated from Papua New Guinea), and the ancestral (“ancient”) sequence-diverse *hopQII*, from the *cag*-negative hpAfrica2 population strain K26A, for interactions with Hop and Cag proteins and human receptors using BLI and plate-binding assays. Using BLI, binding of ancient HopQII K26A to human receptors integrin and CEACAM1 was found to have higher affinities than binding of modern HopQII or HopQI to these receptors (Table 1), and more pH-dependent for human receptor binding (Fig 3B) than modern HopQII (PNGhigh12A). In multi-well plate binding assays, performed comparatively for the three purified HopQ variants at pH 7, ancient HopQII bound much better to integrin (Fig 3G) and CEACAM (Fig 3H) than modern HopQII and HopQI, corroborating the BLI assays. Hence, our results suggest that ancient HopQII has higher affinity to human cell surface receptor proteins than modern, presumably CagT4SS-adapted, HopQII. Modern HopQII exhibited an intermediate binding phenotype (Table 1; Fig 3F). We therefore suppose (see also discussion), below supported by functional assays, that modern HopQII alleles, if expressed, have sufficiently adjusted to the CagT4SS to functionally complement HopQI in this task, possibly by evolving towards weaker binding to human cell receptors. CagN binding affinities to the different HopQ alleles were not very different in BLI (high nanomolar range), however slightly pH dependent (Table 1), with the highest binding for HopQI at neutral pH. CagL-HopQ binding was more divergent, with very high affinities to ancient HopQII (from *cag*T4SS-negative strain) at neutral pH, followed by HopQI, and with the lowest affinity towards modern HopQII type (Table 1; Fig 3A). HopQI had lower affinity to CagL at pH 6, while modern HopQII showed similar affinities to CagL at both pH values.

### Differences between HopQ and HopZ in interaction with human cells

Since our above-described results of novel single-protein interactions also extended to novel interactions of both HopQ and HopZ towards human cell-surface exposed receptors, we also designed test systems to detect and roughly quantitate binding differences of the different Hop alleles to human cells. For this purpose, gastric epithelial cell lines AGS and NCI-N87 and HEK293-T cells were cultured to confluent monolayers in 96-well plates and preserved by aldehyde fixing. Subsequently, recombinantly expressed and purified HopQ and HopZ variants were co-incubated with the fixed cell layers in different concentrations in order to detect binding. The binding assays were performed at both pH values of 7 (Fig 3I) and 6 (Fig 3J). For AGS cells, HopQI and HopZI bound more avidly to the AGS cell surface than HopZII and modern HopQII (Fig 3I, 3J). HopQI and HopZI bound about equally well in this assay at pH 7 (Fig 3I). At pH 6 (Fig 3J), HopQI maintained highest binding to AGS cells, followed by HopZII, while HopZI showed relatively decreased binding. Highest HopQI binding of all the four tested variants was also confirmed for a second gastric epithelial cell line, NCI-N87 (S4C, S4D Fig), however in this line, HopZII was the second-best binder at both pH 7 and pH 6 (S4 Fig), indicating a host or cell type specificity. For NCI-N87 cells, the effect was different, since all HopQ/HopZ variants seemed to bind more strongly at pH 6. The cell type-specific divergent result points strongly to a cell type and/or individual donor specificity for HopQ and/or HopZ binding, also in a pH-dependent manner. We predicted that those differences may potentially impact on functionalities with regard to the CagT4SS, which we tested in the functional assays described below. For plate binding assays with HEK cells, which do not have CEACAM1 (47), we found that HopZ (both variants) were binding best, followed by HopQI (S4E Fig).

### **Strain- and** *cag*PAI-dependent **transcriptional activity and expression of** *hopQ*/HopQ **and** *hopZ*/HopZ

The isotype-specific binding results (above) and initial surface localization assays of HopQ and HopZ HiBiT-tagged bacteria (Methods) prompted us to ask the question whether strain-specific differences in HopQ and HopZ expression, for both allelic types, exist, which could instruct our strain selection for further assays. This information may seem to be tangential at first glance, but, in the very variable species *H. pylori*, this information is extremely important to interpret possible reasons for divergent phenotypes between strains, for example with respect to surface localization, human cell interactions, T4SS functionalities in vivo, or microscopic detection and localization. Transcript analysis using qPCR indeed revealed extremely divergent strain-specific transcript amounts of *hopQ* and *hopZ* alleles, despite the fact that the strains were all grown to mid-exponential growth phase under identical conditions (S5 Fig). Among the tested strains, Africa2 (*cag*PAI-negative), P12 and L7 showed the lowest *hopQ* expression levels (all normalized to 16S rDNA of the respective strain) (S5A, S5B, S5C Fig). Strains SU2, J99, B3 and D3a also had relatively low *hopQ* levels, in comparison to high expression in strains 26695 and N6 that we used for most of our present experiments. For *hopZ*, the transcript amount variation between strains was similarly strong (S5D Fig). The amounts by qPCR and Western blot were generally at much lower levels than for *hopQ*. Isogenic deletion mutants in the *cag*PAI ((13); three strains tested) showed minor changes in *hopQ* and *hopZ* transcript amounts (S5B, S5E Fig) in comparison to parental strains. As already known, strain 26695 was switched OFF for *hopZ*/HopZ by CT repeat length variation (61), but still maintained *hopZ* transcript by qPCR (S5D, S5E Fig).

Western blots using specific custom-produced antisera against HopZ (both alleles) (60) and HopQ (this work) confirmed the strongly variable strain-specific protein expression of HopQ and HopZ at the protein level in various wild type isolates (S5F Fig). As expected, in particular *hopZ* alleles with CT- dinucleotide repeats in predicted OFF status (frame-shifted ORF with early stop codon) did not yield detectable full-length HopZ protein in the Western blots (S5F Fig). Of the tested wild type strains, the *cag*PAI-negative Africa2 isolate showed the lowest protein expression of HopQ (S5F Fig).

### High-resolution fluorescence microscopy with dual tagging verifies partial co-localization of OMPs HopQ and HopZ with surface-localized CagT4SS protein CagN

The surprising determination of novel interactions between the outer membrane proteins HopQ and HopZ with CagT4SS proteins in vitro prompted us to use high-resolution fluorescence microscopy to verify whether the same proteins may colocalize in bacteria in situ. For this purpose, we initially established single HiBiT-tag insertions and their surface localization in HopQ and HopZ. We generated amino acid alignments and AlphaFold 3 structural models (S1A, S1B, S1C Fig) for full-length HopQ (56) and HopZ to reveal potential surface-associated loops. Three loops in HopQ and two in HopZ were selected for the insertion of tags (S1 Fig) that served for verification and quantitative detection of their surface localization (Methods, S6 Fig). Loop2-insertion in HopQ and LoopY-insertion in HopZ turned out to be the best candidates for surface detection of the proteins (S6A – S6G Fig). Engineered *cag*- negative (deleted *cag*PAI) isogenic mutants of strain 26695, containing a HiBiT insertion in HopQI (loop 2), showed no differences in surface expression of HopQI (S6H Fig), suggesting that Hop surface expression is not per se related to or influenced by *cag*PAI presence or function.

Subsequently, in order to develop strains suitable to identify localized protein-protein interactions in situ, in addition to the single HopZ and HopQ loop-tag fusions (above and Methods), we established internal HiBiT fusions of CagN and CagL, which both have been reported as surface-located “outer” proteins of the CagT4SS (22, 36, 40, 65), in strain N6. CagN-HiBiT was generated as a fusion in an internal loop that has been assumed due to preliminary predictions (AlphaFold) to be surface-located in the protein (S6 Fig and Methods) and was introduced into *H. pylori* (strain N6) as a plasmid construct, providing about 5-fold overexpression. CagL is also located at the bacterial surface, where it interacts with human integrins (37, 65, 66). Therefore, we designed CagL-HiBiT as a second tagged construct, with the HiBiT tag inserted at the N-terminus of CagL, immediately downstream of its proposed leader peptide cleavage site. CagL-HiBiT was chromosomally integrated in strain N6 using allelic exchange mutagenesis. Both HiBiT insertions were shown by Western blot to be expressed in the respective strains (S6G Fig; not shown for CagL). Both insertions were also verified using the HiBiT surface detection methodology, using reconstituted luciferase (Methods), to be surface-associated in intact bacteria (S6F, S6I Fig). The CagN-HiBiT insertion construct was highly expressed and strongly surface- detectable, using HiBiT surface detection (S6F Fig), and also in fluorescence microscopy of non-permeabilized bacteria (Fig 4A, 4B), making this construct more amenable for double-tagged analysis. The N-terminal CagL-HiBiT fusion construct yielded markedly lower surface fluorescence intensity in HiBiT surface detection (S6I Fig). Both CagN-HiBiT and CagL-HiBiT strains, albeit at lower intensity for CagL, presented with a similar dot pattern on the surface of the bacterial cell, with three to four dots on average detectable per bacterium in fluorescence microscopy (S6J Fig; and Fig 4). For the interaction partners HopQI and HopZII, we created V5 epitope tag insertion constructs in the same predicted extracellular loops described above for the HiBiT insertions, Loop2 and LoopY (S1B, S1C Fig), which are predicted not to be relevant for the overall structure of the proteins (S2 Table for mutagenesis plasmids). Using the Hop-V5 insertions and CagN-HiBiT constructs, ultimately, *H. pylori* strains with double tags were generated by allelic exchange mutagenesis in strain N6 that contain both, a V5 tag in HopQI (Loop2, see alignment in S1A Fig) or HopZII (LoopY; protein alignment in S1A Fig) and the HiBiT insertion in CagN (S6F, S6G Fig, S1 Table). Using those NQ1 and NZ10 double-tagged strains, we first verified that they are expressed and surface–exposed (S6F, S6G Fig), and do not lose CagT4SS functionality with respect to heptose-dependent pro-inflammatory signalling on human epithelial cells (S6J Fig). Subsequently, we performed fluorescence microscopy on double-labelled non-permeabilized bacteria (Fig 4), detecting both CagN and Hop proteins on the bacterial surface in situ. CagN localized on the bacterial surface in dot-like patterns (two to eight per single bacterium). HopQ was clearly membrane associated, as expected for an outer membrane protein. It localized around the entire periphery of the single bacterial cells (Fig 4A) with some dot-like accumulations, which partially co- localized with the CagN label (Fig 4E). HopZ, in contrast, showed a rather dot-like distribution in the periphery of the single bacterial cells, with smaller dots also partially co-localizing with tagged CagN dots (Fig 4C, Fig 4E)). Quantitative assessment (paired counts) of HopQ and CagN, HopZ and CagN, respectively, showed that more than two thirds of the dots appeared to co-localize in each combination (Fig 4E).

**Figure 4:**
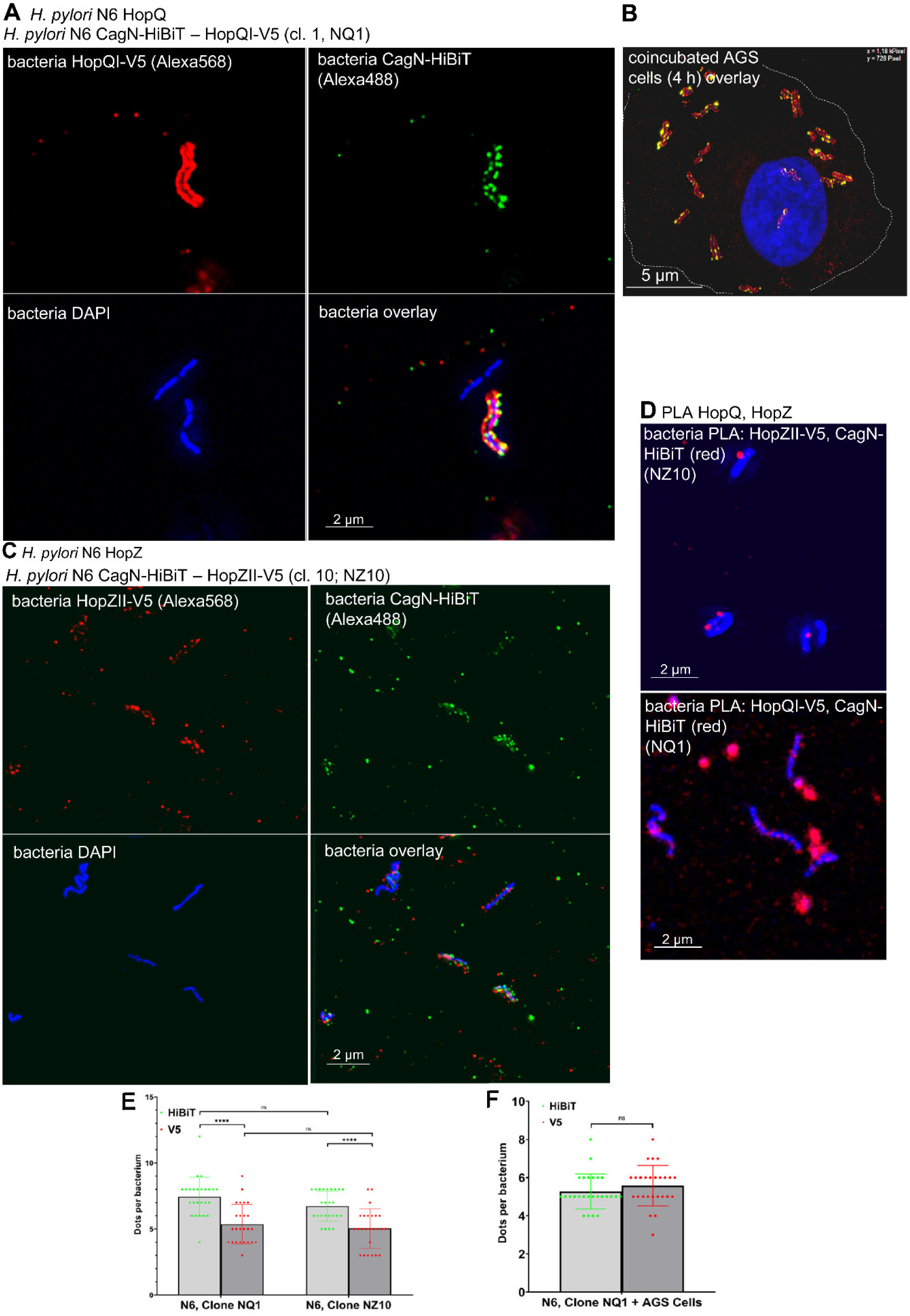
High-resolution microscopy colocalization of CagN and HopQ/HopZ in bacteria without and with coincubation with human gastric epithelial cells. *H. pylori* (strain N6) bacteria were doubly tagged with V5 (inserted in either HopQ or HopZ) and with HiBiT tag in the CagN protein, which is a surface-associated protein of the CagT4SS. The doubly-tagged strains, fixed and non-permeabilized, were immuno-labelled at their surface with both rabbit anti-V5 antibody (1:250) and mouse anti-HiBiT (1:100) antibodies (Methods) and fluorescently-labelled secondary antibodies (green, anti-mouse Alexa 488-coupled and red, anti-rabbit Alexa564-coupled); high-resolution fluorescence microscopy was performed on the fixed samples. The labels are explained in the figure panels. **A)** doubly-labelled N6 HopQI-V5, CagN-HiBiT; bacteria only **B)** bacterial co-incubation of doubly-labelled N6 HopQI- V5, CagN-HiBiT with gastric epithelial AGS cells for 30 min, showing bacteria attached to the cell surface; white hyphenated line indicates the AGS cell periphery; **C)** doubly-labelled N6 HopZII-V5, CagN-HiBiT; bacteria only. **D)** *H. pylori* N6 bacteria in the absence of cells, expressing CagN-HiBiT combined with HopQI-V5 or HopZII-V5, subjected to proximity ligation assay (PLA) and subsequent high-resolution microscopy; red color of the PLA label indicates direct interactions or very close apposition of the two proteins. Negative controls of bacteria not expressing HiBiT or V5 tags and incubated with the same antibodies were negative for any antibody staining for V5 or HiBiT (not shown). Panel **E)** quantitative dot counts in microscopy for double labelled bacteria (strain N6) with HiBiT-tagged CagN and V5-tagged HopQ (clone NQ1) or V5-tagged HopZ (clone NZ10), respectively (corresponding to panels A and B). Non-permeabilized fixed bacteria alone were surface-stained as above (A, B, C). HiBiT dots are highlighted in green. HopQ or HopZ dots (red), respectively, were only counted if they were colocalized (paired) with green HiBit dots in the same bacteria. **F)** quantitative dot counts in microscopy for CagN-HibiT, HopQ-V5 doubly-tagged bacteria co-incubated with AGS cells, stained as in B (strain N6, clone NQ1). HiBiT-dots were counted for CagN-HiBiT and all clearly recognizable dots for HopQ (red) were separately counted (not paired) in the same bacteria, where the majority of dots were co-localizing with green HiBiT dots. Statistics in E) and F) were performed by Student‘s *t*-test. Significance of differences: ****p<0.0001, ns = not significant.

**Fig 5.**
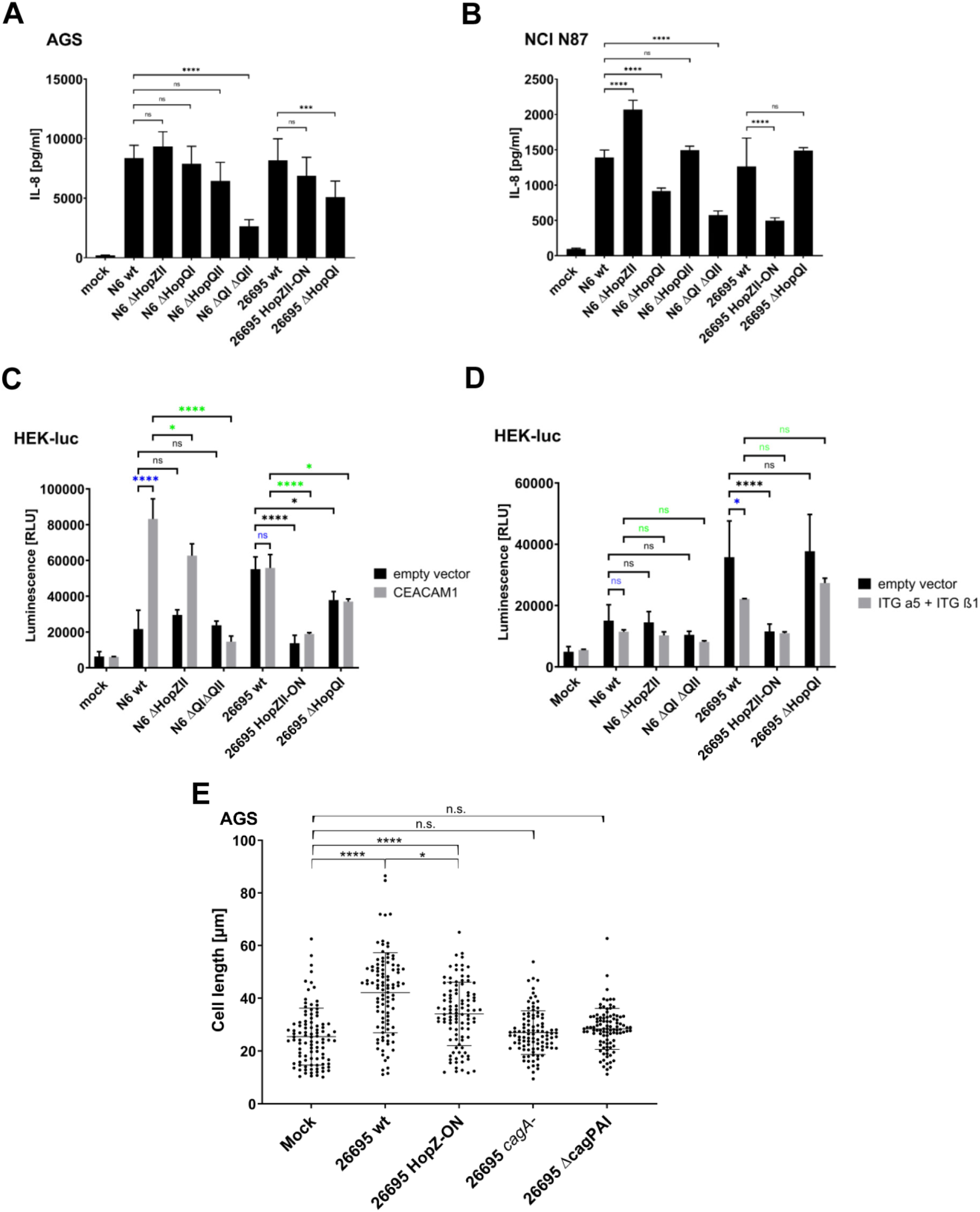
Characterization of. *H. pylori hopQ* **and** *hopZ* **mutants in cell activation assays in human epithelial cells shows that CagT4SS-dependent proinflammatory activation is dependent on both HopQ and HopZ and is inversely regulated by HopQ and HopZ.** Cell assays for pro-inflammatory activation were performed with various *H. pylori* strains and isogenic *hopZ* and *hopQ* mutants in two different gastric epithelial cell lines (AGS and NCI-N87, activation determined quantitatively as IL-8 release by ELISA) and HEK_luc NF-κB luciferase reporter cells (activation quantitated as reporter luciferase activity). **A)** AGS and **B)** NCI-N87 gastric epithelial cell lines were co-incubated with *H. pylori* strains N6 and 26695 and isogenic *hopZ* and *hopQI/II* mutants for 4 h; results of cell activation were quantitated using IL-8 ELISA from cell supernatants (at least four experiments were performed in triplicates); statistical analysis was performed for the comparison between wild type-co-incubated versus mutant- co-incubated cells for each mutant (One-way ANOVA with pairwise comparisons and Tukey’s post-test). **C)** HEK-luc reporter cells were transfected with human CEACAM1 expression plasmid or empty vector control; 24 h post transfection, cells were co-incubated with *H. pylori* strains and isogenic mutants for 3 h before recording activation (luminescence as relative luminescence units [RLU]); strain-specific aspect in CEACAM1-dependent T4SS function is revealed. **D)** HEK-luc reporter cells were transfected with human integrin-α5β1 expression plasmids or empty vector control; 24 h after transfection, cells were co-incubated with *H. pylori* strains and isogenic mutants for 3 h before measuring activation (luminescence as relative luminescence units [RLU]); for statistical analyses of differences between conditions in **C)** and **D),** again Two-way ANOVA with pairwise comparisons with Tukey’s test was calculated. *p<0.05; **p<0.01; ***p<0.001; ****p<0.0001; ns = non-significant. Color code of statistics: blue indicates significant differences between empty vector-transfected and expression vector-transfected; Black indicates differences between independent strains co-incubated with cells transfected with only empty vector, and green indicates differences between independent strains co-incubated with cells transfected with plasmids expressing either human CEACAM1 (C) or integrin-α5β1 (D). Co-incubation assays were performed in at least four biological replicates; All experiments were repeated at least twice on different days with similar results. **E)** CagA translocation assay (hummingbird phenotype, see Methods) for bacteria co-incubated with human gastric epithelial cells (AGS), developed in the following conditions: mock co-incubated, co-incubated with 26695 wild type (wt), 26695 HopZ-ON-complementant, 26695 *cagA* mutant, or 26695 DcagPAI mutant. Statistical evaluation by One-way ANOVA (Kruskal-Wallis test) with pairwise comparisons; **p<0.01; ****p<0.0001.

The results showed a partial co-localization of HopQ or HopZ with CagN (T4SS) in the absence of host cells in situ, although HopZ dots were smaller and much more difficult to spot visually. In order to detect closely appositioned, possibly interacting, proteins, rather than mere visual (partial) co- localizations, and to verify the results using another method, we employed Proximity Ligation Assay (PLA, see Methods) on the same double-tagged, non-permeabilized, bacterial strains (Fig 4D), followed by fluorescence microscopy. Thereby, we detected HopQ and CagN, and HopZ and CagN, respectively, in very close neighbourhood by the PLA label, supporting the notion of direct interactions between HopQI-CagN and between HopZII-CagN at the bacterial surface (Fig 4D). When we double-labelled HopQI together with CagN, without permeabilization, after bacteria had been co-incubated with gastric epithelial cells for 30 min, we detected an even more significant co-localization of most CagN dots with HopQI in microscopy (Fig 4B; Fig 4F). This was not the case for HopZ, which rather lost the majority of its bacteria-located immunolabel early (<= 30 min) during the cell co-incubation, for reasons yet unclear (not shown). Despite several attempts, we did not succeed in generating doubly- tagged bacteria for CagL-HiBiT together with HopQI-V5, which hindered the further evaluation.

### Functional characterization of interactions reveals role of novel interactions and a dampening function of HopZ in heptose-dependent proinflammatory signalling in human gastric epithelial cells

We ultimately assayed, whether the newly identified direct protein-protein interactions, in particular those involving the *H. pylori* outer membrane proteins HopQ and HopZ, had specific functional implications during interactions with human cells. We therefore generated isogenic insertion- inactivation mutants of strain N6 by allelic exchange mutagenesis targeting *hopZ* (corresponding to previous strategy in (60); Methods and S1 Table), and *hopQ* (including a *hopQI* and *hopQII* double- negative mutant in strain N6, which harbours both *hopQI* and *hopQII* alleles), by inserting antibiotic resistance cassettes into the respective chromosomal loci. We also constructed HopZ-ON constitutive mutants (complementants) in two strains, 26695 and L7 (10), for both of which the wild type is in the OFF status for *hopZ* due to the frame-shifted CT dinucleotide repeat region in the 5’-terminus of the gene (60, 61) (S1 Fig; S7A Fig; Methods). Both engineered HopZ-ON strains expressed HopZ in comparison to the original wild types, which did not show detectable HopZ protein (Western blots in S7B, S7D Fig).

For further functional evaluations, we then performed comparative cell infections (Fig 5, S7C, S7E Fig), both in AGS and NCI-N87 human gastric epithelial cells, measuring IL-8 cytokine secretion as a read- out. We obtained phenotypes for heptose-transport-dependent IL-8 secretion (13, 14) after short-term co-incubation (4 h) of the strains with the cell lines. *hopQI-hopQII* double mutants in strain N6 and *hopQI* single mutants in strain 26695, as expected (15), exhibited a significantly reduced IL-8 release with AGS cell co-incubation (Fig 5A), supporting the known important role of HopQ in the functionality of the CagT4SS, reported in the context of human CEACAMs (15). IL-8 induction in AGS cells was not affected by the constitutive ON-switch engineered in *hopZ* in strain 26695 (Fig 5A, 5B) nor by the *hopZ* inactivation mutation in strain N6. Surprisingly, in the second gastric cell type, NCI-N87, in contrast, the HopZ-deficient isogenic mutant of N6 showed a significantly divergent phenotype compared to the wild type, inducing about 30% higher IL-8 secretion in the cells, in comparison to the bacterial wild type (Fig 5B). Furthermore, the constitutive HopZ-ON mutant in strain 26695 induced a significantly diminished IL-8 secretion (Fig 5B, S7C Fig). This was also the case for HopZ-ON in the second strain L7 (S7E Fig). The latter results emphasized a cell-type dependent, direct, involvement of HopZ in CagT4SS functionality. As another cell- and strain-specific phenotype, the *hopQI-hopQII* double mutant of strain N6 showed a significantly reduced IL-8 secretion phenotype in NCI cells (Fig 5B), while the 26695 *hopQ* mutant did not show an altered phenotype. Together with the newly detected direct interactions of HopZ with several CagT4SS proteins and HopQ, we suggest a direct cooperative function of the two Hop proteins with the T4SS outer proteins which can inversely affect the functionalities for HopQ (decrease) versus HopZ (increase). In each strain-cell line combination, the quantitative effects were very stable, however, effects were clearly strain- and cell-type-dependent.

We also carried out a functional characterization of cell activation by *hopQ* and *hopZ* mutants in HEK293 cells (HEK_luc, containing a luciferase-based NF-κB reporter, Methods) in which cell type *H. pylori* major adhesins SabA and BabA (50, 64) play a less important role, due to the lack of specific cell surface receptors, which also includes a lack of CEACAM1. Only for strain 26695, this setting revealed a significant contribution of HopZ, but not HopQ, in CagT4SS-mediated heptose transport-induced NF-κB activation, quantitated as luciferase induction, under all conditions; HopZ-ON mutants showed reduced reporter cell activation by more than 60% during the early co-incubation of 3 h (Fig 5C, 5D) or reduced IL-8 production (data not shown), in comparison to the HopZ-deficient 26695 wild type strain. N6 and its isogenic mutants had no strong phenotype and rather low activation in the HEK cell model when the cells were co-incubated plain or empty vector-transfected (Fig 5C, 5D). As a second T4SS- dependent, transport-dependent phenotype, determined by CagA translocation efficiency, that may also be influenced by HopZ, we analyzed cell elongation (termed hummingbird phenotype (13, 68, 69)) in AGS gastric epithelial cells (strain 26695). Converging with the reducing activity of HopZ on pro-inflammatory cell activation, a significant reduction of the exclusively CagA-dependent phenotype, revealing diminished CagA translocation, was quantitated for the constitutive HopZ-complementant in comparison with the *hopZ*-negative 26695 parent (Fig. 5E). *cagA*-mutant and *cagPAI*-deletion mutant were negative for the effect. Consistently, for the HopZ-deficient isogenic mutant in strain N6, an enhanced hummingbird phenotype was shown, indicating increased CagA transport (own data not shown).

In parallel, HEK-NF-κB reporter cells transiently transfected with plasmids to express human CEACAM1 or α5β1 integrin were also exposed to the same *H. pylori* strains, in order to verify a functional relevance of those two human receptors for HopZ function in the T4SS context. Interestingly, this revealed an important strain-specific difference: strain N6 was almost completely dependent on CEACAM1 expression for NF-κB activation and showed the known, enhanced, HopQ-dependent T4SS functionalities only when CEACAM1 was expressed (Fig 5C). The effect of *hopZ* in this strain was minor in HEK cells under all test conditions. For strain 26695, in contrast, T4SS pro-inflammatory function was not CEACAM1-dependent at all and showed similar phenotypes, with a significant deficiency in activation of the 26695 HopZ-ON mutant, in the empty-vector conditions as well as with either CEACAM1 or integrin α5β1 expressed (Fig 5C, 5D). The HopZ-dependent decrease in cell activation for 26695 was not significantly influenced by additional CEACAM1 or integrin expression (Fig 5C, 5D).

We also tested other possibly relevant human CEACAMs (3, 5, and 6), as reported in (47, 57, 59), in the HEK reporter cells after transient transfection, of which hCEACAM3 promoted significantly elevated NF-κB activation with HopZ-ON mutant (S7F Fig), while hCEACAM5 and hCEACAM6 did not show significant divergence compared to wild type, as with CEACAM1 transfection (S7G, S7H Fig). This set- up also confirmed that HopQ-mediated T4SS increase for strain N6 (but not 26695) was largely CEACAM-dependent in HEK cells. Integrin expression in the HEK reporter cells generally dampened CagT4SS-induced responses but did not alter the relative contributions of HopQ or HopZ to the strain-specific phenotypes (Fig 5D).

## Discussion

The *cag*PAI genetic island geared for host cell manipulation by assembling a functional T4SS was presumably acquired by *H. pylori* in a single event during evolution more than 60,000 years ago, and has since further evolved to be genetically highly diverse, with particularly the genes coding for surface-exposed proteins of the presumed Cag outer protein complex underlying strong diversifying selection (10). These surface proteins of the *H. pylori* CagT4SS, reported to form a pilus-like structure under certain conditions of human host cell co-incubation (35), mainly CagL (VirB5 homolog), CagI, CagH, possibly supported by CagC (VirB2 homolog), are known so far to be important for the function of the export machinery and to directly or indirectly interact with host cell factors (35, 39, 44, 67). CagY with CagL, as well as CagL with CagI, and CagI with CagH were previously reported to interact directly (36, 40, 46).

Although in recent years, more extensive structural studies, utilizing the improving technology of high-resolution Cryo-EM or Cryo-ET, have contributed to a better understanding of the overall CagT4SS structure. However, even if it has already been proposed that a CagT4SS surface assembly or structure might exist (40, 67), in particular the composition and interaction of the surface-exposed Cag proteins, for instance CagI, CagL, CagN, CagC, and CagA, the definition of a potential CagT4SS surface protein complex or structure, and the interplay of Cag proteins with regard to host cell interaction and the export functions of the CagT4SS are still poorly understood (21, 30, 33).

In this study, bacterial two- and three-hybrid assays were initially used to fill the gaps of so far uncharted territory in CagT4SS protein-protein interactions, in particular of the CagT4SS surface proteins. Thereby, we aimed to uncover potential unknown interactions and interaction partners, in order to improve the understanding of the Cag apparatus and possibly also shed more light on the entire CagT4SS transport process. Positive interactions identified in these assays were further characterized by analyzing purified recombinant proteins for interactions by biolayer interferometry and further methods. In addition to HopQ (two alleles), we also included the sequence-variable and phase-variable closely related HopQ paralog HopZ (60, 61) (two alleles) as a possible functional interaction partner of the CagT4SS. While the interaction of both HopQ alleles with human CEACAMs, in particular CEACAM1, was studied before to some detail biochemically, structurally as well as functionally (56, 57), a receptor for HopZ on the human side was not yet known (59, 61). We newly identified here that HopZ interacts with itself, with HopQ, and with human CEACAM1, using BACTH, multi-well plate-binding assays and direct BLI interaction assays using purified recombinant HopZ. Surprisingly, we also identified and quantitated both HopQ and HopZ to interact with human integrin (α5β1) in binding assays. Despite our positive binding results and the established evidence that CagT4SS proteins such as CagL, CagI, CagY (44, 46, 66), and CagA (46, 70) bind to integrins, our functional results so far corroborate previous findings that tested integrins are not involved in important CagT4SS functionalities (15).

The Hop protein HopQ was revealed as new interaction partner of the CagT4SS surface proteins CagL, CagN, CagI, CagC, CagA, possibly contributing to a proposed CagT4SS surface assembly, and of human integrins (α5β1). Since we only cloned and tested the larger portions of HopQ and HopZ (lbd and sbd as defined in S1 Fig) that are not associated with the outer membrane-integrating beta-barrel structures, we can only safely summarize that the detected novel interactions are supposedly located predominantly in the extracellular (binding) domains of those Hops. HopQ (both alleles) was shown in the present study to directly interact in vitro with CagA and CagL at nanomolar affinities, in a strongly pH-dependent manner. HopQ also exhibited high-affinity interactions in vitro with CagN. While both proteins, HopQ and CagL, can play a role in bacterial adhesion to host cells (59–61, 65, 66, 71), and the interaction of HopQ with human CEACAMs plays a vital role in the translocation of CagA in some cell models, and also for the induction of early pro-inflammatory signaling (47), mediated by heptose metabolite transport (13), little has been known about the functional details of those processes. HopQ close paralog HopZ interacted with CagA, CagL, CagN, and CagC, as well as with integrin and CEACAM1. Interestingly, ancient HopQII, in contrast to modern, Cag-adapted HopQs, interacted more strongly with human receptors and less strongly with Cag proteins. Considering that frequent recombination between *H. pylori* strains occurs, which supports fast adaptation, this likely has led to the situation that most modern HopQI or HopQII protein variants have frequently interacted with Cag proteins in their strain genetic environment. This presumably created opportunities to adapt specific Hops over time to interaction with CagT4SS proteins in the highly recombinogenic species. HopQI bound best to gastric epithelial cells at all conditions in contrast to the other HopQ and HopZ variants, although this was not true for isolated human receptor CEACAM1- and integrin-binding.

The results of this study also suggest an interaction and partial co-localization of both the Hop proteins HopQ and HopZ in situ with proteins of the CagT4SS, and specifically Cag proteins such as CagN at the bacterial surface. CagN is linked to the other Cag surface proteins by the newly identified CagN interactions with both CagL and CagI, and by prior positive interaction studies of CagL with CagI, CagI with CagH, and CagL with CagY (36, 40, 46), the latter being an integral protein of the OMCC (33). Double immunofluorescence labelling with dual-tagged HopQ, or HopZ, and CagN proteins in intact non-permeabilized bacteria demonstrated partial co-localization of HopQI and, to a lesser extent, HopZII, with CagN (T4SS) in situ on the bacterial surface. HopQI in particular, which partially associated with CagN on the bacterial surface in the absence of host cells, co-localized more strongly during co-incubation conditions with human gastric epithelial cells. Close apposition of HopQI (and HopZII) with

CagN by PLA assay, likewise suggesting interaction, was detectable. This corroborates and expands previous fluorescence microscopy showing focal dot-like localization of CagY and CagT (VirB7) on the bacterial surface of *H. pylori* co-incubated with AGS cells (72). The propensity of the HopQ (and HopZ) proteins to form at least dimers, if not oligomers, also indicates a protein assembly, which may be associated with some Cag surface proteins. Taking all the findings into account, it seems well conceivable that Hop proteins may be clustering with CagT4SS surface proteins on the bacterial membrane, possibly even triggered by host cell interaction. The latter hypothesis needs to be explored in more detail in a future study. The finding also implies an important role of HopQ and HopZ for the function of the secretion system, not only through adhesions to host cell receptors, but also via direct interaction with the surface-exposed proteins of the proposed CagT4SS surface protein complex. It still remains to be proven that the interacting proteins actually form a structure or complex at the bacterial surface and close to the membrane-spanning secretion system under some relevant conditions. Future efforts at isolating a surface protein assembly, for instance by crosslinking and/or pull-down approaches in *H. pylori*, should help to clarify such questions, although manipulation may subvert assembly and it is challenging to distinguish between direct and indirect interaction in the complex setting. Very recently, a study reported a successful immune-purification experiment using the CagA chaperone CagF as bait which enriched CagA, low amounts of CagN, CagL, CagI, CagH, and high amounts of OMCC components from *H. pylori* solubilized fractions containing the OMCC_Cag_ structure, but this was so far not linked to the bacterial surface (73). This may be further evidence for the assembly of a larger complex, but its existence and localizations need to be verified.

The strong, pH-dependent, interactions of HopQ and HopZ with themselves as well as with CagA and CagL might indicate a structural change of the proteins in response to environmental conditions in the stomach niche, which might affect Hop, Cag and host cell proteins in a functionally convergent manner. A similar pH-dependent binding characteristic, mediated by a structural change, was shown before for BabA-receptor interaction (64) and for CagL tertiary structure and its adherence to human integrins (65). We also detected a pH-dependence of the binding of the four HopQ and HopZ variants to cells. In addition, cell-specific phenotypes of so far unknown causality were observed, as all four protein variants bound better to NCI-N87 cells at pH 6, while, to AGS cells, they bound better at pH 7. Interestingly, lower pH, which is supposed to elongate the structure of CagL (65), led to a reduced CagL-integrin binding, while, derived from our new results, the interaction of CagL with HopQ seems to be stronger at pH 6.0 than at pH 7.0 in solution. This may suggest a competitive action between host cell integrins versus CagL towards HopQ at different pH values, possibly pointing to some dynamics of the secretion apparatus components at the bacterial surface.

The initial screening results also revealed a previously unknown interaction of CagM with CagH, and CagM with CagI. CagL appears to compete with the CagM-CagI interaction. Together with our own results generated prior to this study for CagN and CagL (22), CagM is thereby shown to interact, we assume at least transiently, with the majority of CagT4SS proteins associated with the bacterial surface. Other novel interactions identified here were CagC-CagC (also reported in a recent preprint (34)), CagA with CagC, and CagC with CagI, with CagL, and with HopZ. These results may support a previously proposed function of CagM in guiding and transporting Cag proteins to the bacterial surface and suggest that affinity-driven competition could provide export-order prioritization (22). Despite extensive screening, no interaction of CagM with CagA was found here, in agreement with previous studies (31–33), which supports the hypothesis that CagA export may be CagM independent.

Quite surprisingly, cell- and strain-specific functionalities of the CagT4SS in combination with HopZ revealed that HopZ has the potential to act antagonistically to HopQ with respect to the pro-inflammatory function of the CagT4SS (in NCI-N87 and HEK cells), which depends on bacterial metabolite transport. This was also the case for CagA-dependent cell shape changes (hummingbird phenotype (68, 69)) which was decreased in the HopZ constitutive-ON complementant, while HopQ enhances CagA translocation (47, 59). This functional divergence between HopQ and HopZ may also be linked to our unexpected finding of strong expression differences of both HopQ and, in particular, HopZ between different strains. HopZ expression can be enforced by the previously identified CT- dinucleotide repeat-dependent switch mechanism (61). We speculate that HopZ might be selected for an ON switch when a changed condition such as pH or transmission to a new host requires its dampening function, or to OFF, when reduction of T4SS function is not needed in a stomach niche or altered environment of the host, e.g. during chronic infection. To expand our previous report that in a smaller strain collection, *hopZ* allelic type did not significantly correlate with the *cag*PAI (60), we found when analyzing a larger strain collection of the *H. pylori* Genome Project (74) that in particular the *hopZI* type shows a significant correlation with *cag*PAI presence (own data not shown). The dampening function of HopZ was not dependent on human CEACAM1 or integrin-α5β1 receptors in our assays. Human CEACAM3, but not CEACAM1, CEACAM5, CEACAM6, or human integrin-α5β1, seemed to influence HopZ-dependent cell activation. Strain differences comprised a strong dependence of CagT4SS functionality on CEACAM1 in specific strains (strain N6, similar to previously characterized strain P12 (15, 47)), versus a CEACAM-independent T4SS activity in other strains, such as 26695. In contrast to previous reports (15, 47, 58), HopQ-dependent, T4SS-mediated cellular activation was only promoted by CEACAMs in one (N6) of two tested strains. We can only speculate that HopQ, which shows sequence variation between strains, is more versatile than previously reported and can also use other cellular receptors to promote CagT4SS functionality.

Taken together, we have identified novel interactions between Cag outer proteins, including CagC, CagA, CagL, CagN and CagI. Furthermore, novel direct interactions between *H. pylori* outer membrane proteins HopQ and HopZ and multiple Cag proteins point to a direct functional involvement of those interactions in contact with and during transport processes towards human cells. This may render the possibility of a larger-order complex of proteins on the bacterial surface, mediating those functions, more plausible to further look into. Intriguingly, CEACAM1, as previously reported for human integrin (44, 46, 65, 67) not only bound HopQ and HopZ, but also interacted with some CagT4SS surface proteins (e.g. CagC, CagL), that we propose may form a CagT4SS surface protein assembly, possibly together with outer membrane proteins.

Limitations of the present study that should be covered in future investigations include clarification of a proposed assembly or structure at the surface of the CagT4SS and its possible components. A mechanism for the HopZ-mediated dampening functions on the T4SS-mediated activites is still lacking. There is also a persistent lack of information on other potential binding factors and receptors on the host cell side, since not all effects and cell-directed phenotypes that we detected here could be explained by CEACAM and/or integrin binding, in particular for HopZ, but also for HopQ. We also do not know yet whether other *H. pylori* outer membrane proteins can be involved. We assume this might be the case since multiple paralogs with structural homology to HopZ or HopQ exist in the *H. pylori* Hop family (48, 75), which indicates a certain functional redundancy. This may also help explain strain- and cell type-specific effects. Using other *H. pylori* mutants defective in known cell adhesins, specific Hop/OMP or CagT4SS proteins, or site-directed mutants of those proteins deficient in novel interactions, the knowledge about the interplay of interaction partners and their role in the transport of effectors through the T4SS should be intensified. Strain differences in the versatility of CagT4SS functions and the underlying causes should be better characterized, taking *H. pylori* strains of diverse origins into account. Furthermore, the role of pH and glycan modifications of cell surface proteins and potential receptors, which also might influence cell line-, cell type-, and individual binding or functional characteristics, still have to be investigated in larger molecular detail. Uncharted territory also still has to be filled for detailed molecular interaction analyses (including site-and domain-specific interactions between proteins), and stoichiometric considerations. With this newly generated knowledge of functional protein interactions and complex composition of CagT4SS outer protein assemblies, further functional research should be implemented to characterize and visualize structural detail and export processes of the T4SS in contact with human epithelial cells.

## Materials and Methods

### Bacterial culture conditions (*H. pylori*)

All described *H. pylori* wild type strains and mutants (Table S1) were grown on blood agar plates (Oxoid Blood Agar Base No. 2) under microaerobic conditions created by humidified Anaerocult C bags (Merck) in gas tight jars. Agar plates contained 10 % horse blood, amphotericin (4 mg/L), polymyxin B (2,500 U/L), vancomycin (10 mg/L), trimethoprim (5 mg/L) and, depending on the resistance markers in the mutant strains, also chloramphenicol (5 mg/L) and/or kanamycin (10 mg/L). *H. pylori* bacteria, which are rather slow growing (minimal duplication time ca. 4 h) were cultured for 20 to 24 h at 37°C on plates (reaching mid-log phase) before being used or passed to fresh plates for continuous cultivation. Bacteria were discarded or thawed freshly latest after passage number 10. *E. coli* strains (S1 Table) were grown on LB plates supplemented with relevant selective antibiotics, or in LB or TB broth for propagation of plasmids, BACTH assays (76), or protein expression. For comparative protein and RNA preparations from *H. pylori* wild type strains, some of which grow very poorly in liquid culture, we used 20 h plate-grown bacteria, as the bacteria auto-synchronize under those conditions, are highly proliferative and motile (checked by motility assay and microscopy).

## Protein methods and Western blot

Samples from *H. pylori* were prepared by harvesting the bacteria directly from 20 h grown blood agar plates into PBS with sterile cotton swabs. Cell disruption was performed by ultrasonication for 2 cycles of 45 s at 4°C (Branson sonifier, Power setting = 5). Soluble and insoluble fractions were separated by centrifugation (12.000 x g, 10 min, 4°C) and the protein concentration per fraction was determined by BSA assays. 10 µg of total protein per sample was loaded on 11.8% or 14% SDS-PAGE Gels run in Laemmli buffer (25 mA per gel) before tank blotting onto BA85 nitrocellulose membranes (Schleicher & Schuell) in Towbin buffer (300 mA, 2 h). Western blot membranes were blocked in 5 % skim milk in TBS-T (0.1% Tween) for 1 h at room temperature. Specific primary antibodies (see figure legends) diluted in 5 % skim milk in TBS-T were incubated at 4°C overnight. Secondary POX-conjugated goat-anti rabbit or POX goat-anti mouse antibodies were incubated for 1 h at room temperature in a 1:10.000 dilution in 5% skim milk in TBS-T. Immobilon HRP chemiluminescence substrate (Merck Millipore) was used for signal detection, and blots were imaged using a chemiluminescence imager (BioRad).

### **Generation of** *hopQI* **and** *hopQII* **single and double insertion-inactivation mutants**

The isogenic *hopQI* mutants (N6 and 26695) were generated by initially constructing a plasmid (Table S2) which contains an insertion of a kanamycin resistance cassette (AphA3’-III) in pUC19 between about 400 bp flanking arms of *hopQI* in a single overlap PCR cloning step (primers in Table S3). Subsequently, a PCR product was derived from the resulting plasmid, pSUS3322, using the flanking primers for *hopQI*. This PCR product was transformed into *H. pylori* wild type strain (N6). Clones were selected on kanamycin, expanded and characterized by PCR with relevant primer combinations and by Western blot using HopQ-specific antiserum for detecting lack of HopQI expression (not shown).

*hopQII* isogenic inactivation mutants were generated by cloning of *hopQII*[aa22-455] from *H. pylori* Khoisan26A into the pUT18c-vector backbone by enzymatic digestion with BamHI and KpnI. By reverse amplification of the construct via PCR (Roche, Expand High fidelity PCR system) a 299-bp deletion in the mid-segment of the gene was achieved as well as the introduction of a SpeI digestion site. Utilizing this SpeI site, a chloramphenicol resistance cassette was inserted for religation of the final construct (S1 and S2 Tables). *H. pylori hopQ* single- and double-insertion mutant strains were then generated by allelic exchange mutagenesis after natural transformation of *H. pylori* N6 wild type. Clones were selected on blood agar plates with chloramphenicol, kanamycin, or chloramphenicol and kanamycin for correct selection. The correct insertion-inactivation of the mutated *hopQII* gene or double insertion mutation was again checked via PCR and Western blotting. The existing *hopZ* mutant in strain N6 was used (60)

### Isogenic HopZ-constitutive-ON mutants for complementing HopZ loss of function by natural OFF switch

We also constructed constitutive HopZ-ON mutants (complementants) in two *H. pylori* strains, 26695 and L7 (10), for both of which the wild type is in the OFF status for *hopZ* due to the frame-shifted CT dinucleotide repeat region in the 5’-terminus of the gene (S1 Fig; S7A Fig). Briefly, wild type *hopZ* sequences together from each target strain with flanking nucleotide sequences up- and downstream of the CT-repeats were cloned into pUT18c vector using primers with suitable restriction enzymes (S3 Table). After confirmation of the correct plasmid insertions, each vector was amplified using specific primer pairs with the shortened CT-repeats (here: 7 CTs) and religated afterwards using T4 ligase. *E. coli* was transformed with the religated plasmids, and clones were screened via direct Sanger sequencing. Correct plasmids with shortened 7CT repeats (Table S2) were again PCR-amplified using the initial primer pairs. A PCR product with the correct CT-repeat length and flanking sequences was then used for transformation in *H. pylori* together with the *rdxA-cm* selection marker, according to the

MUGent strategy (77). Correct identity of the engineered strains, including expression of HopZ, was verified by PCR, Sanger-sequencing the altered nucleotide segment, and by Western blot (S7 Fig).

### **Construction of** *H. pylori* **HopQ- and HopZ-tag insertion mutants**

Amino acid alignments and AlphaFold 3 structural models (S1A, S1B, S1C Fig) were generated for full-length HopQ (56) and HopZ to reveal potential loops and surface-associated regions. Three loops in HopQ and two in HopZ were selected for the insertion of tags (S1 Fig) that could serve for verification of surface localization and for surface quantification of the proteins. We chose two short epitope tags (HiBiT and V5) to insert into HopQ and HopZ at the respective loop locations. For the generation of *H. pylori* strains with V5- and HiBiT-tagged HopQ and HopZ proteins (S1 Table), wild type sequences of *hopQI* from *H. pylori* 26695, *hopQII* from PNGhigh12A and *hopZII* from *H. pylori* 26695 cloned into pUT18c Vector (as used in BACTH assays) were utilized. By reverse amplification of the constructs via PCR (Roche, Expand High fidelity PCR system), a SpeI digestion site along with the 42 bp long V5-tag or 33 bp long HiBiT-tag sequence was introduced into pre-defined surface-exposed loop regions of the proteins (S1 Fig). By enzymatic digestion and subsequent religation of the PCR product, the tag insertion in the genes was achieved in the final plasmids (S2 Table). *H. pylori* mutant strains were then generated by natural transformation of N6 or 26695 wild type with a PCR product of the tag-inserted *hop* gene along with an PCR product containing a chloramphenicol or kanamycin resistance cassette with arms of homology of the *H. pylori rdxA* gene for homologous recombination into the chromosome and resistance selection of clones. The correct introduction of the tags was checked via PCR, using tag-specific primers (S3 Table), and Western blotting.

### **Detection and validation of** *H. pylori* **HopQ and HopZ tag insertion mutants**

The HopZ and HopQ loop HiBiT-tag insertions (highlighted in S1B, S1C Fig), were then tested for surface HiBiT-tag localization in intact bacteria using an extracellular detection system with luciferase reconstitution. All those loops showed strong surface exposure with HiBiT extracellular detection in intact live bacteria (several different clones tested of each insertion; S6 Fig). HopQ-Loop1 insertions had weaker reconstituted surface luciferase signals than HopQ-Loop2 and HopQ-Loop3 insertions (S6A Fig). HopQII-Loop2 insertions gave weaker signals than HopQI-Loop2 insertions, indicating lower expression of HopQII (double-HopQ-expressing strain N6). Western blot confirmed expression of the HiBiT-fused constructs (S6B Fig). Loop1 insertion reduced the overall expression of HopQI (S6B Fig). For HopZII, LoopX insertions gave relatively stronger surface luciferase signals at the surface than the LoopY tag variants (S6C Fig), while they showed slightly lower expression in Western blot (S6D Fig), indicating that LoopY insertions may be impaired in membrane insertion. Relevant fusions of HopQ and HopZ were also similarly constructed as single V5 fusions and tested (see e.g. S6E Fig). The positive tests for expression and surface localization of single and double-tagged loop insertions for HopQ, HopZ and CagN and confirmed surface localization (S6 Fig) and offered the possibility to use such loop insertions for microscopy applications at the bacterial surface.

### Generating and validating double-tagged mutants CagN-HiBiT with HopQ-V5 and HopZ-V5 (NQ & NZ)

For double tagging of both CagN and Hop proteins in *H. pylori* N6, first an expression plasmid containing *cagN* (strain 26695) with an insertion of a HiBiT tag between amino acids 214 and 215 (both duplicated) was generated (S1 Table, S2 Table) and transformed into N6 wild type strain. The transformants were checked by resistance, PCR with relevant primers targeting the plasmid, plasmid re-isolation, and by CagN-HiBiT detection in Western blot and using Nano-Glo HiBiT Extracellular Detection System (Promega #N2420) for bacterial surface detection of HiBiT-tagged proteins (S6 Fig). Subsequently, plasmids, containing V5 tags to express in exposed loops of HopQ (Loop2) and HopZ (LoopY) (S1 Fig; S2 Table) were created. The V5-inserted *hopQ* and *hopZ* genes were amplified by PCR. The mutants were shuttled into the *H. pylori* N6 chromosome of the CagN-HiBiT expression strain by natural transformation and recombination, using the MuGENT principle (77), co-transforming with a PCR of an *AphA3’-III* kanamycin resistance cassette insertion in the *rdxA* gene of *H. pylori*. The double-labelled clones were selected on kanamycin and chloramphenicol, verified by PCR sequencing, and characterized for conserved HiBiT surface detection. They were also characterized for correct HiBiT and V5 expression by Western blot (S6 Fig).

### Surface labelling of HiBiT-tagged proteins in situ (*H. pylori*)

To verify the membrane localization of the HiBiT-tagged HopQ, HopZ, CagN and CagL proteins and the surface exposure of the insertion loops in intact bacteria, the Nano-Glo® HiBiT Extracellular Detection assay (Promega #N2420) was performed with all generated HiBiT strains. This is based on the reconstitution of the small HiBiT luciferase segment by the larger LgBit segment which is added in solution. *H. pylori* strains were grown on blood agar plates for 24 h and subsequently harvested in PBS with sterile cotton swabs. In white 96-well F-bottom plates (Thermo Scientific, #236105), 50 µl of this bacterial suspension diluted to OD_600_ = 0.1 were mixed with equal amounts of the Nano-Glo® HiBiT Extracellular Buffer containing LgBiT Protein (1:1,000) and Nano Glo® HiBiT Extracellular Substrate (1:50). Plates were incubated for 10 min at room temperature on an orbital shaker at 750 rpm. Luminescence was measured in a Victor Nivo Multimode microplate reader (PerkinElmer). All samples were measured in triplicates. Bacteria without the tags were used as negative controls for background luminescence. Bacteria were always used after one-day growth and freshly harvested, to avoid lysis or non-specific exposure of HiBiT label on the bacterial surface. Washing intact bacteria with PBS, which was performed as an additional non-specific control to remove any potential non-specific HiBiT label with possible surface adherence from the bacterial surface, yielded the same values as without washing. To avoid misinterpretation due to potential lysis of the bacteria, we also performed specific controls, using a strain which expresses CagN-HiBiT in combination with a knock/out exchange mutation in the CagT4SS ATPase Cagα (28), using an *aphA*-III kanamycin resistance cassette insertion, or a HiBiT fusion of the intracellular RecA protein. Those strains were directly compared with the CagN- HiBiT strain in wild type background, testing for HiBiT surface localization in intact bacteria and at the same time, for total HiBiT yield in bacterial lysates, generated by ultrasonication. The ratio between total HiBiT content (measured upon ultrasonication) and surface label was calculated for the test conditions, and yielded an up to 20-fold higher ratio of total versus surface label for the control HiBiT fusion strains, verifying surface detection of CagN in wild type background.

### **Expression and purification of proteins in** *E. coli*

For recombinant protein expression and subsequent purification, the CagL gene was amplified from *H. pylori* 26695 without its predicted N-terminal signal sequence (amino acids 1-20). CagA was amplified from *H. pylori* 26695 as a truncated protein (CagA[aa1-892]), with omission of the 294 C-terminal amino acids. *hopQI* (sequence from *H. pylori* 26695), and *hopQII* (*H. pylori* K26A) were amplified as a truncated gene to code for a partial protein (HopQI[aa128-424], HopQII_K26A[aa132-412]), lacking the transmembrane bound β-barrel structure as well as hydrophobic C- and N-terminal regions in order to prevent excessive insolubility in *E. coli*. The *cagL*, *cagA*, and *hopQI* genes were cloned into the expression vector pET28a(+) (EMD Biosciences, Novagen) in frame with a C-terminal 6xHis-tag, using the enzymes NcoI and XhoI, while the *hopQII* gene from strain K26A was cloned with two additional N- and C-terminal 6xHis-tags and N-terminal thrombin cleavage site using the enzymes BamHI and XhoI. *hopQII* from *H. pylori* strain PNGhigh12A was designed and amplified as the same truncated binding domain as described for HopQI, but cloned into pET28a(+) in frame with a N-terminal 6xHis-tag and a thrombin site using enzymes BamHI and XhoI. Similarly, *hopZI* (sequence from *H. pylori* SU2) and *hopZII* (sequence from *H. pylori* 26695) were amplified as gene segments to express partial proteins (HopZI[aa37-531], HopZII[aa37-501], *hopZ*-ON sequence as reference; S1 Fig) and cloned into pET28a(+) in frame with a C-terminal 6xHis-tag using digestion enzymes SacI and XhoI. CagC was cloned as an N-terminal GST fusion protein, omitting the first 24 amino acids (forming a predicted leader peptide) initially in pGEX-4T2 (Addgene/GE Healthcare), and then recloned in between SacI and XhoI sites of pET28a(+) (all constructs in S2 Table; cloning primers in S3 Table). The cloning success of all expression constructs was checked via restriction enzyme digestions as well as sequencing. CagN was expressed and purified as described in (22).

### Expression of HopQI[aa128-424]

HopQI (sequence from *H. pylori* 26695) was expressed in *E. coli* Rosetta™(DE3) pLysS (Novagen, Merck, Germany). Expression cultures were inoculated with a starting OD_600_ of 0.1 from an overnight culture in LB medium, into TB medium, and cultured further over night at 30°C, following induction with 0.1 mM IPTG at a culture OD_600_ of 0.6 to 1.0, before culture pellets were harvested by centrifugation at 9.000xg at 4°C.

### Expression of HopQII[aa133-418] from PNGhigh12A and HopQII[aa132-412] from K26A

HopQII (sequence from *H. pylori* PNGhigh12A) was expressed in *E. coli* Rosetta™(DE3) pLysS (Novagen, Merck, Germany) from expression cultures, inoculated with a starting OD_600_ of 0.1 (from LB medium overnight culture) in TB medium and cultured further at 30°C for 4.5 h, following induction with 0.1 mM IPTG at an OD_600_ between 0.6 and 1.0.

### Expression of CagL

CagL expression cultures in *E. coli* Rosetta™(DE3) pLysS were inoculated from an overnight preculture in LB-Medium (50 μg/ml Kanamycin (Sigma)) to a starting OD_600_ of 0.1 in fresh LB medium (50 μg/ml Kanamycin (Sigma)). Cultures grown at 37°C were induced with 0.5 mM IPTG (Sigma-Aldrich) at an OD_600_ between 0.6 and 1. Expression was performed further at 16°C in LB medium, with 180 rpm shaking overnight.

### Expression of CagA, GST-CagC, and GST

Expression cultures were inoculated with a start OD_600_ of 0.1 and induced with 0.1 mM IPTG between OD_600_ 0.6 – 1.0 after initial growth at 37°C, with 180 rpm shaking. Expression of CagA[aa1-892], GST- CagC, and GST (from empty pGEX-4T2) was further carried out for 4.5 h at 30°C and 180 rpm shaking in *E. coli* Rosetta™(DE3) pLysS.

### Expression of HopZI and HopZII

HopZI and HopZII were expressed in *E. coli* Rosetta™(DE3) pLysS (Novagen, Merck, Germany) from expression cultures, inoculated with a starting OD_600_ of 0.1 and grown at 37 °C in LB medium or TB medium respectively. Cultures were induced at an OD_600_ between 0.6 and 0.8 with 0.1 mM IPTG and then grown further at 30 °C for 4.5 h.

### Protein purification from the soluble fraction (CagL, CagA, HopQI, HopQII_K26A, and GST-CagC)

Expression culture pellets were resuspended in Lysis buffer (50 mM Tris/HCl pH 8.0, 300 mM NaCl, 2% Triton-X-100, 0.1 mM DTT) and cells were mechanically disrupted with a French press at 1.9 kBar pressure for 1 cycle (One Shot cell disruptor, Constant Systems LTD). Following subsequent centrifugation at 12.000 x g for 20 min at 4°C, the proteins were purified from the soluble fraction via a Protino Ni^2+^-NTA (Ni-NTA) affinity column (Macherey&Nagel) using an Äkta Prime Plus system (GE Healthcare). The system and column were first equilibrated in purification buffer (50 mM Tris/HCl pH 8.0, 300 mM NaCl, 0.1 mM DTT) and loaded with protein. Following an extensive wash step to remove non-specifically bound protein, an additional high-salt wash step with loading buffer including 1 M NaCl was performed (to remove any remaining nucleotide contamination), followed by two mild imidazole wash steps, by step-wise increasing the Imidazole concentration in the running buffer to 25 mM and then 50 mM. The target protein was eluted by increasing the imidazole concentration in a linear gradient from 50 mM to 500 mM in the running buffer. 1 ml elution fractions were collected and analyzed on SDS gels. High protein-containing fractions were pooled, dialyzed against purification buffer and concentrated to an appropriate protein concentration using centrifugal filter devices (Merck Millipore). Protein purity as well as concentration were assessed via SDS-PAGE (S4 Fig), loading the final protein pools in various amounts along a BSA standard with defined protein concentrations.

Purification of GST proteins (GST-CagC, GST) from the soluble fractions of expression culture pellets was performed similarly to the Ni-NTA purifications. Pellets were resuspended in GST-Lysis buffer (50 mM Tris/HCl pH 8.0, 300 mM NaCl, 2% Triton-X-100, 5 mM DTT) for mechanical cell disruption as described. Using the Äkta Prime plus system (GE Healthcare) a Protino GST/4B 1 ml FPLC column (Macherey&Nagel) was equilibrated in GST-purification buffer (50 mM Tris/HCl pH 7.5, 150 mM NaCl, 5 mM DTT) before loading of the protein. Following an extensive wash step with equilibration buffer, to remove non-specifically bound protein, the target protein was eluted using a linear gradient from 0% to 100% GST elution buffer (50 mM Tris/HCl pH 8.0, 10 mM reduced glutathione, 5 mM DTT). 1 ml elution fractions were collected and analyzed as described before for Ni-NTA purified protein. An SDS gel of CagC-GST after purification is included in the supplement, S4 Fig.

### Protein purification from the insoluble fraction (HopZI, HopZII, HopQII_PNGhigh12A)

Expression culture pellets were resuspended in denaturing lysis buffer (50 mM Tris/HCl pH 8.0, 300 mM NaCl, 6 M Urea, 0.1 mM DTT) and cells mechanically disrupted with a French press at 1.9 kBar pressure for 1 cycle (One Shot cell disruptor, Constant Systems LTD). The crude cell extract was incubated at 4°C for 1 h under constant shaking on a spinning wheel prior to separation of soluble and insoluble fraction by centrifugation at 12.000 x g for 20 min at 4°C.

HopZ proteins and HopQII from PNGhigh12A were purified from the insoluble fraction via Protino Ni- NTA affinity columns (Macherey&Nagel) using an Äkta Prime plus system (GE Healthcare) equilibrated in purification buffer (50 mM Tris/HCl pH 8.0, 300 mM NaCl, 6 M Urea, 0.1 mM DTT). Following an extensive wash step with running buffer and two mild imidazole wash steps, by stepwise increasing the imidazole concentration in the running buffer to 25 mM and then 50 mM, the target protein was eluted by increasing the imidazole concentration in a linear gradient from 50 mM to 500 mM in the running buffer. 1 ml elution fractions were collected and analyzed via SDS-Page. High protein- containing fractions were pooled and step-wise dialyzed against purification buffer with decreasing urea concentrations using centrifugal filter devices (Merck Millipore). Purified HopZI and HopZII proteins were dialyzed to final urea concentrations of 0.5 M and 2 M, respectively, which was tested before in small aliquots to still support solubilization, to avoid proteins to precipitate. For subsequent assays, proteins were diluted further (at least 1:10), to appropriate concentrations, in the respective assay buffers, immediately before start of the assay.

For HopQII_PNGhigh12A specifically, high protein containing fractions after purification were pooled and concentrated to a protein concentration of 10 mg/ml using centrifugal filter devices (Merck Millipore). The concentrated protein pool was then dropwise diluted and subsequently incubated in ice-cold refolding buffer (50 mM Tris/HCl pH 8.0, 300 mM NaCl, 210 mM urea, 0.1 mM DTT) under constant stirring for 45 min. The resulting fraction was again loaded onto a Protino Ni-NTA affinity column (Macherey&Nagel) using an Äkta Prime plus system (GE Healthcare) equilibrated in native purification buffer (50 mM Tris/HCl pH = 8.0, 300 mM NaCl, 0.1 mM DTT). Following the protein loading, a wash step with a flow rate of 0.1 ml/min was employed for 45 min for a final on-column refolding of the HopQII protein. Elution was subsequently performed by a single step increase of the imidazole concentration to 500 mM in the running buffer. High protein containing fractions were again pooled, dialyzed against native purification buffer to reduce the urea content as far as possible, and concentrated to an appropriate protein concentration using centrifugal filter devices (Merck Millipore). As a final step for all proteins, protein purity as well as concentration were determined on SDS gels (S4 Fig), loading the final protein pools in various amounts along a BSA standard with defined protein concentrations.

### Protein analysis using analytical size exclusion chromatography

Quantitative analytical size-exclusion chromatography (SEC) was performed to determine the native state of highly purified HopQI protein. SEC data were collected on a Cytiva ÄKTAmicro instrument equipped with a Superdex 200 Increase (10/300) column (Cytiva). Data were evaluated with the OmniSEC software package supplied with the instrument. A commercial molecular mass standard (BioRad Gel Filtration Standard) was run before HopQ to calibrate the SEC for mass determination with the same column and buffer. Using the peak positions of the standard, we generated a calibration curve that was then used to calculate the molecular masses of the peaks in the HopQ elution profile. For analysis of protein eluted in each SEC peak of a preparative SEC run in parallel, 2 ml fractions were collected and analyzed on SDS-PAGE (S4 Fig).

### Bacterial two-hybrid (BACTH) assay

Protein-Protein interactions were studied by generation of C- or N-terminal fusions of all proteins of interest with either one of the two (T18 or T25) domains of the *Bordetella pertussis* adenylate cyclase (Cya) enzyme (76). By co-transformation of two plasmids each carrying a protein of interest fused to one of two domains of the *Bordetella pertussis* adenylate cyclase enzyme, a protein-proteins interaction can be made visible and quantifiable. Upon close interaction of the studied proteins the Cya-Enzyme activity is restored leading to cAMP synthesis and ultimately to the transcription of the β- galactosidase enzyme in the *E. coli* BTH101 host. β-galactosidase activity as a readout for the studied protein-protein interactions was visualized by cleavage of the artificial substrate X-Gal (5-Bromo-4- chloro-3-indolyl-β-D-galactopyranoside, Sigma-Aldrich) in LB-Agar (blue coloring after cleavage) as well as quantitated by an enzyme activity assay in solution measuring cleavage of ONPG (yellow coloring, Miller units).

All studied proteins of interest were cloned in frame (leaving out canonical membrane domains and predicted leader peptides) with either one of the two domains of the Cya gene in the Vectors pUT18c or pKT25 (for C-terminal fusions) and pUT18 or pKNT25 (for N-terminal fusions). Expression constructs and primers used for cloning are listed in Tables S2 and S3, respectively. A number of BACTH expression constructs, designated in Table S2 as (LT) were generously donated by Laurent Terradot, UMR 5086, Lyon, France. Inserts in all generated constructs were verified via control digestion, selected PCR, and partial sequencing. For each BACTH assay, 10 ng plasmid DNA encoding a protein of interest with a T18 domain fusion was co-transformed into *E. coli* BTH101 with 10 ng plasmid DNA of a T25 domain-fused protein. Transformed bacteria were plated on Luria Bertani (LB) plates with 200 µg/ml ampicillin, 50 µg/ml kanamycin, 0.5 mM IPTG (Sigma-Aldrich) and 40 µg/ml X-Gal and incubated at 30°C for 48 h. Three single clones per transformation were jointly inoculated in 600 µl LB medium with 200 µg/ml ampicillin and 50 µg/ml kanamycin in 2 ml Eppendorf tubes. Cultures were grown at 30°C for 24 h at 180 rpm shaking. 50 µl per preculture were then used to inoculate the expression overlay-culture on top of 800 µl solid LB agar in individual wells of a 24-well cell culture plate (Greiner) in duplicates. Expression cultures were incubated at 30°C for 24 h and subsequently harvested by washing the bacteria from the agar with 500 µl 0.9% NaCl into Eppendorf tubes. Bacteria were pelleted by centrifugation at 5.000 x g at 4°C for 10 minutes and pellets were stored at -20°C until further use. In total, we tested 327 new plasmid combinations (negative and positive controls not counted). In the bar graph figures, for any specific positive result, only the combination with the highest outcome is shown. The full matrix of plasmid combinations is available upon request.

To analyse the expression of each fusion protein, one duplicate pellet per sample was resuspended in 100 µl 0.9% NaCl before cell lysis by sonication (4°C, 2x 45 sec at 100% Output on Level 5, Branson sonifier, Emerson, Danbury, USA). Soluble and insoluble fractions were separated by centrifugation at 4°C for 20 min at 9.000 x g. Total protein concentration was determined by bichinchoninic acid assays (BCA) before 10 µg total protein were loaded per lane on a 11.8% or 14% SDS-PAGE Gel run in Laemmli buffer (25 mA) for subsequent tank blotting on nitrocellulose membranes (BA85, Schleicher & Schuell) at constant 300 mA for 2 h in Towbin buffer. Antibodies specific for either the expressed proteins or the Cya enzyme domains T18 or T25 used in the fusion constructs were used to verify expression of both interaction partners. Expression testing was done for all combinations.

For quantification of the protein-protein interactions, ß-galactosidase activity measurements were performed from each sample as described before (76), and analyzed in triplicates (calculated in Miller units). Positive and negative controls were incorporated in each individual assay to control for potential biological or technical variation. A leucine zipper from yeast protein GCN4 with a high affinity to itself, fused to both T25 and T18 in the vectors pKT25 and pUT18 (76), respectively, was used as a positive control. The empty vector constructs pKT25 and pUT18, co-transformed, served as the negative control. Additional negative controls tested incorporated one plasmid with a gene insertion together with one empty plasmid. The detection limit for positive interaction was determined at above 1.5-fold the negative control over at least 10 experiments. Western Blots for all experiments were performed, using specific custom-generated antibodies against *H. pylori* CagN, CagM, CagI, CagH, HopZ and HopQ, or alternatively against the T25 (αT25-antiserum generously donated by Daniel Ladant at Pasteur Institute, Paris) and T18 (αT18-antibody obtained from Santa Cruz Biotechnology, mouse monoclonal antibody 3D1 against *Bordetella pertussis* Cya, #sc-13582) fusion fragments of *B. pertussis* adenylate cyclase Cya. The latter two antisera also detect the T25 and T18 fragments alone, produced by empty control plasmids. Results of BACTH experiments are provided as β-galactosidase quantification (bar graphs) and as matrix heat-maps, providing fold-change values over negative control mean (explained also in the figure legends).

### Bacterial three hybrid (BAC3H) assay

The bacterial three hybrid assay is based on the principle of the BACTH (described above) with the addition of a third protein of interest via the arabinose-inducible compatible vector pAB184a (78).

By addition of the third protein, expressed from a compatible plasmid backbone without a Cya-enzyme domain fusion, the outcome of the initial two-hybrid interactions can be altered. A reduction of ß- galactosidase signal can either be the result of a higher affinity of one domain fused protein to the third interaction partner or the formation of a three-protein complex that sterically hinders the restoration of Cya-enzyme activity. A weak two-hybrid interaction may also be stabilized by the formation of a three-protein complex without steric hindrance, thereby increasing the measured ß- galactosidase activity. For the BAC3H, similar as for the BACTH, 10 ng plasmid DNA of each plasmid were co-transformed into *E. coli* BTH101. In addition to the supplements used in the BACTH procedure, 50 µg/ml chloramphenicol was used in LB-plates for selecting clones after transformation as well as for preculture media and expression agar. A final concentration of 0.1% arabinose was used in the transformation plates as well as in the expression agar for the pAB184a vector expression induction. Protein expression control of all three potential interaction partners as well as the ß-galactosidase measurements were performed as described for the BACTH assays. Positive and negative controls were the same as described for the BACTH above, likewise with additional Western blots performed for each experiment. Additional negative controls consisted in the third plasmid pAB184a introduced as empty vector alongside the two relevant pUT18 and pKT25 clones with inserts. Results of BAC3H are provided as β-galactosidase values.

### Biolayer interferometry measurements (BLI) for quantitation and affinity of direct protein-protein interactions

For the determination of kinetic and affinity parameters of selected protein-protein interactions biolayer interferometry (BLI) measurements with the Octet RED96 System (Sartorius) were performed. Highly pure proteins (our own purified proteins for the Cag and Hop proteins, see above) and commercial preparations of hCEACAM1 (Abnova, #P6737) and α5β1-Integrin (R&D Systems, #3230- A5)) were immobilized on Amine Reactive 2nd-Generation (ARG2G) biosensors activated with 20 mM EDC, 10 mM NHS in H_2_O. Protein coupling was achieved by EDC-catalyzed amine-bond formation at a concentration of 20 mg/ml for 600 sec in 10 mM sodium acetate buffer at pH = 5.0. Sensors were subsequently inactivated in 1 M ethanolamine at pH 8.5. Analyte interactions were measured at different concentrations, in most experiments for at seven concentrations between 0.125 µM to 2 µM in 1x kinetic buffer (Sartorius), separately at both pH 7.0 or pH 6.0. Association and dissociation steps were carried out for 300 sec each. An activated reference sensor with immobilized protein but measuring a buffer blank during association and dissociation steps was used and subtracted for all kinetic measurements. All experiments were performed with 200 µl filling volume per well of a black 96-well plate (Greiner #655209) at a constant temperature of 30°C and constant shaking at 1000 rpm. All parameters were analyzed and fitted (1:1 model) and Coefficient of Determination R^2^ calculated using the ForteBio Octet red analysis software.

### Protein-protein interaction plate assays

20 ng protein were coated per well of 96-well plates (Greiner #650001) in 100 µl 1xPBS overnight at 4°C. Following a blocking step in 200 µl PBS + 10% FCS for 1 h at room temperature, the analyte protein was incubated for 2 h. For pH-dependent binding, analytes were incubated in the protein-coated blocked plates with 50 mM Tris buffer adjusted to either pH 6 or 7, with testing after the incubation period that the pH remained constant. Subsequently, primary antibodies (in most tests except for CagL, we used HIS.H8 α-His (commercial mouse monoclonal, Invitrogen)) or α-CagL antisera (rabbit polyclonal sera) were diluted 1:4,000 or 1:5,000 (α-CagL) in PBS with 2% or 5% (α-CagL) FCS and incubated for 1 h at room temperature. Secondary antibodies (peroxidase-coupled goat-α-rabbit or goat-α-mouse) were diluted 1:10,000 in PBS + 1% BSA and incubated for 1 h at room temperature. Signal was developed by incubation in 100 µl tetramethylbenzidine (TMB) substrate (Sigma Aldrich) for 30 min at room temperature in the dark. The reaction was stopped by the addition of 50 µl phosphoric acid and the signal was detected by measuring the absorbance at 450 nm in a Victor Nivo Multimode microplate reader (PerkinElmer). Three subsequent wash steps with wash buffer (1xPBS + 0.05% Tween20) were performed between all described incubation steps. All incubations were performed at room temperature on an orbital shaker operated at 180 rpm, except for the TMB substrate incubation which was kept without shaking in the dark. For all experiments, wells coated with ligand protein and antibodies but not incubated with analyte served as reference blank values (background), which were subtracted from all other values. Additional controls were performed with commercial preparation of pure BSA as unrelated analyte in solution, which showed no binding and no concentration dependence towards the coupled ligands. Furthermore, plate-bound human integrin was incubated with custom-purified His-tagged *H. pylori* CagM (22) as a Cag control, followed by the same antibody combination which was not showing binding for CagM. All samples were measured at least in triplicate experiments.

### Binding assay of purified proteins to human cells

The binding of purified proteins to human cells was performed similarly as described under protein- protein binding above, in 96 well plates. As preparation, human cells were grown in cell culture 96- well plates to confluency and fixed twice for 1 h in 2% paraformaldehyde in sodium phosphate buffer, pH = 7.0. Subsequently, the fixing agent was washed out, cells treated twice with quenching buffer (0.1% glycine in PBS) and then blocked in PBS + 1% BSA or 50 mM Tris-HCl + 1% BSA with different pH values, pH 6.0 or pH 7.0. After blocking, protein co-incubation was performed in the same blocking buffer. Detection with antibodies was conducted as described above under protein-protein binding plate assays. All binding conditions were performed at least in triplicates.

### Immunofluorescent labelling, Proximity Ligation Assay (PLA) and fluorescence microscopy

The bacterial specimens were fixed two times 1 h using 2% PFA in potassium phosphate buffer, 100 mM, pH = 7, on gelatin-coated coverslips. After a subsequent quenching step, the specimens were washed twice in PBS, blocked in blocking buffer (non-permeabilizing; PBS, 1% BSA, 2% goat serum), and then exposed to primary antibodies in blocking buffer (α-HiBiT, mouse monoclonal 30E5, Promega #N7200, 1:100 diluted; or α-V5, rabbit, Invitrogen # MA5-15253, 1:500 diluted). After an overnight staining step, the samples were washed three times and then incubated for 2 h in secondary antibodies, tagged with fluorescent dyes, diluted in blocking buffer (Molecular Probes Thermo Scientific, anti-mouse, Alexa488, 1:2.500-diluted, anti-rabbit, Alexa564, 1:2.500-diluted). Finally, the samples were washed again carefully three times, counterstained with DAPI (staining bacterial nucleic acids) in PBS for 10 min, and finally embedded in Mowiol on glass slides. Negative control bacteria not bearing any tag which we stained in parallel did not show any fluorescent label with either the anti-V5 or anti HiBiT-tag antibodies. Bacteria only bearing one tag did not show any non-specific background staining when incubated with the respective other antibody (not shown).

Proximity Ligation Assay (PLA) on similar specimens with the same dual-tag insertions prepared in parallel was performed according to the manufacturer’s instructions by DuoLink (Sigma Aldrich, DuoLink in Situ PLA Kit Red #DUO92101-1KT), using two different secondary antibodies (anti-mouse- PLUS and anti-rabbit-MINUS) in combination with the α-HiBiT and α-V5 antibodies, and the PLA labelling Red chemistry. Briefly, the specimens were incubated, after standard blocking as for immunofluorescence, with primary antibodies, washed, and then incubating with the secondary antibodies in PLA blocking buffer. The ligation between the PLUS and MINUS probes, followed by wash steps and amplification reaction with DuoLink PLA Red detection reagent, all in a humid chamber at 37°C, were performed subsequently. After additional, final wash steps, PLA specimens were also counterstained with DAPI for visualization of bacterial DNA. Negative control bacteria without one or both of the tags did not show any PLA label, confirming its specificity. Routine immunofluorescence microscopy on all specimens was performed in an Olympus IX-40 microscope at 100-fold magnification. For high-resolution imaging, samples were exposed to line scanning in a high-resolution Zeiss Thunder Imager microscope with scanning functions (Mi8 microscope, Quantum Stage, highly sensitive K8 camera, and multi-line, high-intensity fluorescence LED light source), equipped with a 63- fold magnification lens. Tif-type images were generated in the different single color channels as black and white images, alternatively overlayed from the different channels in the Thunder software, and exported as single-color or multi-color tif images.

### RNA isolation of bacterial samples, cDNA synthesis and qRT-PCR

For RNA isolation of bacterial samples, *H. pylori* strains were grown on blood agar for approximately 20 h (mid-log phase) and biomass was collected with a cotton swab and shock-frozen in liquid nitrogen and stored at -80°C until further use. For collecting bacterial RNA, bacterial pellets were disrupted quickly by homogenizing with lysing matrix B (MP Biomedicals #116540425) in an MP bead beater. Total RNA of the samples was isolated using the RNeasy Mini Kit (QIAGEN, Germany) following the manufacturer’s instructions. cDNA synthesis and qRT-PCR were performed as described before in (79), applying the respective established MIQE quality standards. All sample results were normalized to bacterial 16S rDNA standards run for each sample in parallel.

### Cultivation of human cells

The human cell lines AGS (ATCC CRL-1739, human gastric adenocarcinoma cell line) and NCI N87 (ATCC CRL-5822, human gastric carcinoma cell line, kindly provided by Michael Naumann, University of Magdeburg, Germany) were cultured in RPMI 1640 medium (20 mM Hepes and GlutaMAX stable glutamine (Gibco, Thermo Fisher Scientific) and 10 % FCS (PromoCell, Germany.) and split routinely every third day. The cell line HEK-NF-κB_luc (BPSBioscience #60650, USA, luciferase reporter cell line) was cultured in DMEM supplemented with GlutaMAX (Gibco, Thermo Fisher Scientific, USA), 50 µg/µl hygromycin B (Invivogen, USA), and 10% FCS. Hygromycin was omitted from the co-incubations with bacteria. All cell cultures were grown at 37°C in a 5% CO_2_ incubator and passaged using 0.05% buffered

Trypsin-EDTA (Gibco, Thermo Fisher Scientific, USA). For NCI N87 cells to reach polarization, they were cultured for at least 7 days in the cell culture plates.

### Transient transfection of human cells

For transient transfection of human HEK-NF-κB_luc cells, Lipofectamine 2000 transfection (Invitrogen Thermo Scientific) was applied according to the manufacturer’s instructions, in 96-well plates, using 50 ng per plasmid per well. Before transfection, cell wells were supplied with 50 µl fresh Optimem medium (Invitrogen-Gibco) supplemented with 5% FCS. Plasmids transfected were: human CEACAM1 (origene); human CEACAM3 (origene); human CEACAM5 and CEACAM6 expression plasmids, kindly provided by Wolfgang Zimmermann (see ref. 47 for details). Integrin-α5 (Addgene #54970), integrin-β1 (Addgene #54129) expression plasmids were a kind gift from Michael Davidson’s lab. All plasmids were used as endotoxin–free preparations. 20 h post-transfection, used transfection medium was removed from the wells, and cells were again supplied with 50 µl of fresh growth medium (DMEM with 10% FCS). Cell co-incubations with bacteria for luciferase reporter activation assays were initiated at 24 h post-transfection and carried out for 3 h, before starting the luciferase assay.

## Co-culture of cells with live bacteria and read-out of cell activation

For co-cultures of human cells with live bacteria, human cells were seeded in 6-well, 24-well or 96-well plates at a confluency of 60-80% one day prior to the co-cultivation in antibiotic –free cell culture medium. The medium was changed 1 h prior to the infection to fresh RPMI or DMEM + 10% FCS. *H. pylori* was harvested after ca. 20 h fresh growth from blood agar plates into the respective cell culture medium, the OD_600_ of the bacterial suspension was measured, and the multiplicities of infections (MOI) were calculated. MOI = 25 was used for routine co-cultivation experiments of AGS and NCI N87 cells in 24-well plates, and MOI = 50 was used for co-cultivation experiments of HEK-NF-κB_luc reporter cells in 96-well plates. Bacterial suspensions were added to the cells and plates were centrifuged to start and synchronize the co-incubation (300 x g, 5 min at room temperature), and then further incubated at 37°C and 5% CO_2_ for different times (indicated in figures and legends). Samples were harvested after taking off the supernatant (kept for ELISA, measuring IL-8 secretion as previously described (13) using human IL-8 ELISA Set, R&D Systems), either by scraping the cells from the bottom of the plate (for RNA isolation), or by adding luminescence substrate to the cells and medium (for NF-KB-activated luciferase quantification). Luciferase production by the reporter cells was measured according to the manufacturer’s protocol using SteadyGlo Luciferase Assay (Promega). Routinely, 50 µl of cell culture medium in in each well of the 96-well plates were provided together with the adherent cells after bacterial co-incubation (for co-incubation times, see figure legends), and mixed in-plate with an equal volume of SteadyGlo luciferase cell lysis and detection reagent. After 10 min of lysis with mixing, all wells were measured in replicates in a victor Nivo multi-well plate reader in luminescence mode.

### Hummingbird phenotype (CagA translocation by *H. pylori* CagT4SS)

The quantitation of CagA-dependent phenotype (read-out for CagA translocation) was performed by *H. pylori* co-incubation of AGS cells, followed by induction and detection of cytoskeletal elongation of cells (termed the hummingbird phenotype), similarly as previously described (13). We applied the assays for two strains, 26695 (Fig 5E) and N6 (further controls, not shown). Briefly, AGS cells were seeded in wells of a 24 well plates to medium confluency (1 x 10^5^ cells per well) in standard media. Cells were let attach and grow overnight. The next day, medium was changed and bacteria were co-incubated with the attached cells at an MOI of 50, centrifuged to synchronize the infection, and let the incubation continue for 9.5 h, while development of hummingbird phenotype was monitored visually over time. Subsequently, cells were washed once to remove non-adherent bacteria, and cells with attached bacteria were fixed in 2% PFA in 100 mM potassium phosphate buffer, pH=7, overnight. PFA was changed once and let incubate for another hour. Cells were shifted to 1xPBS and images were taken in an Olympus IX40 microscope at 20x magnification, with a size marker. Tif images were exported and imported into ImageJ software, where they were further processed. At least 100 cells per condition, counting all cells randomly in each image, were quantitated for cell length in an evaluator-blinded fashion. Cell lengths with mean were calculated for each condition and dot graphs generated from the results in GraphPad Prism. Statistics were performed by One-way ANOVA with pairwise comparisons and Kruskal-Wallis test in GraphPad.

## Supporting information

Supplemental Tables and References

## Acknowledgments

Parts of the work were financially supported by a CRC 900 consortium grant awarded by the German Research Foundation (DFG) to CJ (project no. 158989968/B6) and German Center for Infection Research (DZIF) projects no. 06.809 and no. 06.820 to CJ. Bettina Sedlmaier-Erlenfeld is gratefully acknowledged for expert technical help and support throughout this study. We thank Michael Naumann for generously providing human gastric epithelial cell line NCI-N87. We are also very grateful for additional technical support by Monia Camboni. We are grateful to the intramural graduate program “Infection Research on Human Pathogens@MvPI” at the Max von Pettenkofer Institute, LMU (MMRS), and to the iLIFE RTG program initiative at LMU for supporting FM and JB. We thank all colleagues from the Josenhans laboratory for discussions and helpful suggestions.

## Author contributions

FM: performed, analysed and designed experiments, contributing to draft writing, figure design, final reading and editing; JB: performed, analyzed and designed experiments, contributing to writing (methods), figure design, final reading and editing; AL: performed and analyzed experiments, final reading and editing; GW: designed and analyzed experiments, final reading and editing; KPH: funding, experimental analysis, final reading and editing; RTAM: analyzed experiments, funding, final reading and editing; SB: performed, analyzed and designed experiments, final reading and editing; LT: designed experiments, supervision, provided materials, final reading and editing; KT: provided and analyzed large-scale data, final reading and editing. WF: designed experiments and provided materials, final reading and editing; SS: designed experiments, performed formal analysis, supervision, funding, final reading and editing; CJ: concept of study, performed, analyzed and designed experiments, supervision, funding, figure design, drafting and writing the paper, final reading and editing.

## Supplemental information titles

**Supplemental information**

**Supplementary Information S1:** Supplementary Figures S1 to S7.

**Supplementary Information S2:** S1, S2, S3, Supplementary Tables and References.

**S1 Table.:** *H. pylori* and *E. coli* strains used and generated in this study

**S2 Table.:** plasmids (BACTH, protein mutagenesis, expression)

**S3 Table.:** Primers used for cloning, PCR and sequencing

