## Supplemental Figures for "Functional and intricate interaction network connecting *Helicobacter pylori* Cag Type 4 Secretion System surface proteins with outer membrane proteins HopQ and HopZ"

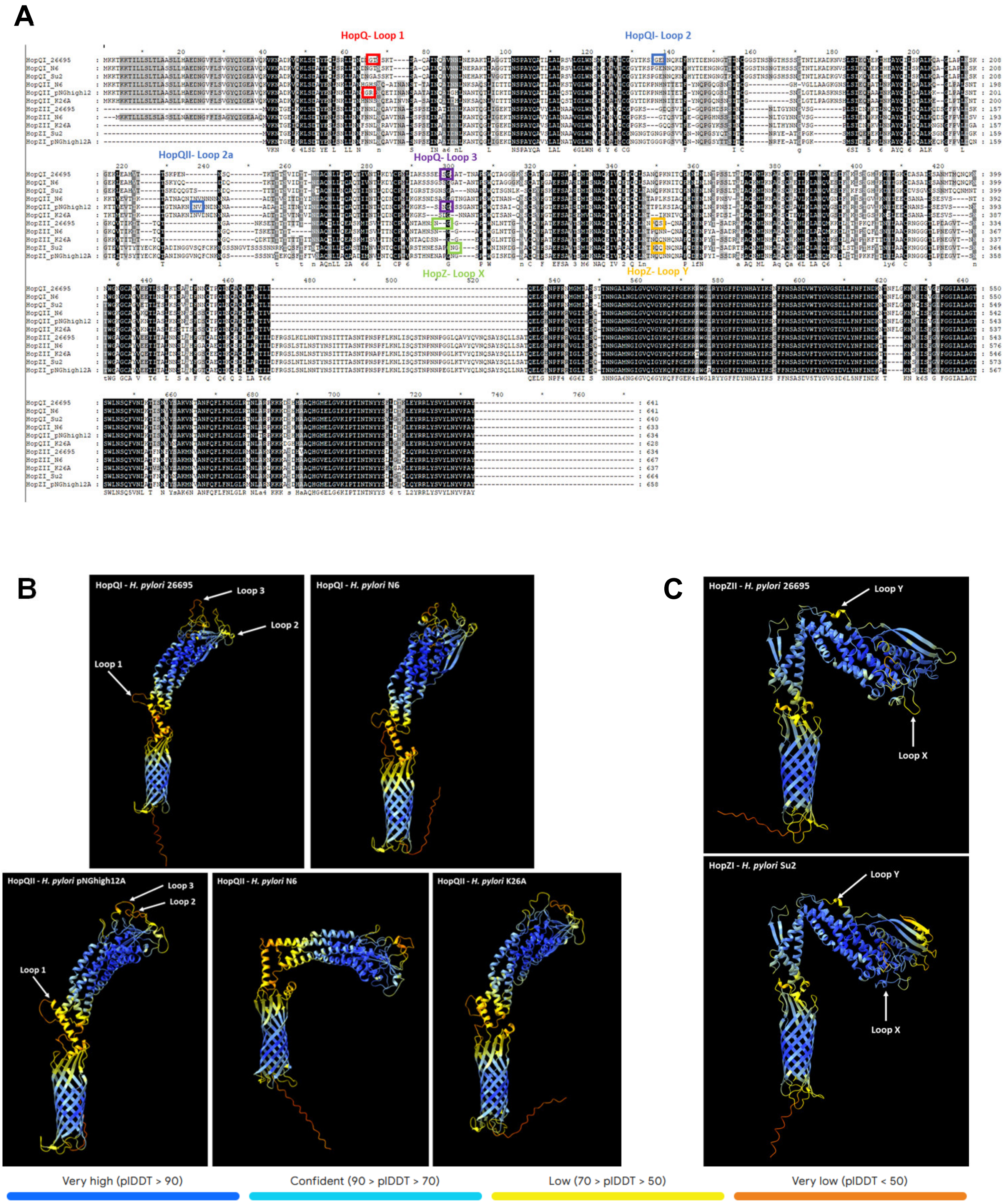


**S1 Fig, continued on next page, Figure legend see following page**

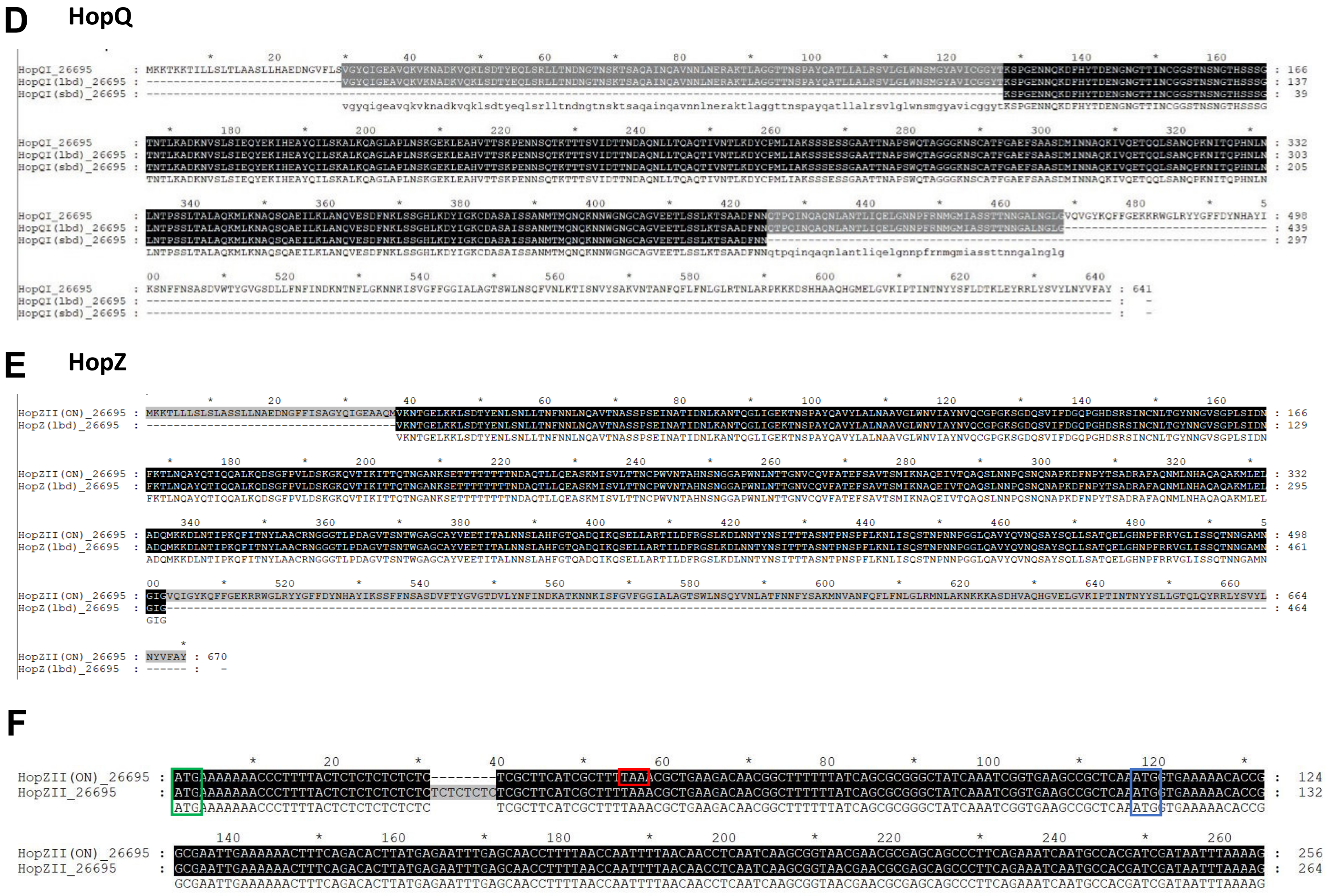


**Continued from previous page:**

**S1 Fig. Information on sequence, structure and recombinantly expressed segments of HopQ and HopZ variants.**

**A)** Amino acid alignment of HopQI, HopQII, HopZI and HopZII sequences of selected *H. pylori* strains. HopZ amino acid sequences from strains with the HopZ-OFF genotype are starting at an alternative methionine (M) start codon downstream of the CT-repeat region. The three, predicted surface-associated, loop regions (here designated as Loop 1, Loop 2 and Loop 3) utilized for HiBiT- and V5-tag insertions in HopQI are marked with colored boxes in the HopQ sequences of strains 26695 and PNGhigh12A used during this study. Two different loop 2 regions (Loop 2 and Loop 2a, respectively, were used for HopQI or HopQII variant tag insertions, as indicated. Likewise, the two loop regions (Loop X and LoopY) utilized for tag insertion in the HopZ sequences of strains 26695 and SU2 are marked.

**B)** Alphafold 3 models of selected full-length HopQI and HopQII proteins from indicated strains. All models are colored according to their predicted Local Distance Difference Test (pLDDT) on a scale of 0 to 100; color legend is given below the models. Loop regions utilized for HiBiT- and V5-tag insertions, marked in the sequence alignments above (in A), are indicated by white arrows in the respective models.

**C)** Alphafold 3 models of selected full-length HopZI and HopZII proteins from indicated strains. Amino acid sequences of HopZ-ON variants used for both models. Both models are colored according to their pLDDT on a scale of 0 to 100 with the color legend given below the models. Loop regions utilized for HiBiT- and V5-tag insertions, marked in the sequence alignment above, are indicated by white arrows in the respective models.

**D)** HopQI and **E)** HopZII amino acid alignments of the respective small-binding (sbd) and large-binding domains (lbd) used for BACTH and recombinant protein expression during this study. HopZII (ON) protein sequence of *H. pylori* strain 26695 was used in E) as a reference for alignment with recombinantly expressed HopZII-large binding domain (lbd). Beta-barrel outer membrane domains of the outer membrane proteins were omitted for expression cloning.

**F)** Gene alignment for HopZII in CT-repeat OFF- and ON-status for *H. pylori* strain 26695. Original start codon for full-length protein expression in *H. pylori* shown in green. Stop codon in open reading frame of HopZII-OFF sequence due to the CT repeat OFF switch is shown in red. Alternative potential start codon downstream of the CT-repeat region is marked in blue.

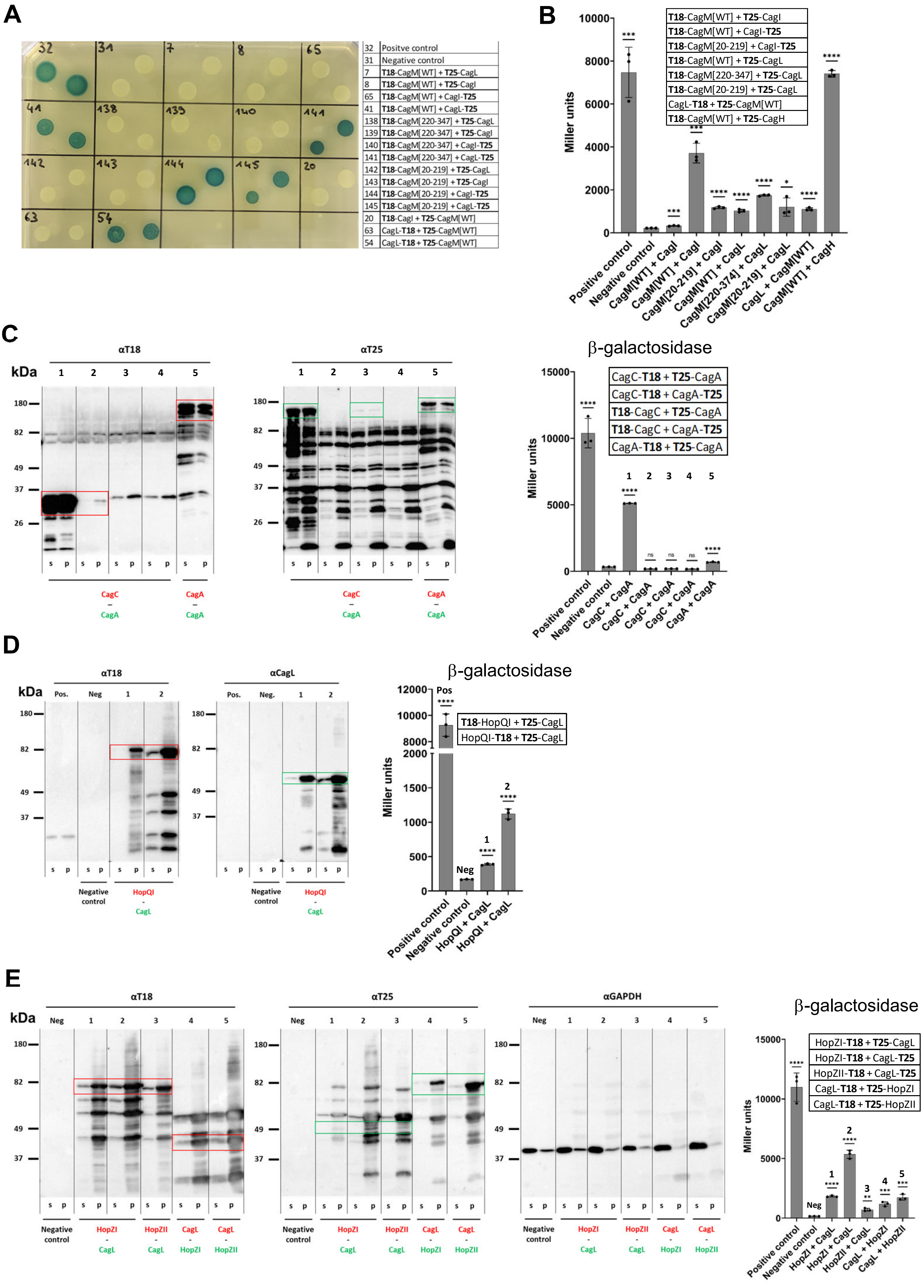


**S2 Fig, continued on next page, Figure legend see following page**

**F** Novel CagT4SS protein interactions

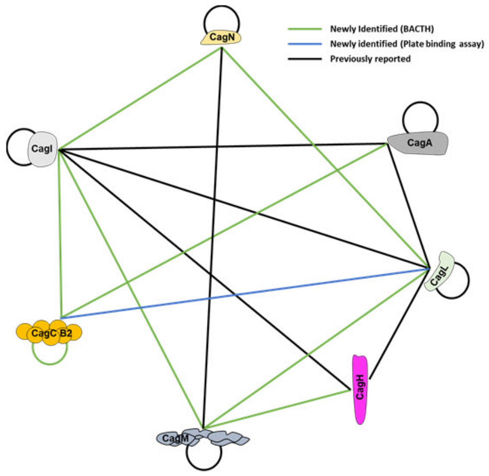


**G** Novel Hop-CagT4SS and human protein interactions

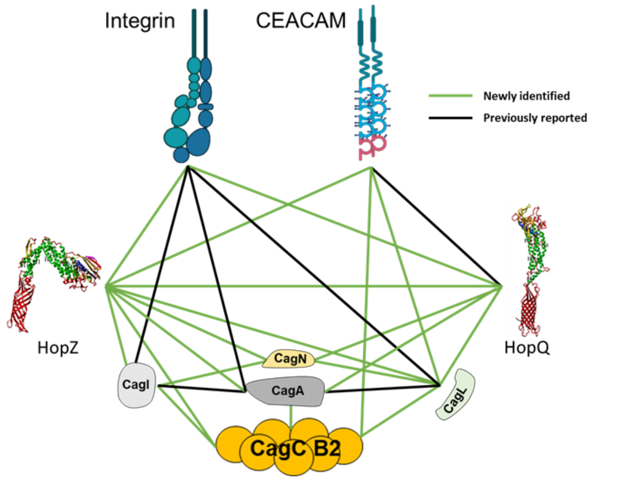


**Continued from previous page:**

**S2 Fig. [to Fig. 2] BACTH assays and controls for novel *H. pylori* CagT4SS outer protein interactions.**

**A)** Exemplary BACTH assay spotting plate demonstrating the interactions of CagM with CagL and CagI in various different expression constructs and N- and C-terminal fusions of the T18 and T25 domains. Blue color indicates above average interactions in BACTH (identity of numbered combinations see table to the right of the spotting plate).

**B)** BACTH results of β-galactosidase activity of tested interactions between CagT4SS outer proteins, depicted as bar graphs in absolute values (Miller units). All assays were performed and quantitated in three-fold replicates; negative and positive controls were run alongside each separate assay and are explained in the methods. The first listed interaction partner for each combination in the x-axis legend was expressed as an C- or N-terminal fusion in the pUT18 or pUT18c BACTH plasmids, respectively; the second listed interaction partner in each pair was expressed in pKT25 or pKNT25, respectively (see also table). CagM[WT] plasmids express full-length CagM fusions in BACTH plasmids, whereas CagM[aa20-219] and CagM[aa220-347] express fusions of either C- or N-terminal domains of CagM, respectively. Statistical analysis was performed for comparisons between each sample versus negative control (two-tailed unpaired Student's *t*-test, p-value shown above each sample. *p<0.05; ***p<0.001; ****p<0.0001. Positive samples in B are also contained in the spotting plate in A.

**C)** Immunoblot and matching β-galactosidase analysis (antibodies: αT18, mouse: 1:5.000 ; αT25, rabbit: 1:20,000) from *E. coli* cell lysates (supernatant (s) and pellet (p) fractions) depicting expression of the proteins assayed in the BACTH for interaction between CagC and CagA in various N- and C-terminal fusions of the T18 and T25 Cya-enzyme domains. Marked in red are the T18-domain fusion proteins, marked in green are the T25-domain fusion proteins. Specific bands of correct sizes are boxed. Fractionated samples 2, 3, 4 show negative interaction outcomes (see right panel), accompanied by very low or no detection of CagA, while CagC was detected (aberrant CagC size observed in 3 and 4). CagA was expressed as C-terminally truncated CagA(aa1-892), since full-length CagA had no detectable expression in any BACTH experiments. Right panel: β-galactosidase activities of BACTH samples shown in the immunoblots, as bar graphs (Miller units) in the same order as loaded on the blots. Statistical analysis was performed for comparisons between each sample versus negative control (two-tailed unpaired Student's *t*-test). ****p<0.0001; ns = non significant.

**D)** Immunoblot and matching β-galactosidase analysis from BACTH samples (antibodies: αT18, mouse: 1:5,000 ; αCagL, rabbit: 1:20,000) from *E. coli* cell lysates (supernatant (s) and pellet (p) fractions) depicting expression of the proteins assayed in the BACTH interaction between HopQI and CagL. Marked in red are the T18-domain fusion proteins, marked in green are the T25-domain fusion proteins. Specific bands of correct masses are boxed. Right panel: BACTH results of β-galactosidase activities of the samples shown in immunoblots, as bar graphs (Miller units), in the same order as on the blots. Statistical analysis was performed for comparisons between each sample versus negative control (two-tailed unpaired Student's *t*-test). ****p<0.0001.

**E)** Immunoblot and matching β-galactosidase analysis from BACTH samples (primary antibodies: αT18, mouse: 1:5,000 ; αT25, rabbit: 1:20,000) using *E. coli* cell lysates (supernatant/soluble (s) and pellet (p) fractions) depicting expression of the proteins assayed in the BACTH interaction between HopZ alleles I and II and the T4SS protein CagL in various N- and C-terminal fusions of the T18 and T25 Cya-enzyme domains. Marked in red are the T18-domain fusion proteins, marked in green are the T25-domain fusion proteins. 10 µg total protein was loaded per lane for all Western blots. Specific bands of correct sizes are boxed. Third panel from left depicts the same blot incubated with anti-GAPDH antibody as loading and fractionation control. Right panel: BACTH results of β-galactosidase activities of samples shown in immunoblots, as bar graphs (Miller units), in the same order as on the blots (numbering above blots and bar graph corresponds to the same samples). Statistical analysis was performed for comparisons between each sample versus negative control (two-tailed unpaired Student's *t*-test), and p-value is shown above each sample. ****p<0.0001; ns = non significant.

**F)** Network schematic and model of interactions between *H. pylori* CagT4SS outer proteins. Green lines are novel interactions found by BACTH, blue line indicates an interaction verified by plate-binding assay, which confirms the potential of the VirB2 (CagC) and VirB5 (CagL) orthologs of the T4SS to bind to each other (see Fig. 1). Black lines designate previously reported interactions. Coloring of lines see also line legend in F) and G.

**G)** Network model of interactions between *H. pylori* CagT4SS surface proteins, outer membrane proteins HopQ and HopZ, and the host cell receptors human CEACAM1 and α5β1-integrin (latter icons generated by BioRender).

A

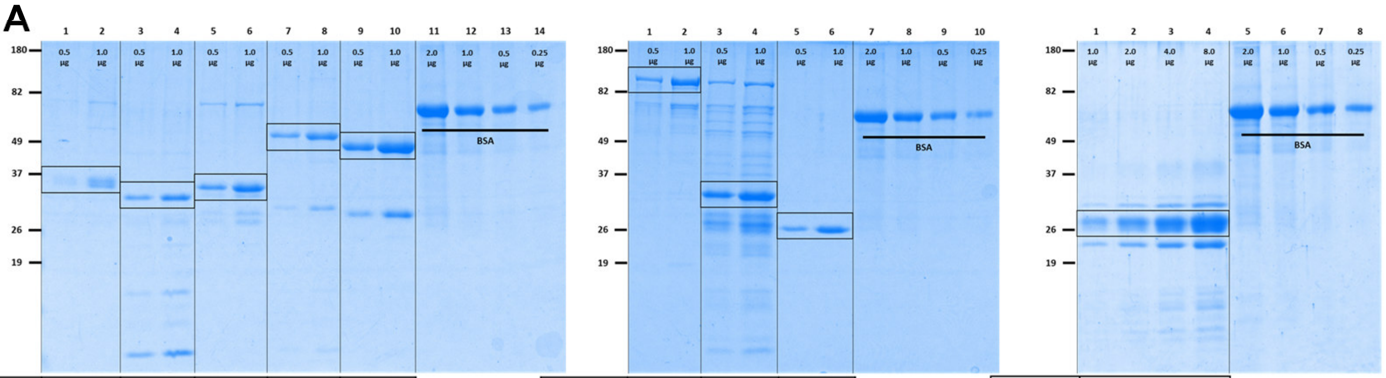


| Expression construct | HopQI [sbd] | HopQII PNGhigh12A [sbd] | HopQII K26A [sbd] | HopZI [lbd] | HopZII [lbd] |
|---|---|---|---|---|---|
| Theoretical size [kDa] | 32.7 | 33.8 | 34.5 | 53.9 | 50.9 |
| Theoretical pI | 6.45 | 8.93 | 6.55 | 7.12 | 6.59 |

| Expression construct | CagA [1-892] | GST-CagC | GST |
|---|---|---|---|
| Theoretical size [kDa] | 101.2 | 36.6 | 27.4 |
| Theoretical pI | 7.01 | 6.31 | 6.32 |

| Expression construct | CagL |
|---|---|
| Theoretical size [kDa] | 25.8 |
| Theoretical pI | 5.85 |

B

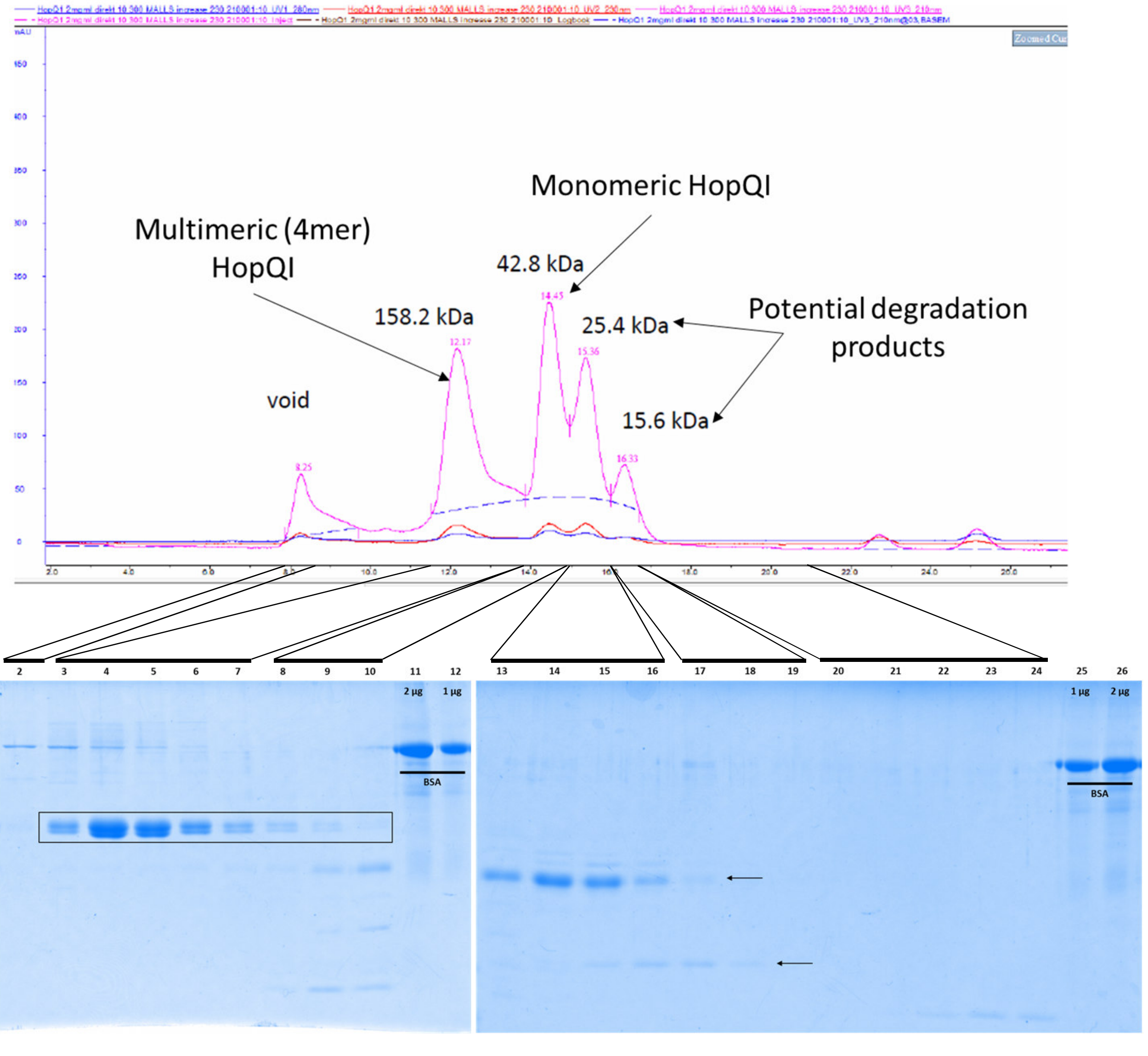


**S3 Fig. Expression, purification and characterization of recombinant purified proteins used in this study.**
**A)** SDS-Gel analysis of all recombinantly expressed and purified proteins (HopQ, HopZ, CagA, CagC, GST, CagL) used during this study. At least two different amounts per purified protein were loaded for each sample (specified on top of each lane). Defined amounts of BSA standard protein are loaded alongside the purified proteins for estimation of protein amounts.Theoretical protein mass and pI values (ProtParam, Expasy) are stated in tables below the gels.
**B)** Analytical Size Exclusion Chromatography (SEC) run of recombinant HopQI[sbd] protein showing a monomeric HopQI main peak as well as a potential multimer peak (probable tetramer). The main protein peak detector (pink curve, UV detector 3) was at 210 nm for most accurate protein detection. Red, secondary detection curve: 280 nm; blue, tertiary detection curve: 230 nm. The calculated molecular masses for each peak are given in the chromatogram. The SEC run for HopQI was repeated several times, with similar results. Lower panels: SDS gel analysis of fractions collected from SEC of recombinant HopQI[sbd] protein and assignment of the eluted fractions to the SEC peaks. Multimer peak band contains no degradation. Defined amounts of BSA are loaded for comparison. Full-length HopQ[sbd] protein in fractions is boxed. Arrows indicate HopQI degradation bands.

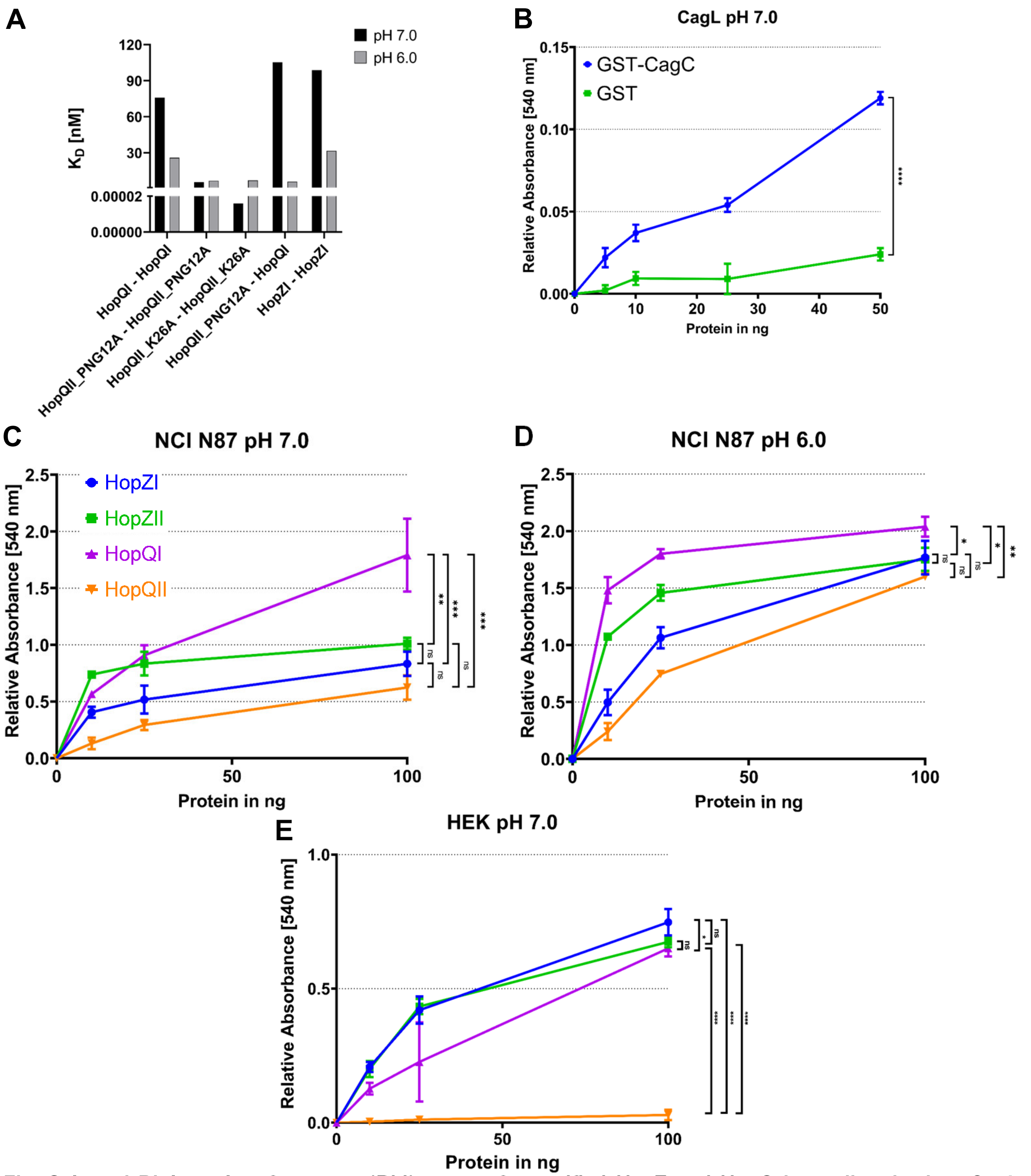


**S4 Fig. Selected Biolayer Interferometry (BLI) assays for purified HopZ and HopQ homodimerization, CagL-CagC plate binding assay and HopQ and HopZ binding to polarized NCI N87 cells.**

**A)** HopQ-HopQ and HopZ-HopZ self-interactions were assayed by biolayer interferometry (BLI) (each in six concentrations of analyte) and yielded highest affinities for HopQII-HopQII interactions, in particular for the HopQII variant from strain K26A. The assay provided medium affinities (Table 1) between HopQI-HopQI or HopZI-HopZI respectively, which were influenced by pH ($K_D$ difference between pH 6 and 7). **B)** multi-well plate binding assay of plate-immobilized purified CagL tested against soluble purified GST-CagC (GST fusion protein) in comparison to similarly purified GST moiety (negative control). Statistical differences between binding conditions shown in B) was assessed by two-tailed, unpaired Student's *t*-test. ****p<0.0001.

Panels **C)** and **D)** depict binding of all four recombinantly expressed and purified HopQ and HopZ type variants to polarized NCI-N87 cells at two different pH settings; HopQI showed the strongest binding under all conditions, while binding was slightly stronger at pH 6 compared to pH 7. HopQII (from strain PNG12_high) showed the weakest binding under all conditions. **E)** shows comparative plate binding assay of all four allelic HopQ and HopZ variants to HEK 293 wild type cells which do not express CEACAMs. Statistical difference between the compared binding assays shown in C), D), and E) respectively, was assessed by Two-way ANOVA with pairwise comparisons. *p<0.05; **p<0.01; ***p<0.001; ns = non significant.

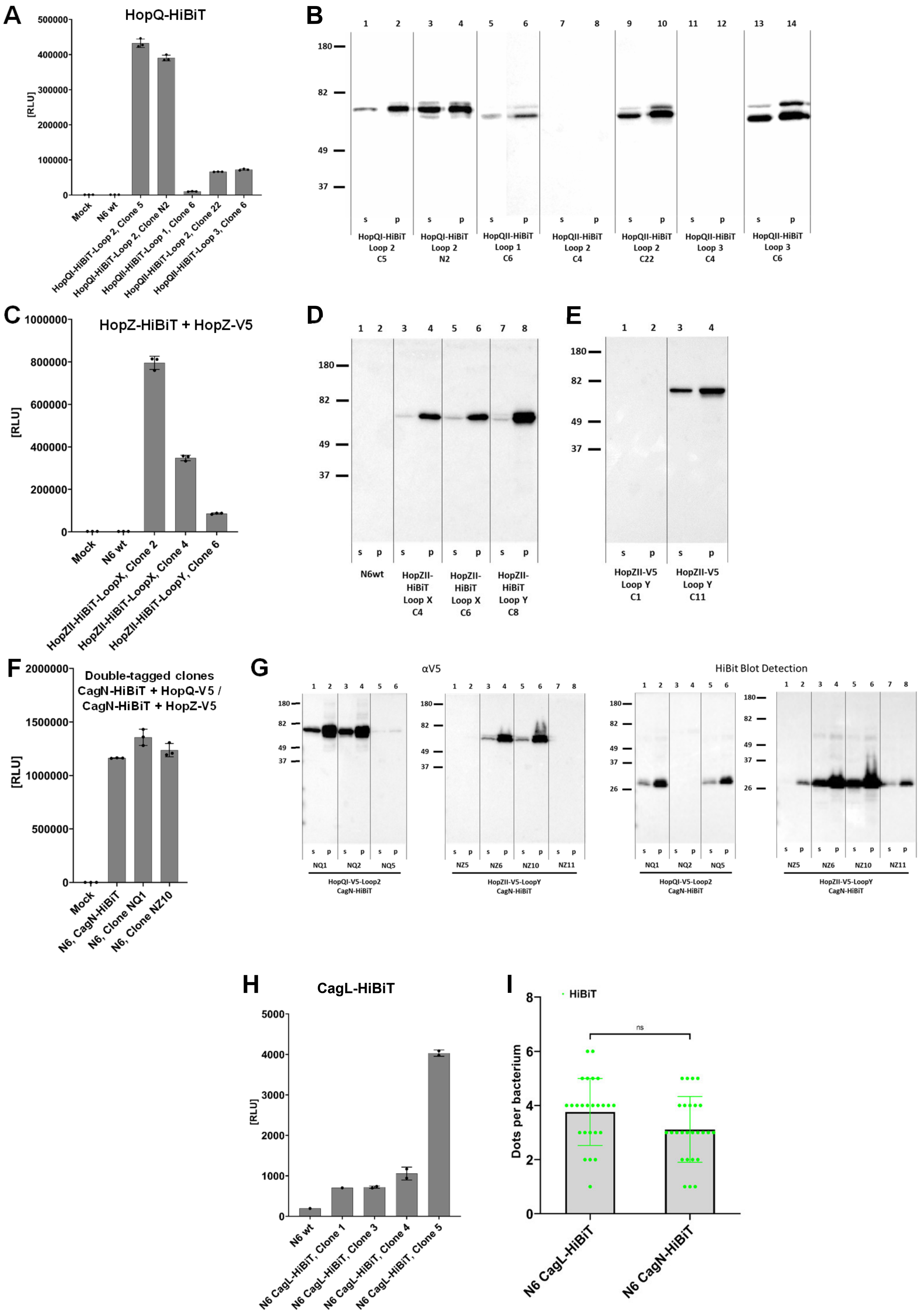


**S5 Fig, continued on following page, Figure legend see following page**

J

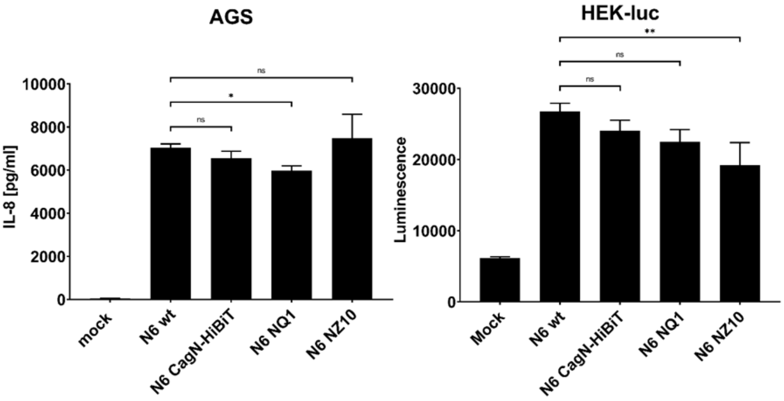


K

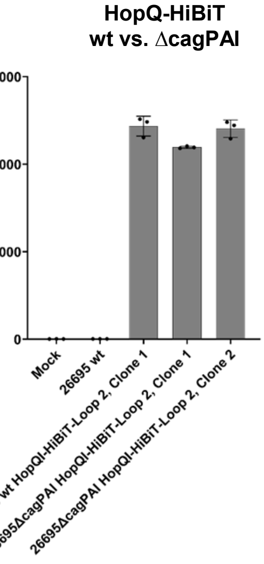


**Continued from previous page:**

**S5 Fig. Characterizing tag insertion mutants in HopQ, HopZ and CagN in strain *H. pylori* N6.**

**A)** HiBiT Extracellular Detection Assay (Promega) results of HopQ-HiBiT tag variants in *H. pylori* strain N6 alongside the N6 wild type strain for control. **B)** Immunoblot analysis with NanoGlo HiBiT Blotting system showing expression of all four generated HopQ-HiBiT variants in supernatant (s) and pellet (p) fractions obtained from seven different clones (see clone names below the lanes) from *H. pylori* N6 mutants. 10 µg total protein was loaded per lane. Some clones did not express the fusion protein, despite being genetically correct, and were not tested for surface localization. HopQI-Loop1 insertion seemed to reduce HopQ expression.

**C)** HiBiT Extracellular Detection Assay results of HopZII-HiBiT tag variants in *H. pylori* strain N6. **D)** Immunoblot analysis with NanoGlo HiBiT Blotting system showing expression of both generated HopZII-HiBiT variants in supernatant (s) and pellet (p) fractions from three different clones from *H. pylori* N6 mutants alongside the N6 wildtype strain. 10 µg total protein was loaded per lane. **E)** Immunoblot analysis with αV5 antibody (1:5,000) showing expression of both HopZII-V5-LoopY variants in supernatant (s) and pellet (p) fractions from four different clones of *H. pylori* N6 mutants. 10 µg total protein was loaded per lane.

**F)** HiBiT Extracellular Detection Assay results of generated doubly-tagged HopQI-V5-Loop2 and HopZII-V5-LoopY insertion mutants in *H. pylori* N6 CagN-HiBiT strain (NQ1: clone 1 for HopQI-V5 insertion in CagN-HiBit strain; NZ10: clone 10 for HopZII-V5 insertion in CagN-HiBiT strain) alongside the N6 CagN-HiBiT parental strain for control.

**G)** Immunoblot analysis with αV5 antibody (1:5,000) and Nano-Glo HiBiT Blotting system (right panels) showing expression of both generated double-tagged HopQI-V5-Loop2 (NQ) and HopZII-V5-LoopY (NZ) tagged, and CagN-HiBiT-tagged proteins in the *H. pylori* N6 mutant strains. Clones with single detection of V5 or HiBiT were discarded.

Panels H) to J) show visually collected microscopy counts for Cag proteins CagN and CagL detected as dots per cell.

**H)** HiBiT Extracellular Detection Assay results for HiBiT-CagL fusion protein (N-terminal HiBiT fusion, own construct not shown in detail), expressed from a chromosomal insertion into the *cagL* gene, in *H. pylori* N6. The results from four different clones are shown. Surface detection values for CagL-HiBiT clones were approximately three orders of magnitude lower than those for the CagN-HiBiT strains.

**I)** Shows microscopy dot counts of single HiBiT-tag-labelled CagN-HiBiT and HiBiT-CagL (as in panel H) bacteria (strain N6) after staining fixed bacteria with anti-HiBiT antibody (1:50-diluted) followed by goat-anti-mouse Alexa 488 (1:500).

**J)** Functional assays by co-incubating cells (AGS gastric epithelial cell line, HEK_NF-kB_luciferase reporter cell line [HEK-luc]) for 4 h or 3 h respectively, with *H. pylori* N6 V5 and HiBiT insertion strains in CagN, HopQ and HopZ, using proinflammatory cell activation as readout (IL-8 ELISA for AGS cells and luciferase measurements for HEK_luc reporter cells), those assays indicated no major loss of function of the double-tagged CagN-HiBiT insertion strains. Strain designations are listed on Y-axis of each panel. Significance of differences between strains was tested using One-way ANOVA and pairwise comparisons (shown are comparisons with regard to wt strain activity), *p<0.05, **p<0.01, ns = non significant; double insertions in CagN and HopQ or HopZ as in panels F) and G) were collected in strain N6. Those doubly-tagged mutant strains (NQ1, NZ10) were partially used in microscopy (see Fig. 5).

**K)** HiBiT extracellular detection assay results of generated HopQI-HiBiT-Loop2 mutants in *H. pylori* 26695 and *H. pylori* 26695ΔcagPAI alongside the 26695 parental strain for control. Statistics were performed using Two-way ANOVA and showed non-significant differences between HopQ surface localization in *cag*+ and *cag*- strain.

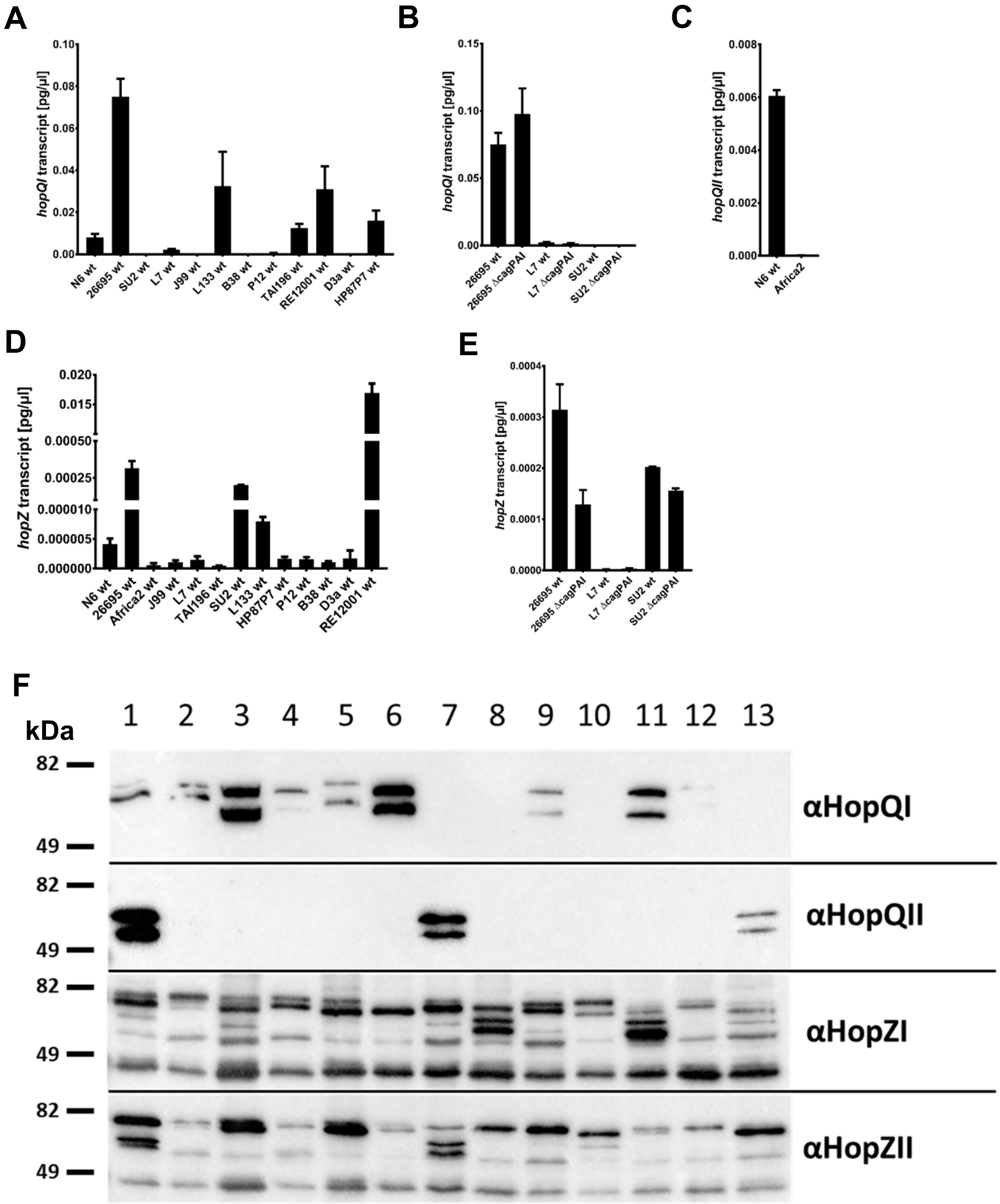


| *H. pylori* Strain | *N6* | *26695* | *Su2* | *L7* | *J99* | *L133* | *B38* | *P12* | *TAI196* | *RE12001* | *D3a* | *HP87P7* | *Africa2* |
|---|---|---|---|---|---|---|---|---|---|---|---|---|---|
| HopQ Allele | I + II | I | I | I | I | I | II | I | I | I | I | I | II |
| HopZ Allele | II | II | I | II | II | I | II | I | I | II | I | II | I |
| HopZ Status | ON | OFF | OFF | OFF | OFF | OFF | ON | ON | OFF | ON | ON | OFF | ON |

**S6 Fig. Transcripts and Western Blots of *hopQ*/HopQ and *hopZ*/HopZ: strain-specific expression.**

**A)** Quantification of absolute *hopQI* transcript amounts in different *H. pylori* wildtype strains. **B)** Quantification of absolute *hopQI* transcript amounts in three different *H. pylori* wild type strains alongside their respective Δ*cag*PAI mutants. **C)** Quantification of absolute *hopQII* transcript amounts in two different *H. pylori* wild type strains. **D)** Quantification of absolute *hopZ* (type I or type II) transcript amounts in different *H. pylori* wild type strains. **E)** Quantification of absolute *hopZ* transcript amounts in three different *H. pylori* wild type strains alongside their respective Δ*cag*PAI mutants. All shown qPCR assays were performed in technical duplicates. All values are normalized against 16S rRNA quantification and used as correction factor for each strain. Strain N6 (16S rDNA amount) was used as a reference for normalization of transcript amounts between strains.

**F)** Immunoblot analysis for protein expression of HopQ and HopZ in various *H. pylori* wild type strains. Antisera (polyclonal rabbit) used were: αHopQI, 1:2,000; αHopQII, 1:10,000; αHopZI, 1:5,000; αHopZII, 1:5,000. Tested were pellet (membrane-containing) fractions generated from different *H. pylori* wildtype strains, showing the expression of both alleles of the outer membrane proteins HopQ and HopZ. HopQ and HopZ alleles present in the genomes and *hopZ* ON- or OFF-status of each strain are given in the table below the blot.

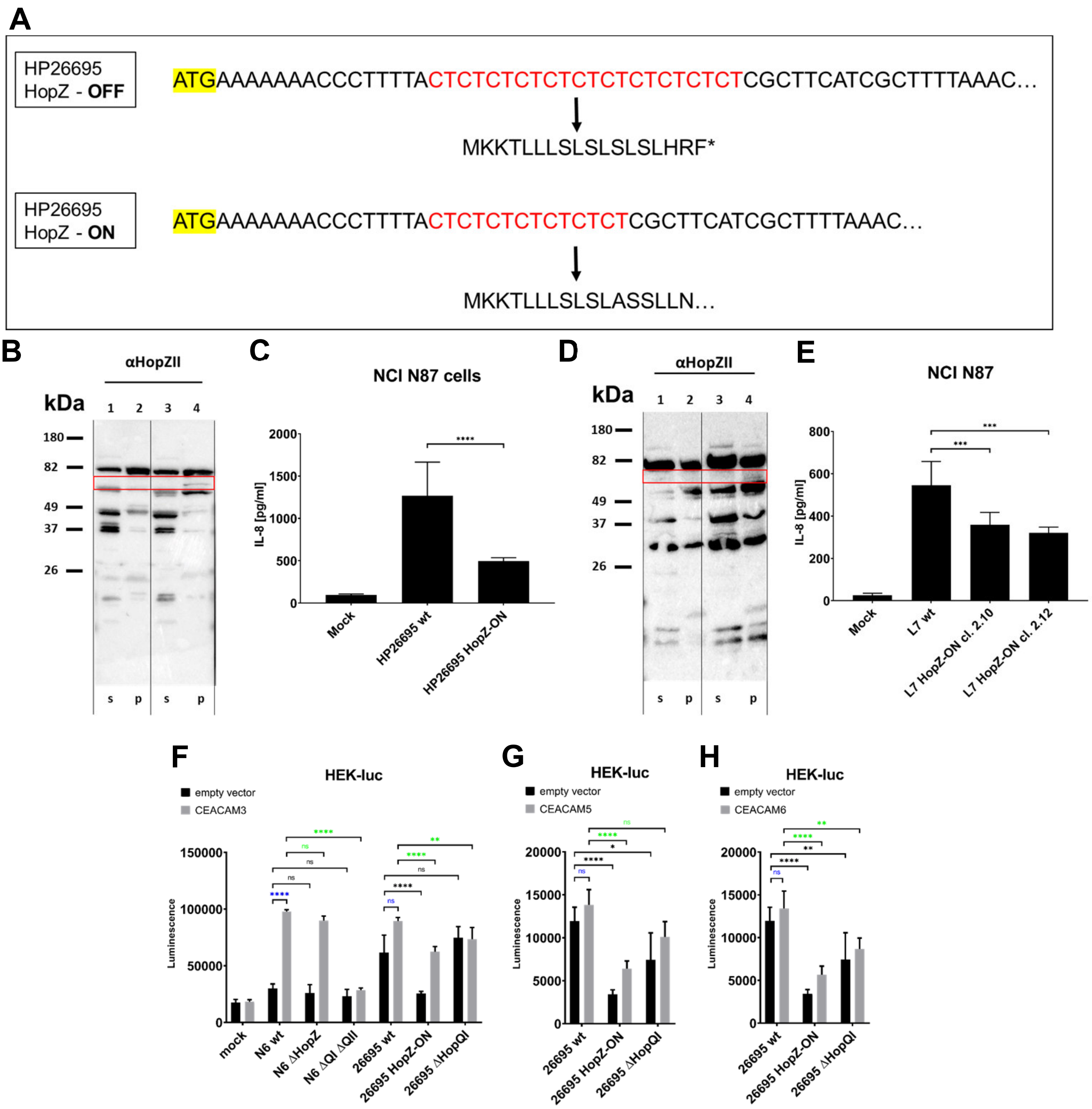


**S7 Fig. Switching ON *hopZ* in *hopZ* CT-OFF strain 26695 and detecting increased HopZ production and corresponding phenotypes in NCI-N87 and AGS gastric epithelial cell lines. Phenotypes of HopZ and HopQ mutant strains in hCEACAM3-, hCEACAM5-, hCEACAM6-transfected HEK293-luciferase reporter cells.**

**A)** Sequence of *hopZII* gene from *H. pylori* strain 26695 in its natural OFF (11 CT repeats at the beginning of the coding sequence) or engineered to be in its constitutive ON (7 CT repeats) status. **B)** Western Blot of the 26695 wild type and the CT-ON mutant shows that HopZII is expressed in the ON mutant. **C)** functional analysis of the 26695 HopZ OFF and engineered ON strain in co-incubation assays (4 h) with NCI N87 cells. IL-8 secretion, quantitated by ELISA, is significantly reduced in the ON mutant. **D)** Western Blot of the L7 wild type and its CT-ON mutant shows that HopZII is expressed in the ON mutant. Immunodetection in B) and D) was performed with custom-produced αHopZII antibody (rabbit, 1:5,000). **E)** functional analysis of the L7 HopZ-OFF wild type and engineered ON strain (two clones) in co-incubation assays (4 h) with NCI N87 cells. Statistical difference between the different strains compared in co-incubation assays in C), E), respectively, was assessed by One-way ANOVA with pairwise comparisons. ***$p<0.001$; ****$p<0.0001$; **F)**, **G)**, **H)** show functional assays (cell activation) in HEK293 NF-κB luciferase reporter cells (HEK-luc) co-incubated with *H. pylori* N6 and 26695 wild type (wt) strains and their isogenic *hopZ* and *hopQ* mutants, including the 26695 HopZ-ON strain. Cells were transfected with empty vector or with either human CEACAM3, CEACAM5, or CEACAM6 variants for 24 h, before testing them with bacterial co-incubation for 3 h. All reporter assays were performed at least three times independently, in biological triplicates. Statistical difference between the different strains compared in co-incubation assays in F), G), H), respectively, was assessed by Two-way ANOVA with pairwise comparisons. *$p<0.05$; **$p<0.01$; ***$p<0.001$; ****$p<0.0001$; ns = non significant. Black symbols indicate statistical comparisons between empty vector-transfected conditions; blue symbols indicate comparisons between empty vector-transfected and CEACAM plasmid-transfected conditions; green symbols indicate comparisons between CEACAM plasmid-transfected conditions.
