## Supplemental Tables and References for "Functional and intricate interaction network connecting *Helicobacter pylori* Cag Type 4 Secretion System surface proteins with outer membrane proteins HopQ and HopZ"

**Table S1:** *H. pylori* and *E. coli* strains used and generated in this study

| Strain name | description | reference |
|---|---|---|
| ***H. pylori* strains** | | |
| *H. pylori* N6 wt | clinical isolate (France, population hpEurope) | (1) |
| *H. pylori* 26695 wt | clinical isolate (USA, population hpEurope) | (2) |
| *H. pylori* L7 wt | clinical isolate (Japan, population hpEastAsia) | (3) |
| *H. pylori* N6 *hopZII*:Ω:*aphA3'*-III | insertion mutant of kanamycin resistance cassette in *hopZII* gene | (4) |
| *H. pylori* N6 Δ*hopQI* Ω:*aphA3'*-III (exchange mutagenesis) | insertion mutant with kanamycin resistance cassette in *hopQI* gene (pSUS3322) | This study |
| *H. pylori* N6 Δ*hopQII* | insertion mutant with chloramphenicol resistance cassette in *hopQII* gene (pCJ2365) | This study |
| *H. pylori* N6 Δ*hopQI* Δ*hopQII* | insertion mutant with kanamycin resistance cassette in *hopQI* gene (pSUS3322) and chloramphenicol resistance cassette in *hopQII* gene (pCJ2365) | This study |
| *H. pylori* 26695 *hopZII*-ON | shortening CT repeat in *hopZII* N-terminus to 7 to constitutively switch *hopZ* ORF ON (pCJ2221) | This study |
| *H. pylori* L7 *hopZII*-ON | shortening CT repeat in *hopZII* N-terminus to 7 to constitutively switch *hopZ* ORF ON (pCJ2223) | This study |
| *H. pylori* 26695 Δ*hopQI*: Ω:*aphA3*-III | insertion mutant with kanamycin resistance cassette in *hopQI* gene (pSUS3322) | This study |
| *H. pylori* N6 HopQI-V5-Loop2 (pCagN-HiBiT) (NQ1) | double tag mutant in *H. pylori* N6, with V5-Tag in HopQI-Loop2; expression plasmid pHel2-CagN-HiBiT (pCJ1871) | This study |
| *H. pylori* N6 HopZII-V5-LoopY (pCagN-HiBiT) (NZ10) | double tag mutant in *H. pylori* N6, with V5-Tag in HopZII-LoopY; expression plasmid pHel2-CagN-HiBiT (pCJ1871) | This study |
| *H. pylori* N6 *cagα*:Ω:*aphA3'*-III; (pCagN-HiBiT) | insertion mutant of kanamycin resistance cassette in *cagα* gene; expression plasmid pHel2-CagN-HiBiT (pCJ1871) | This study |
| ***E. coli* strains** | | |
| *E. coli* DH5α | used for cloning; Genotype: F- *endA1 recA1* hsdR17 Δ(l*acZYA-argF*) U169 *thi1* supE44 *gyrA96 relA1* | Thermo Fisher |
| *E. coli* MC1061 | used for cloning; Genotype:<br>F- *araD139* Δ(*ara-leu*) 7696 *galE15 galK16* Δ(*lac*)X74 *rpsL* ($Str^R$) *hsdR2* (rK + mK +) *mcrA mcrB1* | Thermo Fisher |
| *E. coli* BTH101 | used for BACTH and B3H assays; Genotype:<br>F– *cya$^-$99 araD139 galE15 galK16 rpsL1*($Str^R$) *hsdR2 mcrA1 mcrB1* | (5) |
| *E. coli* Rosetta™(DE3)pLysS | used for protein expression<br>Genotype:<br>B *ompT gal dcm lon hsdS*$_B$($rB^- mB^-$) λ(DE3 [*lacI lacUV5*-T7p07 *ind1 sam7 nin5*]) [*malB*$^+$]$_{K-12}$($λ^S$)<br>pLysSRARE[T7p20 *ileX argU thrU tyrU glyT thrT argW metT leuW proL* ori$_{p15A}$]($Cm^R$) | Novagen |

**Table S2:** plasmids (BACTH, protein mutagenesis, expression)

| Plasmid number | Vector backbone | Insert | Template | Reference |
|---|---|---|---|---|
| **BACTH constructs** | | | | |
| pCJ2300 | pUT18c | CagCΔaa1-30 | Strain 26695 | This study |
| pCJ2301 | pKT25 | CagCΔaa1-30 | Strain 26695 | This study |
| pCJ2303 | pUT18 | CagCΔaa1-30 | Strain 26695 | This study |
| pCJ2304 | pKNT25 | CagCΔaa1-30 | Strain 26695 | This study |
| pCJ2302 | pAB184a | CagL[aa21-237] | Strain 26695 | This study |
| pCJ2305 | pKNT25 | CagA[aa1-892] | Strain 26695 | This study |
| pCJ2306 | pKT25 | CagA[aa1-892] | Strain 26695 | This study |
| pCJ2308 | pAB184a | CagA[aa1-892] | Strain 26695 | This study |
| pCJ2309 | pUT18 | CagA[aa1-892] + RBS | Strain 26695 | This study |
| pCJ2310 | pUT18 | CagA[aa1-892] | Strain 26695 | This study |
| pCJ2311 | pAB184a | CagCΔaa1-30 | Strain 26695 | This study |
| pCJ2313 | pKT25 | HopQII[lbd] | Strain pNGhigh12A | This study |
| pCJ2314 | pUT18c | HopQII[lbd] | Strain pNGhigh12A | This study |
| pCJ2315 | pUT18 | HopQII[lbd] | Strain pNGhigh12A | This study |
| pCJ2316 | pKNT25 | HopQII[lbd] | Strain pNGhigh12A | This study |
| pCJ2320 | pUT18c | HopQI[lbd] | Strain 26695 | This study |
| pCJ2321 | pUT18 | HopQI[lbd] | Strain 26695 | This study |
| pCJ2322 | pKT25 | HopQI[lbd] | Strain 26695 | This study |
| pCJ2323 | pKNT25 | HopQI[lbd] | Strain 26695 | This study |
| pCJ2327 | pAB184a | CagNΔaa1-24 | Strain 26695 | This study |
| pCJ2328 | pAB184a | HopQI[sbd] | Strain 26695 | This study |
| pCJ2329 | pAB184a | HopQII[sbd] | Strain pNGhigh12A | This study |
| pCJ2330 | pUT18 | HopZI[lbd] | Strain Su2 | This study |
| pCJ2331 | pUT18c | HopZI[lbd] | Strain Su2 | This study |
| pCJ2332 | pKT25 | HopZI[lbd] | Strain Su2 | This study |
| pCJ2341 | pKNT25 | HopZI[lbd] | Strain Su2 | This study |

| | | | | |
|---|---|---|---|---|
| pCJ2333 | pUT18 | HopZII[lbd] | Strain 26695 | This study |
| pCJ2334 | pUT18c | HopZII[lbd] | Strain 26695 | This study |
| pCJ2335 | pKT25 | HopZII[lbd] | Strain 26695 | This study |
| pCJ2342 | pKNT25 | HopZII[lbd] | Strain 26695 | This study |
| pCJ2336 | pUT18 | hCEACAM1 | AGS cells (cDNA) | This study |
| pCJ2337 | pUT18c | hCEACAM1 | AGS cells | This study |
| pCJ2338 | pKT25 | hCEACAM1 | AGS cells | This study |
| pCJ2339 | pKNT25 | hCEACAM1 | AGS cells | This study |
| pCJ2340 | pAB184a | hCEACAM1 | AGS cells | This study |
| pCJ1825 | pUT18c | CagN[WT][aa25-306] | Strain 26695 | (6) |
| pCJ1806 | pKT25 | CagN[WT][aa25-306] | Strain 26695 | (6) |
| pCJ1853 | pKNT25 | CagN[WT][aa25-306] | Strain 26695 | (6) |
| pCJ1810 | pUT18c | CagM[WT][aa25-306] | Strain 26695 | (6) |
| pCJ1823 | pUT18c | CagM[aa220-374] | Strain 26695 | (6) |
| pCJ1824 | pUT18c | CagM[aa20-219] | Strain 26695 | (6) |
| pCJ1826 | pKT25 | CagM[WT] | Strain 26695 | (6) |
| pCJ1820 | pUT18c | CagA[WT] | Strain 26695 | This study |
| pCJ1812 | pUT18c | CagA[aa1-892] | Strain 26695 | This study |
| pCJ1852 | pKT25 | CagL[aa21-237] | Strain 26695 | (6) |
| pCJ1861 | pKT25 | CagC[Δaa1-44] | Strain 26695 | This study |
| pUT18c_CagI(LT) | pUT18c | CagI[aa21-381] | Strain 26695 | This study |
| pUT18_CagI(LT) | pUT18 | CagI[aa21-381] | Strain 26695 | This study |
| pUT18_CagH(LT) | pUT18 | CagH[WT] | Strain 26695 | This study |
| pUT18_CagL(LT) | pUT18 | CagL[aa21-237] | Strain 26695 | This study |
| pKT25_CagI(LT) | pKT25 | CagI[aa21-381] | Strain 26695 | This study |
| pKT25_CagH(LT) | pKT25 | CagH[WT] | Strain 26695 | This study |
| pKNT25_CagI(LT) | pKNT25 | CagI[aa21-381] | Strain 26695 | This study |
| pKNT25_CagL(LT) | pKNT25 | CagL[aa21-237] | Strain 26695 | This study |
| pUT18c | pUT18c | None | - | (5) |
| 18Z | pUT18c | Leucine zipper sequence | Yeast protein GCN4 | (5) |
| pKT25 | pKT25 | None | - | (5) |
| 25Z | pKT25 | Leucine zipper sequence | Yeast protein GCN4 | (5) |

| | | | | |
|---|---|---|---|---|
| pUT18 | pUT18 | None | - | (5) |
| pKNT25 | pKNT25 | None | - | (5) |
| **Expression or deletion/insertion constructs (for *E. coli* or *H. pylori*)** | | | | |
| pET28a | pET28a(+) | None | - | Novagen |
| pGEX-4T2 | pGEX-4T2 | None | - | GE Healthcare/ Addgene |
| pCJ2312 | pET28a(+) | CagA[aa1-892] | Strain 26695 | This study |
| pCJ2318 | pET28a(+) | GST-CagC | Strain 26695 | This study |
| pCJ909 | pGEX-4T2 | GST-CagCΔaa1-29 | Strain 26695 | This study |
| pCJ2324 | pET28a(+) | CagL[aa21-237] | Strain 26695 | This study |
| pCJ2325 | pET28a(+) | HopQI[sbd] | Strain 26695 | This study |
| pCJ2366 | pET28a(+) | HopQII[sbd] | Strain K26A | This study |
| pCJ2209 | pET28a(+) | HopQII[sbd] | Strain pNGhigh12A | This study |
| pCJ971 | pET28a(+) | CagN[aa25-306] | Strain 26695 | (6) |
| pCJ974 | pET28a(+) | CagN[aa25-216] | Strain 26695 | (6) |
| pCJ975 | pET28a(+) | CagN[aa25-306; C174, 187, 240, 243A] | Strain 26695 | (6) |
| pCJ1871 | pHel2 (expression in *H. pylori*) | CagN-HiBiT insertion ($CagN^{1-215}$-HiBiT-$CagN^{214-306}$) | Strain 26695 | This study |
| pCJ2213 | pET28a(+) | HopZI[lbd] | Strain Su2 | This study |
| pCJ2215 | pET28a(+) | HopZII[lbd] | Strain 26695 | This study |
| pCJ2220 | pUT18c | 300 bp up- and downstream of start codon of *hopZII* | Strain 26695 | This study |
| pCJ2221 | pUT18c | 300 bp up- and downstream of start codon of *hopZII* from *H. pylori* 26695 with 7 CT repeats instead of 11 | Strain 26695 | This study |
| pCJ2223 | pUT18c | 300 bp up- and downstream of start codon of *hopZII* from *H. pylori* L7 with 7 CT repeats instead of 8 | Strain 26695 | This study |
| pCJ2365 | pUT18c | *hopQII* with chloramphenicol resistance cassette | Strain K26A | This study |
| pCJ2349 | pUT18c | HopQI-V5-Loop2 | Strain 26695 | This study |
| pCJ2351 | pUT18c | HopZII-V5-LoopY | Strain 26695 | This study |
| pSUS3322 | pUC19 | *hopQI* with *aphA3'-III* kanamycin resistance cassette | Strain 26695 | This study |

**Table S3:** Primers used for cloning, PCR and sequencing

| Plasmid number or target gene | Name | Sequence | Reference |
|---|---|---|---|
| pCJ2300 & pCJ2301 | HP0546_d30_BamHI_fw | TTTTGGATCCTACCAGTCCTGCAGAAGGCG | This study |
| | HP0546_EcoRI_rv | TTTTGAATTCTTAGCTAGCTCCTCCGCTCT | This study |
| pCJ2303 & pCJ2304 | HP0546_d30_BamHI_fw2 | ATAATATGGATCCAGGAGGTAGATATGACCAGTCCTGCAGAAGGCG | This study |
| | HP0546_EcoRI_rv2 | ACTGGAATTCGCGCTAGCTCCTCCGCTCTC | This study |
| pCJ2302 | HP0539_fw3 | ATAATATCTGCAGAGGAGGTAGATATGGAAGATATA ACAAGCGGTT | This study |
| | HP0539_rv3 | ATAAGAATCGGCCGTCATTTAACAATGATCTTACTTG | This study |
| pCJ2305 & pCJ2310 | HPcagA_PstI_fw2 | ATAATATCTGCAGATGACTAACGAAACTATTGATC | This study |
| pCJ2305, pCJ2309, pCJ2310 | HPcagA_SacI_rv2 | ATAAGAATGAGCTCCGTTTGAGTCCATTATTATTGTTA | This study |
| pCJ2306 | HPcagA_BamHI_fw | TTGGATCCCATGACTAACGAAACTATTGATC | This study |
| | HPcagA_KpnI_rv2 | TTGGTACCTTATTTGAGTCCATTATTATTGTTATT | This study |
| pCJ2308 & pCJ2309 | HPcagA_PstI_fw1 | ATAATATCTGCAGAGGAGGTAGATATGACTAACGAAACTATTGATC | This study |
| pCJ2308 & pCJ2311 | HPcagA_EagI_rv1 | ATAAGAATCGGCCGTTATTTGAGTCCATTATTATTGTTA | This study |
| pCJ2311 | HPcagC_SbfI_fw1 | ATAATATCCTGCAGGAGGAGGTAGATATGACCAGTCCTGCAGAAGGCG | This study |
| pCJ2313, pCJ2314, pCJ2320, pCJ2322 | HPhopQ_BamHI_fw1 | TTGGATCCCGAAGACAACGGCGTTTTTTTAAGC | This study |
| pCJ2313 & pCJ2314 | HPhopQ_KpnI_rv1 | TTGGTACCTTACCCAAGCCCGTTCATCGC | This study |
| pCJ2315, pCJ2316, pCJ2321, pCJ2323 | HPhopQ_PstI_fw1 | TATCTGCAGAGGAGGTAGATATGGAAGACAACGGCGTTTTTTTAAGC | This study |
| pCJ2315 & pCJ2316 | HPhopQ_SacI_rv1 | GAATGAGCTCCGCCCAAGCCCGTTCATCGC | This study |
| pCJ2320 & pCJ2322 | HPhopQI_KpnI_rv1 | TTGGTACCTTACCCAAGGCCATTCAAGGC | This study |
| pCJ2323 & pCJ2322 | HPhopQI_SacI_rv1 | GAATGAGCTCCGCCCAAGGCCATTCAAGGC | This study |
| pCJ2327 | HPCagN_PstI_fw1 | TATCTGCAGAGGAGGTAGATATGATCAATACAGCATTATTGCCG | This study |
| | HPCagN_EagI_rv1 | ATAAGAATCGGCCGTCATTTTTTCCCATGAGCGATGC | This study |
| pCJ2328 | HPHopQI_PstI_fw1 | TATCTGCAGAGGAGGTAGATATGAAAAGTCCAGGCGAAAACAATC | This study |
| | HPHopQI_EagI_rv1 | ATAAGAATCGGCCGTCAGTTGTTAAAATCAGCGGCAC | This study |
| pCJ2329 | HPHopQII_PstI_fw1 | TATCTGCAGAGGAGGTAGATATGGATAAACCCAATCACAACATC | This study |
| | HPHopQII_EagI_rv1 | ATAAGAATCGGCCGTCAGCTGTCAAAAGAAGCGTTGC | This study |
| pCJ2330 – pCJ2332 & pCJ2341 | HPhopZI_PstI_fw1 | TATCTGCAGAGGAGGTAGATATGGTGAAAAACACCGGCGAATTG | This study |

| | | | |
|---|---|---|---|
| | HPhopZI_SacI_rv1 | GAATGAGCTCCGGCCGATCCCGTTCATCGC | This study |
| | HPhopZI_BamHI_fw1 | TTGGATCCCGTGAAAAACACCGGCGAATTG | This study |
| | HPhopZI_KpnI_rv1 | TTGGTACCTTAGCCGATCCCGTTCATCGC | This study |
| pCJ2333 – pCJ2335 & pCJ2342 | HPhopZII_PstI_fw1 | TATCTGCAGAGGAGGTAGATATGGTGAAAAACACCGGCGAATTG | This study |
| | HPhopZII_SacI_rv1 | GAATGAGCTCCGGCCGATCCCATTCATCGC | This study |
| | HPhopZII_BamHI_fw1 | TTGGATCCCGTGAAAAACACCGGCGAATTG | This study |
| | HPhopZII_KpnI_rv1 | TTGGTACCTTAGCCGATCCCATTCATCGC | This study |
| pCJ2336 – pCJ2340 | CEACAM1_BamHI_fw1 | TTGGATCCCCAGCTCACTACTGAATCCATGC | This study |
| | CEACAM1_PstI_fw1 | TATCTGCAGAGGAGGTAGATATGCAGCTCACTACTGAATCCATGC | This study |
| | CEACAM1_SacI_rv1 | GAATGAGCTCCGCGGGTATACATGGAACTGTCC | This study |
| | CEACAM1_KpnI_rv1 | TTGGTACCTTACGGGTATACATGGAACTGTCC | This study |
| | CEACAM1_EagI_rv1 | ATAAGAATCGGCCGTCACGGGTATACATGGAACTGTCC | This study |
| pCJ1812 & pCJ1820 | HPcagA_BamHI_fw1 | TTGGATCCCATGACTAACGAAACTATTGATC | This study |
| | HPcagA_KpnI_rv1 | TTGGTACCTTAAGATTTTTGGAAACCACCTT | |
| | HPcagA_KpnI_rv2 | TTGGTACCTTATTTGAGTCCATTATTATTGTTATT | This study |
| pCJ1861 | HP0546_PstI_fw | TTTTCTGCAGTTAAAACTTTGGTTATTCAGATCATTT | This study |
| | HP0546_BamHI_rv | TTTTGGATCCTTAGCTAGCTCCTCCGCTCT | |
| pCJ1871 | HP0538_Luc_fw1 | ATAAGGATCCGTGAGCGGCTGGCGCCTGTTTAAAAAAATTAGCAAATTTTCTAGACAACAC TTGAGTGGTTT | This study |
| | HP0538_Luc_rv1 | TATAGGATCCAAATTTGTTATTTTCTGTCTCTATTTC | This study |
| | HP0538_Luc_fw2 | ATAAGGATCCGTGAGCGGCTGGCGCCTGTTTAAAAAAATTAGCAGACAACACTTGAGTGGT TTAAAA | This study |
| | HP0538_Luc_rv2 | TATAGGATCCAGAAAATTTGTTATTTTCTGTCTCTA | This study |
| pUT18c_CagI(LT) | pUT18c-CagI_fw | GCGCTGCAGCGACAGAAGTAGTAATAACGCTTGAAC | This study |
| | pUT18c-CagI_rv | TAAGGATCCTCATTTGACAATAACTTTAGAGCTAG | This study |
| pUT18_CagI(LT) | pUT18-CagI_fw | GCGCTGCAGGATGACAGAAGTAGTAATAACGCTTGAAC | This study |
| | pUT18-CagI_rv | TAAGGATCCGCTTTGACAATAACTTTAGAGCTAG | This study |
| pUT18_CagH(LT) & pKT25_CagH(LT) | pUT18-CagH fw | GCGCTGCAGGATGGCAGGTACACAAGCTATATATG | This study |
| | pUT18-CagH rev | TAAGGATCCGCCTTCACGATTATTTTAGTTTGCAC | This study |
| pUT18_CagL(LT) | pUT18-CagL_fw | GCGCTGCAGGATGGAAGATATAACAAGCGGTTTAAAG | This study |
| | pUT18-CagL_rv | TAAGGATCCGCTTTAACAATGATCTTACTTGATTG | This study |
| pKT25_CagI(LT) | pKT25-CagI_fw | GCGCTGCAGCGACAGAAGTAGTAATAACGCTTGAAC | This study |

| | | | |
|---|---|---|---|
| | pKT25-CagI_rv | TAAGGATCCTCATTTGACAATAACTTTAGAGCTAG | This study |
| pKNT25_CagI(LT) | pKNT25-CagI_fw | GCGCTGCAGGATGACAGAAGTAGTAATAACGCTTGAAC | This study |
| | pKNT25-CagI_rv | TAAGGATCCGCTTTGACAATAACTTTAGAGCTAG | This study |
| pKT25_CagL(LT) | pKNT25-CagL_fw | GCGCTGCAGGATGGAAGATATAACAAGCGGTTTAAAG | This study |
| | pKNT25-CagL_rv | TAAGGATCCGCTTTAACAATGATCTTACTTGATTG | This study |
| pCJ2312 | HPcagA_NcoI_fw1 | TTCCATGGATGACTAACGAAACTATTGATC | This study |
| | HPcagA_XhoI_rv2 | GAATCTCGAGTTTGAGTCCATTATTATTGTTA | This study |
| pCJ2318 | GST-CagC_NcoI_fw1 | TTGAGCTCAGGAGGTAGATATGTCCCCTATACTAGGTTATTGGAAA | This study |
| | HP(GST-)CagC_XhoI_rv1 | GAATCTCGAGGCTAGCTCCTCCGCTCTC | This study |
| pCJ909 | HP0546_EcoR1F | TATGAATTCCCATGGTCACCAGTCCTGCAG | This study |
| | HP0546_Not1R | ATAAGAATGCGGCCGCTTAGCTAGCTCCTCCGCTC | This study |
| pCJ2324 | HPCagL_NcoI_fw1 | TTCCATGGATGGAAGATATAACAAGCGGTT | This study |
| | HPCagL_XhoI_rv1 | GAATCTCGAGTTTAACAATGATCTTACTTGAT | This study |
| pCJ2325 | HPHopQI_NcoI_fw1 | TTCCATGGATGAAAAGTCCAGGCGAAAACAATC | This study |
| | HPHopQI_XhoI_rv1 | GAATCTCGAGGTTGTTAAAATCAGCGGCAC | This study |
| pCJ2366 & pCJ2209 | HopQII_sbd_BamHI_fw | AAAAGGATCCGATAAACCCAATCACAACATCA | This study |
| | HopQII(sbd)_K26A_XhoI_rv1 | GAATCTCGAGGCTAGAAAAAGAAGTAGTGC | This study |
| | HopQII_sbd_XhoI_rv | AAAACTCGAGCTAGCTGTCAAAAGAAGCGTT | This study |
| pCJ2213 & pCJ2215 | HopZI(LBD)_XhoI-rv1 | GAATCTCGAGGCCGATCCCGTTCATCGC | This study |
| | HopZ(LBD)_SacI_fw1 | TTGAGCTCAGGAGGTAGATATGGTGAAAAACACCGGCGAATTG | This study |
| | HopZII(LBD)_XhoI_rv1 | GAATCTCGAGGCCGATCCCATTCATCGC | This study |
| *H. pylori rdxA* | rdxA_f1 | TTTAAATTTGAGCATGGGGCAG | (7) |
| *H. pylori rdxA* | rdxA_rv1 | TGAAAACACCCCTAAAAGAGCG | (7) |
| pCJ2220 | 26695-Z-CT-BamHI-Fw | AAAAGGATCCCAATGCCTGATATGGGTGGCA | This study |
| | 26695-Z-CT-KpnI-rv | AAAAGGTACCCGCCAAATACACCGCTTGATA | This study |
| pCJ2221 | HP26695-HopZ-7CT-Fw | CTCTCTCTCTCTCTCGCTTCATCGCTTTTA | This study |
| | HP26695-HopZ-7CT-Rv | AGAGAGAGAGAGAGTAAAAGGGTTTTTTTC | This study |
| pCJ2223 | L7-HopZ-300up-BamHI-Fw | AAAAGGATCC CAACCCCAGCAATGCCTGAT | This study |
| | L7-HopZ-300do-KpnI-Rv | AAAAGGTACC ACATTCCACAACCCTACCGC | This study |
| *H. pylori hopZ* | HopZ_qPCR_Fw | TCCACTAACCCTAATAACCCC | This study |
| *H. pylori hopZ* | HopZII_qPCR_N6_Rv | TCCCATTCATGGCACCATTGT | This study |
| *H. pylori hopQI* | HopQI-check-Fw | AGCAGTCTTACGGCTTTAGCT | This study |

| | | | |
|---|---|---|---|
| *H. pylori hopQI* | HopQI-check-Rv | CCCAATTGTTCTTTTGATTTTGCATTGTC | This study |
| *H. pylori hopQII* | HopQII-check-Fw | ACGCTCAATCTCAAGCAGCG | This study |
| *H. pylori hopQII* | HopQII-check-Rv | CGCTTTCACTTGAAGCATTCG | This study |
| *H. pylori* 26695 *hopQI*-km | hopQ_fwd_BamHI | TATGGATCCGAGTTAGCCTTTTTTGCCCCC | This study |
| | hopQ_OL_rev | TTTCACGCCCAAATCGCTCTCCACTTGATTGGC | This study |
| | hopQ_OL_fwd | GTGGAGAGCGATTTGGGCGTGAAAATCCCTACC | This study |
| | hopQ_rev_PstI | TAtTCTGCAGGGTAAAAACCGCTGTAAGCCC | This study |
| pCJ2365 | HopQII(K26A)Del_rv_SpeI | AAAA ACTAGTCGGTTTGGATAATTTGATACGC | This study |
| | HopQII(K26A)Del_fw_SpeI | AAAA ACTAGTTTTGGCACGGAGTTTAGC | This study |
| | pCAT1_SpeI | CGCACTAGTAACAGCTATGACCATGATTACG | (8) |
| | pCAT2_SpeI | CGCACTAGTGATATCGCATGCCTGCAGAG | (8) |
| pCJ2349 | HopQI_V5_Loop2_fw_SpeI | AAAAACTAGTGGTAAGCCTATCCCTAACCCTCTCCTCGGTCTCGATTCTACGGAAA0ACAAT CAAAAAGATTTCC | This study |
| | HopQI_HiBiT_Loop2_rv_SpeI | AAAAACTAGTGCCTGGACTTTTGGTATAAC | This study |
| pCJ2351 | HopZII_V5_Loop2_fw_SpeI | AAAAACTAGTGGTAAGCCTATCCCTAACCCTCTCCTCGGTCTCGATTCTACGAGCAATCAAA ACGCGCC | This study |
| | HopZII_HiBiT_Loop2_rv_SpeI | AAAAACTAGTTTGCGGGTTGTTAAGGC | This study |
| BACTH Sequencing Primer | pUT18_fw | GAGTTAGCTCACTCATTAGG | This study |
| | pUT18_rev | CGTCGTAGCGGAACTGG | This study |
| | pUT18_rev2 | TGAAAACCTCTGACACATGC | This study |
| | pUT18C_fw1 | TATGTCTTCTACGAGAACC | This study |
| | pKT25_fw1 | CGGATATCGACATGTTCG | This study |
| | pKTN25_fw | CTGGCACGACAGGTTTCC | This study |
| | pKTN25_rev | CTTGATGCCATCGAGTACG | This study |
| | pKTN25_rev2 | CAGCGCCATTCGCCATTC | This study |
| BAC3H Sequencing Primer | pAB184a_seq_rv | TTTGCAACGGAGTCGCTGACT | This study |
| | pBAD28_hp1366_1s | AGCGGGACCAAAGCCATGAC | This study |
| HiBiT and V5 Check Primer | HibiT_check-fw | GTGAGCGGCTGGCGCCT | This study |
| | HibiT_check-rev | AGGCGCCAGCCGCTCAC | This study |
| | V5_R2 | ATA**GCGGCCGC**CGTAGAATCGAGACCGAG | This study |
